# The Origin of Bladder Cancer from Mucosal Field Effects

**DOI:** 10.1101/2021.05.12.443785

**Authors:** Jolanta Bondaruk, Roman Jaksik, Ziqiao Wang, David Cogdell, Sangkyou Lee, Yujie Chen, Khanh Ngoc Dinh, Tadeusz Majewski, Li Zhang, Shaolong Cao, Hui Yao, John N. Weinstein, Neema Navai, Colin Dinney, Jianjun Gao, Dan Theodorescu, Christopher Logothetis, Charles C. Guo, Wenyi Wang, David McConkey, Peng Wei, Marek Kimmel, Bogdan Czerniak

**Author notes:** These authors contributed equally to this study (Jolanta Bondaruk, Roman Jaksik, Ziqiao Wang, David Cogdell). Correspondence to B. Czerniak.

## Abstract

We used whole-organ mapping to study loco-geographic molecular changes in evolution of human bladder cancer from mucosal field effects. The integrative multi-platform analyses based on genome-wide RNA sequencing, methylation, copy number variations, and whole exome sequencing identified over 100 dysregulated canonical pathways involving immunity, tissue differentiation and transformation as initiators of bladder carcinogenesis. Widespread dysregulation of interleukin signaling was the dominant change signifying the important role of inflammation and immunity in the incipient phases of urothelial carcinogenesis. The analyses of mutational patterns identified three types of mutations based on their geographic distribution and variant allele frequencies. The most common were low frequency subclonal mutations restricted to individual mucosal samples which were the progeny of their respective uroprogenitor cells. The two additional types of mutations were associated with clonal expansion and involved large areas of bladder mucosa. The first group referred to as **α** mutations, showed a low mutational frequency across the mucosa. The second group referred to as **β** mutations increased in their frequencies with disease progression and a large proportion of them represented mutated transcriptional regulators controlling proliferation. Time modeling revealed that bladder carcinogenesis is spanning 10-15 years and can be divided into dormant and progressive phases. The progressive phase lasted 1-2 years and was primarily driven by **β** mutations with high proliferative advantage. This is the first detailed molecular characterization of mucosal field effects initiating bladder carcinogenesis on the whole-organ scale. It provides novel insights into incipient phases of bladder carcinogenesis and biomarkers for early detection of bladder cancer as well as targets for preventive therapies.

## INTRODUCTION

Understanding the mechanisms that promote cancer initiation may facilitate the development of strategies to intercept and prevent carcinogenesis in its early phases before it evolves to an intractable, clinically aggressive, and often uncurable disease. Common epithelial cancers evolve from microscopically recognizable precursor lesions such as dysplasia or carcinoma *in situ*. These lesions in turn develop from still poorly understood incipient events in microscopically intact tissue referred to as field effects.^1^ Comprehensive understanding of these initiating mechanisms is not possible unless they are analyzed in the context of the entire organ affected by the disease. Bladder cancer is a particularly useful model for such studies as the simple anatomy of the organ permits the mapping of preneoplastic lesions and field effects in the adjacent microscopically normal mucosa across the entire organ. It originates in the epithelial lining of the urinary tract, traditionally referred to as transitional epithelium or urothelium for its functional and architectural features lie between stratified multi-layered and simple non-stratified epithelia. Because of its direct and virtually constant contact with urine, it is exposed to a wide range of metabolic products and environmental factors which are potentially carcinogenic. In fact, smoking, environmental and patient exposures to carcinogens are strong risk factors for the development of bladder cancer and it can be initiated in rodents using the nitrosamines from cigarette smoke.^2^ These factors combined with infectious agents and chronic inflammation induce molecular changes in microscopically normal appearing urothelium that can initiate carcinogenesis. Here we used a whole-organ histologic and genomic mapping (WOHGM) approach to analyze the molecular profile of bladder cancer evolution from mucosal field effects on a whole-organ scale.^3^ In WOHGM we combined the spatial microscopic assessment of the entire bladder mucosa with multi-platform genomic analyses of mRNA, methylation, copy number variation, and exome mutational profiles to visualize genetic and epigenetic changes of bladder cancer evolving from mucosal field effects through its intrinsic molecular luminal and basal tracks. This whole-organ based genomic analytical algorithm provides in depth molecular characterizations of early events initiating bladder carcinogenesis.

## RESULTS

### Preparation of Whole-Organ Cystectomy Maps

In order to molecularly characterize the evolution of bladder cancer from field effects, we collected geographically annotated mucosal samples from human cystectomy specimens of patients with bladder cancer. The samples corresponded to microscopically normal urothelium (NU), *in situ* preneoplastic conditions referred to as low and high-grade intraurothelial neoplasia (LGIN; HGIN), and urothelial carcinoma (UC). For whole-organ mapping, nine resected human bladders with invasive high-grade urothelial carcinoma were opened along the anterior wall and pinned down to a paraffin block **(Extended Data Table 1)**. A mapping grid was then applied separating the mucosal areas into 1x2cm sealed wells allowing DNA and RNA to be extracted and preserving the urothelium for microscopic examination from which the histologic map of the entire bladder mucosa was reconstructed **(Fig. 1A-G)**. We used whole transcriptome RNAseq on bulk RNA isolated from tumors to assign them to the basal or luminal molecular subtypes using basal to luminal transition scores as described previously **(Fig. 1H)**.^4^ The results revealed that six cystectomy specimens contained luminal cancers and the remaining three were of the basal subtype. We selected one bladder containing a luminal tumor and one bladder containing a basal tumor for multi-platform whole-organ based molecular profiling.

**Fig. 1.**
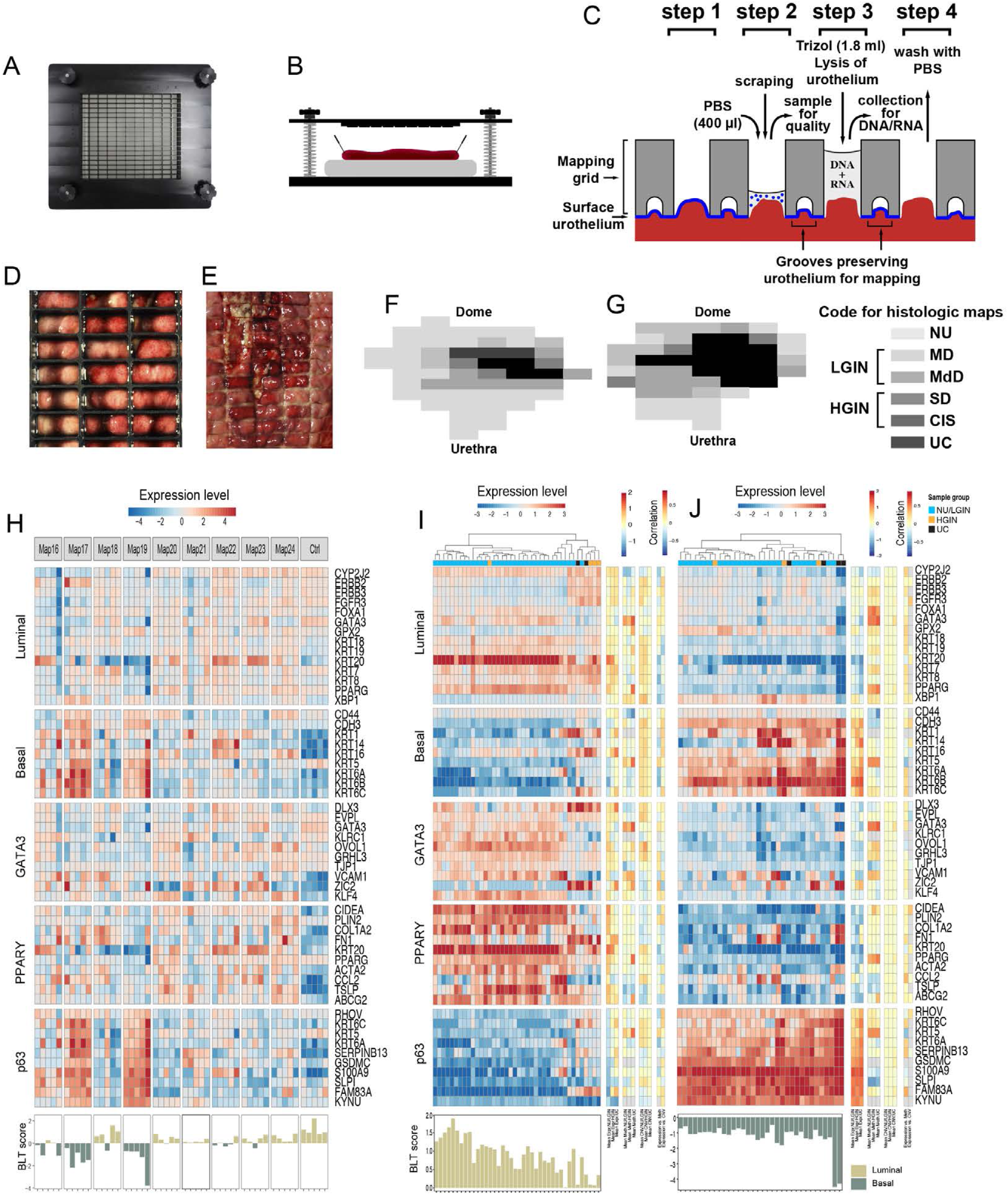
Preparation of Whole-Organ Maps. **(A)** Top view of the mapping grid. **(B)** Open cystectomy samples mounted on a paraffin block. **(C)** Diagram showing the details of the mapping grid preserving the surface urothelium for histologic mapping and facilitating simultaneous DNA/RNA extraction as well as quality assessment of cytologic preparations. The four preparation steps are described in the text. **(D)** Top view of the mapping grid superimposed over bladder mucosa. **(E)** Impression of the mapping grid on the bladder surface. **(F)** A whole-organ histologic map prepared by sampling the entire bladder mucosa of a luminal map (Map 24). **(G)** A whole-organ histologic map prepared by sampling the entire bladder mucosa of a basal map (Map 19). The histologic map code is as follows: NU, normal urothelium; MD, mild dysplasia; MdD, moderate dysplasia; SD, severe dysplasia; CIS, carcinoma *in situ*; UC, urothelial carcinoma. For analytical purposes, samples corresponding to MD and MdD were combined and referred to as low-grade intraurothelial neoplasia (LGIN). Samples corresponding to SD and CIS were combined and referred to as high-grade intraurothelial neoplasia (HGIN). **(H)** Expression analysis of selected mucosal samples of nine cystectomy specimens corresponding to NU, LGIN, HGIN, and UC. The analysis of luminal and basal markers was performed and was supplemented by the analysis of selected target genes of the luminal transcription factors (GATA3 and PPARϒ) as well as basal transcription factors (p63). The quantitative assessment of luminal and basal phenotypes was performed by the basal to luminal transition BLT score. Two cystectomy specimens identified as luminal (Map 24) and basal (Map 19) were selected for the whole-organ multi-platform genomic profiling. **(I)** Expression pattern of luminal and basal markers and BLT score in mucosal samples of cystectomy specimen with luminal cancer. **(J)** Expression pattern of luminal and basal markers and BLT score in mucosal samples of cystectomy specimen with basal cancer.

### RNA Expression

Since the origins of the basal and luminal molecular subtypes are still unresolved we used whole transcriptome expression based on RNA sequencing to characterize the changes in gene expression associated with progression. The luminal cancer developed from luminal field effects whereas the basal cancer was associated with basal field effects **(Fig. 1I, J)**. Quantitative analyses identified positive and negative BLT scores across the entire bladder mucosa in cancers developing along basal and luminal tracks respectively. These observations strongly suggest that the urothelial cancer intrinsic molecular subtypes are determined *de novo* and are consistent with animal models data implicating distinct cells of origin for the luminal and basal subtypes of bladder cancer.

We used unsupervised analyses to compare and contrast the gene expression changes that accompanied progression in the luminal and basal cystectomies. Overall, 3,040 and 2,060 genes were differentially expressed in luminal and basal cancers. In both cases hierarchical clustering using these genes separated all mucosal samples into two major groups **(Extended Data Fig. 1A, B)**. The larger cluster **α** contained nearly all NU/LGIN samples whereas nearly all samples of HGIN and UC were in cluster **β**.

We then performed supervised analyses to characterize the 3 groups of gene expression changes that accompanied the key transition points in transformation and progression. The first group (n=153 in luminal and n=146 in basal maps) contained abnormally expressed genes in samples related to NU/LGIN which retained their abnormal expression pattern with the development of HGIN and progression to UC. The second group (n=41 in luminal and n=193 in basal maps) were abnormally expressed in HGIN and UC. The third group (n=970 in luminal and n=179 in basal maps) showed an abnormal expression pattern only in UC. The top monotonically-dysregulated categories of these genes are shown in **Fig. 2A and Extended Data Fig. 2A**. The genes in the first category showed an aberrant expression pattern in early phases of bladder carcinogenesis and formed large overexpressed or downregulated plaques involving large areas of bladder mucosa. The examples of such genes mapping to chromosome 10 in the luminal map and to chromosome 11 in the basal map with their geographic relationship to field effects and precursor *in situ* lesions are shown in **Fig. 2B-D and Extended Data Fig. 2B-D**.

**Fig. 2.**
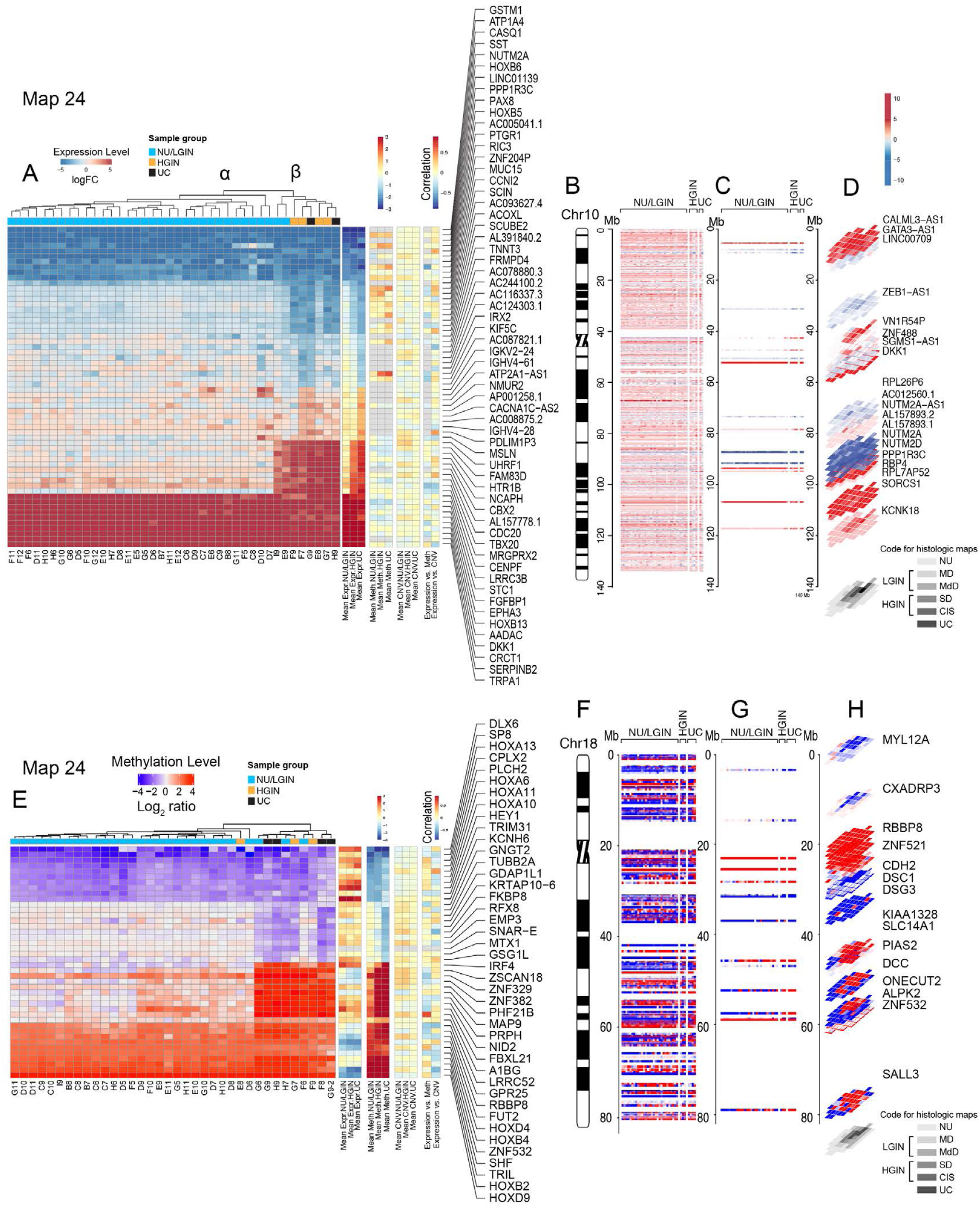
Evolution of Expression and Methylation Changes from Field Effects Through HGIN to UC in Cancer Developing Along the Luminal Track. **(A)** Hierarchical clustering using top 60 downregulated and overexpressed genes showing monotonic expression change in samples corresponding to NU/LGIN through HGIN to UC, HGIN and UC and UC only. **(B)** Whole-organ expression map of chromosome 10 showing chromosomal diagram and expression pattern of genes in individual samples of cystectomy specimen classified as NU/LGIN, HGIN, and UC. **(C)** Expression pattern of downregulated and overexpressed genes with monotonic change involving NU/LGIN, HGIN, and UC. **(D)** 3D pattern of downregulated and overexpressed genes as it relates to the whole-organ map of a cystectomy specimen shown below filtered as in panel **C**. **(E)** Hierarchical clustering using top 46 hypo- and hypermethylated genes showing monotonic expression change in samples corresponding to NU/LGIN through HGIN to UC, HGIN and UC and UC only. **(F)** Whole-organ expression map of chromosome 18 showing chromosomal diagram and methylation pattern of genes in individual samples of cystectomy specimen classified as NU/LGIN, HGIN, and UC. **(G)** Methylation pattern of hypo- and hypermethylated genes with monotonic change involving NU/LGIN, HGIN, and UC. **(H)** 3D pattern of hypo- and hypermethylated genes as it relates to the whole-organ map of a cystectomy specimen shown below filtered as in panel **E**.

In describing these three categories of genes we focused on the top 10 upregulated and downregulated genes which showed a monotonic aberrant expression pattern in samples corresponding to NU/LGIN through HGIN to carcinoma. Among the top 10 overexpressed genes in the luminal map were *DKK1,* which regulates beta-catenin-dependent Wint signaling promoting proliferation and invasion,^5^ *FGFBP1*, which may cooperate with the luminal growth factor receptor, FGFR3 to promote cell proliferation,^6^ and *HOXB13,* in which germline mutations were associated with hereditary prostate cancer.^7^ Upregulated genes in the basal map included CEACAM5/7, which was implicated in metastatic outgrowth in basal/triple negative breast cancer,^8^ SERPINB3/4, also known as squamous cell carcinoma antigen (SCCA1), which is regulated by the basal transcription factor, STAT3, and is overexpressed in the sera of patients with squamous cell carcinomas,^9^ and CXCL1, which is overexpressed in aggressive bladder cancers.^10^ Downregulated luminal genes included *GSTM1,* encoding glutathione S-transferase, genetic polymorphisms which have been linked to increased bladder cancer risk in association with smoking and exposure to environmental contaminants.^11^ The downregulated basal genes included the UPK1/2 encoding uroplakins, major components of asymmetric unit which forms the apical plaques of the umbrella cells and are markers of terminal urothelial differentiation. Their downregulation in the field effect signifies the aberrant differentiation program in bladder mucosa setting up the organ for basal cancer development. Similarly, downregulation of MUC15 and CRTAC1, both encoding extracellular glycoprotein complexes reflecting urothelial differentiation, were also observed in the basal field effect.^12, 13^

In our validation studies we compared the expression levels of the top 10 upregulated and downregulated genes identified in the field effects of the two maps with those in the TCGA cohort. The expression levels of these genes were analyzed in relation to the molecular subtypes of bladder cancer. Using the expression levels of these genes the luminal and basal subtypes can be separated into two major clusters **(Extended Data Fig. 3A-D)**. The cluster **α** in each molecular subtype was characterized by the high expression levels of these genes while cluster **β** was enriched for their downregulation. In general, the expression levels of the top 10 upregulated and downregulated genes in the early field effects were similar in both molecular subtypes. Overall, they represented frequently dysregulated genes in bladder cancer which may be used as effective early diagnostic, prognostic, and therapeutic targets **(Extended Data Fig. 3E-L)**.

Since EMT plays a major role in the development of many epithelial human cancers, including those that originate in the bladder, we assessed its role in the evolution of bladder cancer from mucosal field effects **(Extended Data Fig. 4A,B)**. In solid tumors at the core of the complex EMT circuitry are TGFB1 and p53 with their target genes positively and negativity regulating EMT. The downstream targets of this regulatory network are transcription factors that are members of the *SNAIL*, *TWIST*, *ZEB*, and *FOX* families which downregulate the expression of E-cadherin (CDH1) and other homotypic adhesion molecules including claudin-1 (CLDN1) and tight junction protein 1 (TJP1). We also have previously shown that p63 controls the expression of basal high-molecular-weight keratins (KRT5, KRT6, and KRT14) in urothelial cells.^14, 15^ The central role of p63 in the regulation of EMT was confirmed in several solid tumors.^16^ We previously showed that dysregulation of EMT played a major role in the development of basal bladder cancers and its progression to highly aggressive variants such as sarcomatoid and small cell.^17, 18^ Consistent with these observations, the activation of permissive components of EMT, such as the upregulation of TGFB1 and P53 selected target genes, was evident in field effects but widespread activation of EMT with negative EMT scores was a late event associated with progression to invasive basal cancer. In contrast, there was no major change in EMT activation with the development of luminal subtype **(Extended Data Fig. 4A, B)**.

Immune checkpoint blockade is clinically active in approximately 15% of patients with bladder cancer with the molecular subtypes characterized by distinct immune microenvironments. Therefore, we analyzed immune-related genes in the evolution of bladder cancer from field effects. The tumor developing along the luminal track was depleted from immune infiltration and its cold/null phenotype was evident *de novo* in mucosal field effects **(Extended Data Fig. 5A)**. In contrast, cancer developing along the basal track showed an increased immune signature which was already evident in mucosal field effects **(Extended Data Fig. 5B)**. The null and hot immune microenvironment of mucosal field effects in luminal and basal cancers was complemented by the downregulation and upregulation of immune regulatory genes **(Extended Data Fig. 6A, B)**. It appears that immune null luminal cancer evolved from immune cold field effects while the hot immune microenvironment of basal cancer was evident *de novo* in field effects offering early preventive targeted therapeutic opportunities.

### Methylation Changes in Progression from Field Effect to Carcinoma

We used array hybridization to characterize whole-genome methylation changes across all sections of the cystectomies. In total, 6912 and 7976 genes were found to be differentially methylated in at least one sample in the luminal (map 24) and basal (map 19) cancers when compared to normal urothelium from patients without urothelial neoplasia. As was observed with the whole transcriptome data, hierarchical clustering using the methylation levels of these genes separated mucosal samples from both maps into two major groups **(Extended Data Fig. 7A, B),** with clusters **α** containing the majority of the NU/LGIN samples and clusters **β** containing the HGIN and UC samples. We identified two groups of genes (n=1380 in the luminal map and n = 1658 in the basal map) that were abnormally methylated with the progression of neoplasia from mucosal field effects to carcinoma. The first group (n=125 in luminal and n=427 in basal maps) contained abnormally methylated genes in samples related to NU/LGIN, which retained their abnormal methylation patterns with the development of HGIN and progression to UC. The second group (n=2 in luminal and n=48 in basal maps) contained abnormally methylated genes that emerged with the development of HGIN and were retained in UC. We identified n=1253 genes in luminal and n=1183 genes in basal maps that were distinctively hypermethylated or hypomethylated in foci of carcinoma only. The top monotonically hypermethylated and hypomethylated genes in both maps are shown in **Fig. 2E and Extended Data Fig. 8A**. The genes aberrantly methylated in early phases of bladder carcinogenesis formed hypermethylated or hypomethylated plaques involving large areas of bladder mucosa; examples in the luminal and basal maps are shown in **Fig. 2F-H and Extended Data Fig. 8B-D** respectively. In describing the categories of genes abnormally methylated in the development of bladder cancer we focused on the top 10 hypermethylated and hypomethylated genes which should lead to their repression or activation, which showed monotonic methylation change in samples corresponding to NU/LGIN through HGIN to UCs. These genes were expected to be involved in the initiation of bladder carcinogenesis.

Among the top 10 hypermethylated genes in the luminal map, there were such genes as ZSCAN18 and ZNF382, transcription factors involved in proliferation, differentiation and apoptosis,^19^ MAP9 encoding microtubule-associated protein involved in mitotic progression and cell migration,^20^ NID2, a member of basement membrane proteins that binds collagens 1/4 and laminin^21^ and finally IRF4 and FBXL21, both involved in the activation of innate and adaptive immune system.^22, 23^ Among hypomethylated (activated) genes in the early field effects were DLX6 and SP8, involved in embryonal development,^24^ HEY1, helix-loop helix (bHLH)-type transcriptional repressor involved in the development of sarcomas^25^ and TRIM31, encoding an E3 ubiquitin-protein ligase, a negative regulator of cell growth.^26^ Several members of HOX gene family involved in embryonal development and body patterning were hypomethylated in field effects and included HOXA6,10,11, and 13.^27^

Among the top 10 hyper methylated genes in the basal cancer were TRH, a member of the thyrotropin-releasing hormone family that controls proliferation and inhibits apoptosis in epithelial cells, GPR25, the G-protein coupled receptor, ZNF532, Zinc Finger Protein transcription factor, and MLLT6, PHD Finger containing transcription factors involved in the regulation of several oncogenic pathways including NFκB.^28–30^ Similar to the luminal map, several members of the HOX gene family were hypermethylated in samples corresponding to NU/LGIN and included HOXB2 and 4 as well as HOXD4 and 9.^27^

Similarly to the expression analysis we compared the methylation changes in early field effects of the two maps with those in the TCGA cohort. Using the **β** value reflecting the proportion of methylated DNA for these genes, we clustered the luminal and basal subtypes of bladder cancer, which can be separated into two groups designated as **α** and **β** with enrichment of hypermethylated and hypomethylated genes, respectively **(Extended Data Fig. 9A-D)**. Dysmethylation patterns of these genes were similar in luminal and basal subtypes of bladder cancer. Overall, the top dysmethylated genes identified in early field effects were frequently hypermethylated or hypomethylated in bladder cancer and may represent significant early diagnostic, prognostic, and therapeutic targets **(Extended Data Fig. 9E-L)**.

### Mutational Heterogeneity of Field Effects and its Clonal Enrichment in Progression to Carcinoma

The analysis of whole exome sequencing on DNA from the geographically-mapped mucosal samples identified non-synonymous variant alleles in 1,379 and 2,687 genomic loci in the luminal and basal cystectomy maps, respectively **(Extended Data Table 2)**. A heat map of VAFs (variable allele frequencies) of these non-silent mutations in individual mucosal samples is shown in **Fig. 3A** and **Extended Data Fig. 10A**. They were separated into two major clusters designated as **A** and **B**. Clusters **A** contained mutations with low VAFs restricted to individual mucosal samples, whereas Clusters **B** contained mutations that were shared by multiple mucosal samples and displayed increases of VAFs with histologic progression from normal to HGIN to UC. Focusing on the latter, we analyzed VAFs in all mucosal samples that exhibited significant increases in the number of clonal mutations when the disease evolved to HGIN and carcinoma. We restricted our analytical approach to variant alleles showing alterations in at least three mucosal samples with VAFs ≥1% in at least one sample. The results yielded 157 genes with non-silent SNVs or indels in the luminal map and 198 genes in the basal map **(Fig. 3B and Extended Data Fig. 10B)**.

**Fig. 3.**
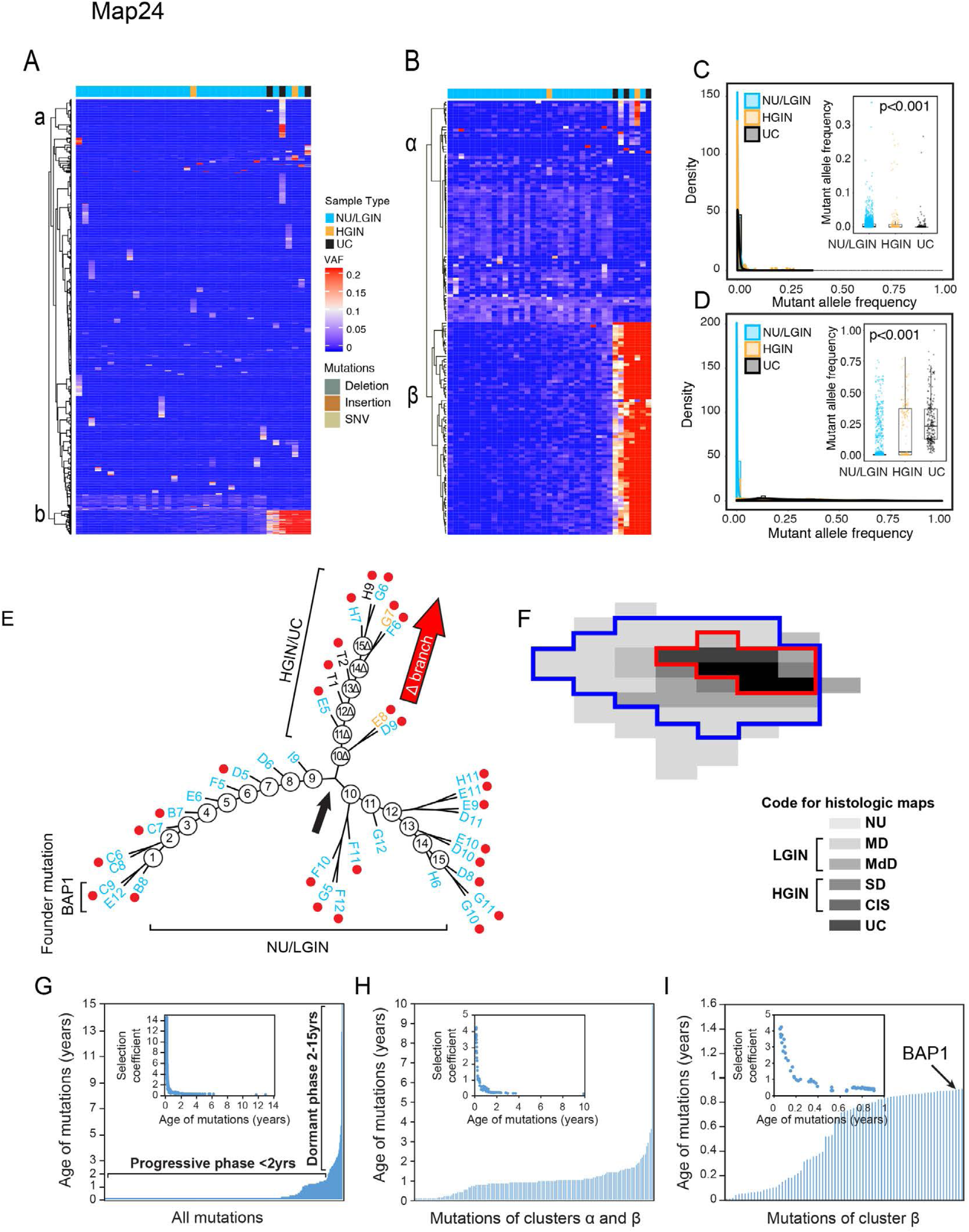
Clonal Enrichment of Mutations in Evolution of Bladder Cancer from Field Effect Along the Luminal Track. **(A)** Heatmap of non-silent mutations showing VAFs in individual mucosal samples. **(B)** Heatmap of VAFs in 157 genes showing variant alleles in at least three samples. **(C)** Density plot representing the clonality of non-silent VAFs in cluster **α** with similar low level of frequency which is decreasing in frequency with progression to HGIN and UC. **(D)** Density plot representing the clonality of non-silent VAFs in cluster **β** with a statistically significant increase in clonality with progression of HGIN and UC. Inset shows the boxplot analysis of VAFs in three groups of samples corresponding to NU/LGIN, HGIN, and UCs. **(E)** Parsimony analysis showing evolutionary tree with nine nodes of clonal expansions of successive clones in the field effect corresponding to NU/LGIN with major branching at node nine designated as Δ branch, which evolved to HGIN and UC. All foci of HGIN and UC evolved from successive waves of clonal expansion in branch Δ. The main NU/LGIN clone continued to evolve in successive waves of mutational changes (nodes 10-15) with complex branching pattern which did not progress to HGIN or UC. **(F)** Whole-organ histologic map showing a plaque-like area outlined by a blue line with a founder mutation of BAP1. The red line outlines an area corresponding to the wide-spread CNV alterations. **(G)** Age of all synonymous and nonsynonymous mutations predicted by mathematical modeling. Inset shows the selection co-efficient in relation to the predicted age. **(H)** Age of mutations in cluster **α** and **β** predicted by mathematical modeling. Inset shows the selection co-efficient of mutations of **α** and **β** clusters in relation to their age. **(I)** Age of mutations in cluster **β** predicted by mathematical modeling. Inset shows the selection co-efficient of mutations in the **β** cluster in relation to their age.

Hierarchical clustering demonstrated that VAFs of these genes formed two clusters with distinctive behaviors in the progression to carcinoma. The first clusters, designated as **α,** contained 80 variant alleles in the luminal map and 43 variant alleles in the basal map that showed a low mutational frequency patterns across the mucosa and even decreased in their frequencies with progression to HGIN and UC **(Fig. 3B, C and Extended Data Fig. 10B, C)**. Of the 80 luminal **α** cluster genes, 77 were altered by nonsynonymous single nucleotide substitutions resulting in amino acid change **(Extended Data Table 3)**. The remaining genes were altered by an insertion (one gene) and deletions (two genes) resulting in frameshifts. In the luminal map, cluster **α** was enriched with mutations in genes that control invasion and migration, including RASGRF1, which stimulates the disassociation of GDP from RAS,^31^ and DOCK7, which encodes a guanine nucleotide exchange factor that regulates the activity of the Rho family proteins,^32^ RAC1 and RAC3,^33^ DSG4, encoding a member of desmosomal cadherins,^34^ CEACAM7, encoding a surface glycoprotein and member of the carcinoembryonic antigen family,^35^ and ITGA7, encoding a member of the integrin alpha chain family.^36^

In the basal map, cluster **α** contained 43 variant alleles that showed a similar frequency pattern across the mucosa **(Extended Data Fig. 10C and Extended Data Table 4)**. The most frequently mutated genes in cluster **α** in the basal map were involved in transcriptional and cell cycle regulatory pathways and included genes involved in androgen receptor signaling, chromatin remodeling and DNA repair. Mutations in ZAP70 and POU2AF1 with their role as major regulators of T-cell and B-cell development and their activation signify field effect involvement of immune regulatory pathways.^37, 38^

The second clusters, designated **β,** contained 77 genes in the luminal map and 155 genes in the basal map that exhibited a significant increase in VAFs with evolution to HGIN and UC **(Fig. 3B, D, Extended Data Fig. 10B, D, and Extended Data Tables 5 and 6)**. Noteworthy cell cycle mutations in the luminal map included CDKN1A, a negative regulator of cyclin-cyclin-dependent kinase 2 controlling cell cycle progression^39, 40^ and FBXW7, which binds directly to cyclin E and targets its ubiquitin-mediated degradation,^41, 42^ and APC, a tumor suppressor protein implicated in colorectal cancers and an antagonist of the Wnt signaling pathway.^43, 44^

In the basal map the 155 mutated genes in cluster **β** also increased in their frequencies with progression of neoplasia **(Extended Data Fig. 10D and Extended Data Table 6)**. The members of transcriptional regulators and genes controlling cell cycle represented the largest group of mutated genes in this cluster and comprised of HORMAD1,^45^ CDK13,^46^ CDKN2A,^47^ BTG2,^48^ BATF^49^ among others. In addition to mutational change, CDKN2A is frequently inactivated in many cancers including bladder cancers by homozygous deletions and methylation.^47^ The second major group of mutated genes comprised of genes involved in signal transduction and metabolism. It involved ACSL6 regulating fatty acid metabolism^50^ and BAIAP3 a member of the secretin receptor family involved in angiogenesis.^51^ A distinctive group of genes mutated in basal cancer represented genes involved in chromatin remodeling, DNA repair and RNA regulation including 15 unique KMT2D and 10 unique KDM6A mutation, two of the most frequently mutated chromatin remodeling genes in bladder cancer. Interestingly several genes mutated in basal cancer were involved in inflammatory responses. These genes included TLR4/5, the toll-like receptor family members that play a fundamental role in pathogen recognition and activation of innate immune responses and PIK3AP1 involved in the survival of mature B cells and negatively regulating inflammatory cytokine production.^52, 53^

To further analyze the clonal architecture of the mucosal field effect and its evolution to cancer, we investigated the number of private and shared mutations in mucosal samples **(Fig. 12F)**. On average, 50 (ranging from 1-100 mutations) were shared by all samples but many areas of the bladder mucosa exhibited only distinct private mutations. In both maps there was an increase of shared mutations with progression of neoplasia to HGIN and UC **(Extended Data Figs. 10E and 11A)**. Kruskal-Wallis rank sum tests showed a statistically significant increase in VAFs comprising the mutations in clusters **β (Extended Data Figs. 10F and 11B)**.

Overall, the analyses of the geographic mutational landscapes in bladder mucosa disclosed remarkable similarity of cancers evolving along the luminal and basal tracks. It revealed the existence of pronounced mutational heterogeneity in mucosal field effects with significant change and clonal expansion of a unique set of genes in each of the maps with the disease progression to HGIN and UC.

We performed validation studies of the mutations identified in both maps focusing on mutations of clusters **α** and **β** since they appear to be involved in the development and progression of the disease. Seventeen mutations from clusters **α** and **β** in both maps were confirmed by manual PCR-based Sanger sequencing. In addition, two founder mutations, BAP1 and CAPRIN1, were confirmed in multiple mucosal samples by blocking PCR combined with Sanger sequencing.

Additional validation analyses of mutations in clusters **α** and **β** were performed on the TCGA cohort.^54, 55^ Among the genes from clusters, **α** the most frequently mutated genes identified in the luminal map were RB1 (16%), FAT3 (15%) and DNAHR (10%). Among the genes from the basal map, the most frequently mutated genes in the TCGA cohort were LRRK2 (11%) and ERCC2 (10%) **(Extended Data Figs. 12 and 13)**. Among the genes from cluster **β**, which were associated with the increase in clonality in the progression of the disease, the most frequently mutated genes of the luminal map were LRP1 (10%), BSN (8%) and CDKN1A (8%). The most frequently mutated genes identified in the basal map were KM2C (21%), KDM6A (24%), and BIRC6 (15%) **(Extended Data Figs. 14 and 15)**.

### Mechanism of Mutagenesis Driving the Development of Cancer from Field Effect

To characterize the evolution of the mutational signatures associated with progression from field effects to neoplasia to carcinoma, we analyzed the six single-base nucleotide substitutions (C> A, C>G, C>T, T>A, T>C, and T>G) and their context motifs in all mucosal samples of both maps.^56^ In both maps, the frequent C>T mutations increased at the point of evolution from NU/LGIN to HGIN and UC with significant changes in 20 different mutational signatures **(Extended Data Figs. 16A-C and 17A-C)**. The most prominent signatures in the luminal map (1, 13, 29, and 30) have been attributed to aging, APOBEC, tobacco use, and an unknown etiology, respectively **(Extended Data Fig. 16D-F)**. Similarly, signatures 1, 2, 3, 6, and 30, which have been attributed to aging, APOBEC, defective HR, defective MMR, and an unknown etiology, predominated in the basal map **(Extended Data Fig. 17D-F),** Signatures 1, 13, and 30 were dominant in progression to carcinoma in the luminal map with the APOBEC and 13 weight scores significantly increasing with progression of the disease. In order to evaluate the contribution of individual mutagenesis pattern to the mutational landscape of mucosal samples, we perform bootstrapping analysis and calculated the p-values to assess their significance (p<0.05 was considered statistically significant). The approach confirmed the dominance of signatures 1, 2, 6, 13, 29, and 30 in the luminal map **(Extended Data Fig. 16G)** and signatures 2, 6, 13, and 30 in the basal map **(Extended Data Fig. 17G)**.

### Evolution of Copy Number Changes from Field Effects to Carcinoma

We used genome-wide SNP microarrays to analyze copy number changes in the two cystectomy specimens.^3^ Changes in the gene copy numbers were less evident in areas of bladder mucosa corresponding to field effects and designated as NU/LGIN. Similar to the mutational changes, genome-wide copy number alterations increased with progression to HGIN and UCs **(Extended Data Figs. 18A, B and 19A, B)**. In both cases they formed clearly-defined plaques outlining areas of bladder mucosa which contained HGIN and UC with some adjacent areas of NU/LGIN **(Extended Data Figs. 18C and 19C)**. These patterns of loco-geographic distribution indicated the involvement of wide-spread copy number variations in later phases of tumor evolution associated with the progression to HGIN and UC. The copy number changes resulted in both gains and losses of multiple chromosomes that involved large segments of **p** and **q** arms and in many instances entire chromosomes. Clustering of mucosal samples using copy number variations and the Euclidean distance revealed in both instances two major clusters, designated **α** and **β**, segregated mucosal samples with low and high numbers of CNVs. Practically all of the UC and HGIN samples were contained in clusters **β**. Using the Hamming distance we detected significant differences in copy number variability among samples classified as HGIN/UC compared to NU/LGIN **(Extended Data Figs. 18D and 19D)**. The copy number changes in NU/LGIN were minimal and the notable exception was a copy number gain on chromosome 11q, which were present in a few samples microscopically classified as NU/LGIN and involved a frequently amplified gene in bladder cancer.

### Modeling of Carcinoma Evolution from Mucosal Field Effects

In order to understand the pattern of clonal evolution of cancer development from field effects, we used non-silent and silent mutations to construct cancer evolutionary trees. This revealed complex branching patterns with multiple waves of clonal expansion represented by at least 15 and 17 nodes in luminal and basal cancers respectively **(Fig. 3E and Extended Data Fig. 20A)**. In the luminal tumor there was a divergence at node 9 with the development of a branch designated as **Δ** comprising of nodes 10**Δ** through 15**Δ** which progressed to HGIN and UCs **(Fig. 3E)**. In the basal tumor the divergence occurred after node 12 where the main NU/LGIN clone evolved to HGIN and UCs **(Extended Data Fig. 20A)**. In the luminal tumor the main NU/LGIN clone continued to evolve after node 9 with a complex branching pattern which did not progress to HGIN or UC. In the basal tumor, successive clones of NU/LGIN at nodes 9, 12, 15, and 17 developed branches of clonal expansion which also did not progress to HGIN and UC. Branch Δ in the luminal tumor was characterized by an increase in mutations, their VAFs, as well as the proportion of shared mutations signifying their clonal expansion in the progression process **(Extended Data Figs. 11C-E and 20B-D)**. These were particularly evident for the number of mutations and VAFs in cluster **β** signifying their putative driver roles in the development of HGIN and UC **(Extended Data Figs. 11F, G and Fig. 20E, F)**. Likewise, in nodes 13-17 in the basal tumor, the mutations of cluster **β** showed dramatic increases in their numbers and VAFs confirming their putative driver roles in the development of HGIN and UC **(Extended Data Fig. 20B-F)**. We identified a founder mutation of the BAP1 gene^57^ in the luminal map and of the CAPRIN1 gene^58^ in the basal map **(Fig. 3E and Extended Data Fig. 20A)**. These mutations were present in the samples connected to node 1 in each map i.e. were present in areas of the bladder mucosa that represented the furthest genetic distances from node 9 in the luminal map and node 13 in the basal map that were the checkpoints for progression to HGIN and UC. These mutations were present in 76% of mucosal samples in the luminal map and in 69% of the mucosal samples in the basal map. Most importantly, they formed 74cm^2^ and 84cm^2^ plaques in the luminal and basal maps respectively which were involving large areas of bladder mucosa **(Fig. 3F and Extended Data Fig. 20G)**. We validated the presence of founder mutations in representative mucosal samples by PCR-based Sanger sequencing.

To address the question of how long it takes for bladder cancer to develop, we applied a mathematical modeling algorithm utilizing the mutational profile and a sequence of successive clonal evolution in the nodes of the parsimony tree by a time-continuous Markov branching process.^59^ This approach provides an estimation of progression on a time scale of the transformation process based on maximal parsimonious principles. Initially, we performed the modeling by using all nonsynonymous and synonymous mutations **(Fig. 3G and Extended Data Fig. 20H)**. These analyses showed that both maps evolved over approximately 10-15 years, and the age-related curves of mutations had left-skewed patterns with the early mutations being more than 10 years old. Based on the time scale, the processes could be divided into two major phases. The older (dormant) phase, in which mutations developed gradually over approximately a decade, involved mutations that were characterized by low selection coefficients consistent with the idea that they produced marginal proliferative advantage. The more recent (progressive) phases were less than two years old and were characterized by the accumulation of large numbers of mutations with high selection coefficients indicating that they were associated with clonal expansion and produced high proliferative advantage. We repeated the same analyses focusing on mutations characterized by the clonal expansion i.e. those of clusters **α** and **β**. The overall age-related pattern was similar in both maps spanning approximately 10 years. Again, there was a limited number of mutations, older than two years, corresponding to the dormant phase characterized by low selection coefficient with minimal proliferative advantage. A large proportion of these mutations were younger than two years with high selection coefficient corresponding to the progressive phase of the disease **(Fig. 3H and Extended Data Fig. 20I)**. When the mathematical modeling was restricted to clusters **β**, i.e. the mutations characterized by an increase in their clonality with disease progression and representing the putative drivers of the progression process, it showed that in both maps these mutations developed in the last year and produced high proliferative advantage as shown by the increased selection coefficient **(Fig. 3I and Extended Data Fig. 20J)**.

### Integrated Pathway Analysis

To evaluate global contributions of genome-wide RNA expression changes to bladder cancer development, we used a recently developed tumor messenger RNA expression score (TmS), which showed strong positive correlation with aggressive variants of cancer across the TCGA sites.^60^ In both maps there was a significant increase of TmS with disease progression to HGIN and UC **(Fig. 4A)**. Moreover, TmS of NU/LGIN samples in the basal map was higher as compared to NU/LGIN samples in the luminal map. This observation was validated on the TCGA cohort, which showed higher TmS in the clinically aggressive basal tumors as compared to the luminal tumors **(Fig. 4B)**.

**Fig. 4.**
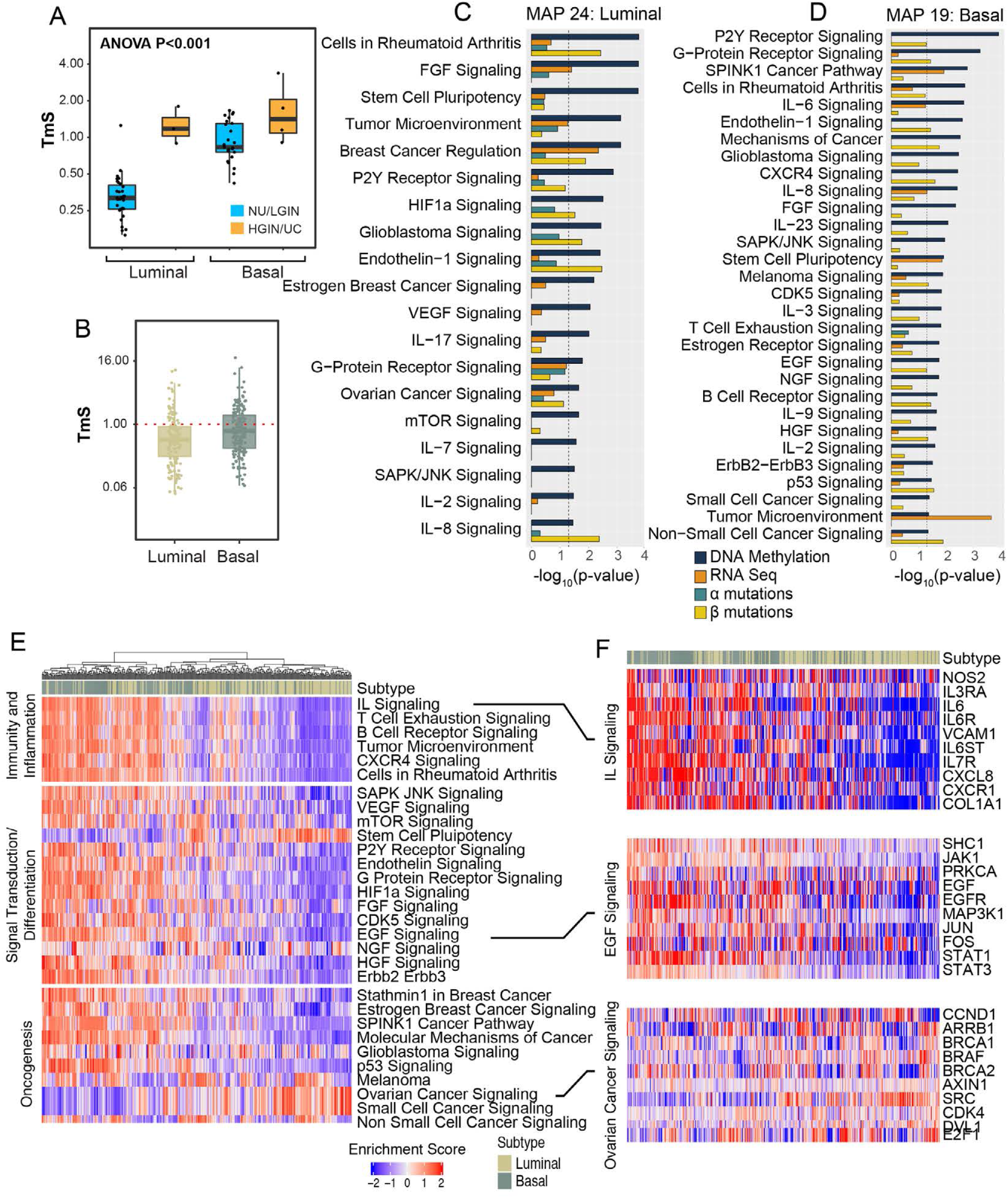
Interactive Analysis of BC Development from Field Effects with Validation in the TCGA Cohort. **(A)** Tumor specific mRNA expression (TmS) in the development of BC from field effects in the luminal and basal subtypes. **(B)** TmS in the luminal and basal subtypes of bladder cancer in the TCGA cohort (n=408). **(C)** Selected monotonically dysregulated pathways in field effects of the luminal map. **(D)** Selected monotonically dysregulated pathways in field effects of the basal map. **(E)** Enrichment scores of the regulons controlling immunity and inflammation, signal transduction/differentiation, and oncogenesis in molecular subtypes of bladder cancer in the TCGA cohort (n=408). **(F)** Expression pattern of selected genes in the regulons of interleukins, EGF, and ovarian cancer signaling in the molecular subtypes of BC in the TCGA cohort (n=408).

To identify pathways in the development of bladder cancer we used genome-wide expression and methylation levels of monotonically altered genes and complemented them with mutations of genes in clusters **α** and **β**. This approach identified multiple dysregulated canonical pathways within each platform. Subsequent integrative analysis identified 91 and 125 canonical pathways dysregulated in NU/LGIN, which were continuously dysregulated in the progression of HGIN and UC in luminal and basal maps respectively **(Extended Data Tables 5 and 6)**. Combined selected pathways in luminal and basal maps dysregulated in mucosal field effects are shown in **Fig. 4C,D**. These pathways can be classified into three major groups i.e. regulating immunity and inflammation, signal transduction and differentiation, and those involved in oncogenesis. Since many of these pathways were related to each other and some of them were altered in both maps, we arranged a list of selected 30 pathways dysregulated in field effects **(Fig. 4E)**. The striking dominant feature of mucosal field effect was the dysregulation of multiple pathways controlling immunity and inflammation. Dysregulation of signaling controlling multiple interleukins such as IL-2, 3, 6-8, 17, and 23 was the dominant feature complemented by changes of B and T cells signaling as well as changes controlling tumor microenvironment. Dysregulation of signal transduction pathways, many of which involve tissue differentiation program, included HIF1, VEGT, mTOR as well as EFG, NGF, and AGF signaling. Dysregulation of multiple oncogenic pathways, including p53, glioma, melanoma, breast cancer, and general cancer signaling signified the transformation effect synergistic with alterations in immunity and tissue differentiation programs as initiating changes in bladder carcinogenesis.

The analysis of 30 regulons controlling immunity and inflammation, signal transduction and tissue differentiation as well as oncogenesis in the TCGA cohort showed that they were frequently involved in bladder carcinogenesis. The activation of many of them was enriched specifically in luminal and basal cancers **(Fig. 4E,F)**.

## DISCUSSION

Our study provides the comprehensive characterization of mucosal field effects associated with the development of bladder cancer on the whole-organ scale. It showed complex genome-wide diffuse mucosal changes in DNA methylation and mRNA expression involving multiple pathways affecting immune responses, urothelial differentiation, and proliferation. Surprisingly, the field effect in microscopically intact tissue harbored dysregulations of multiple oncogenic pathways. It also revealed that the two major intrinsic molecular subtypes of bladder cancer are determined *de novo* i.e. the luminal cancer developed from luminal field effects whereas the basal form evolved from basal field effects. The field effects associated with the development of luminal cancer showed the retention of the luminal differentiation program however, it was altered showing dysregulations of multiple pathways involved in the differentiation of stratified epithelia. In contrast, the basal field effects showed suppression of the luminal program with diffuse mucosal downregulation of terminal urothelial differentiation exemplified by lower expression levels of uroplakins. These results are consistent with the animal models data showing that these two subtypes develop from distinct uroprogenitor cells.^61, 62^

The wide-spread plaque-like mRNA expression and DNA methylation changes in the mucosal field effect were in a background of highly heterogeneous alterations with low allele frequencies implicating that the mRNA and DNA methylation changes were the primary early events of carcinogenesis. Using loco-geographic distribution patterns of mutations and their allele frequencies, three distinct groups of mutations can be identified. The first group comprised of low frequency mutations restricted to individual mucosal samples. They very likely represented the progeny of individual uroprogenitor cells. The other two types of mutations were associated with clonal expansion and involved large areas of bladder mucosa including those with microscopically normal appearing urothelium. The first of these two groups, referred to as cluster **α** mutations, comprised 80 and 43 genes in luminal and basal maps. The mutations of cluster **α** were characterized by low VAFs comprising of typically less than 5% of cells in the urothelium. These mutations had a stable frequency which did not increase with the progression process. In fact, in one of the maps it appeared that this clone was being eliminated with the progression to HGIN and UC. We hypothesized that the clone of cells with these mutations sets the stage for carcinogenesis by disturbing urothelial differentiation, providing growth advantage and dysregulating immunity. It appears that the cells with these mutations harmoniously co-existed with other urothelial cells and were microscopically normal appearing. In contrast, mutations of clusters **β**, comprising of 77 and 155 mutations in luminal and basal maps, increased substantially with disease progression to the fully malignant phenotype supporting their driver’s role in the process. Mutational and clonality analyses identified founder mutations of clones **β** involving BAP1 and CAPRIN1 genes in luminal and basal maps. Consistent with their founder and contributory driver’s role, these mutations formed continuous large plaques involving most of the bladder mucosa and dramatically increased in their frequencies with progression. Time modeling revealed that bladder carcinogenesis is a prolonged and clinically occult process spanning 10-15 years. The long dormant phase lasting approximately 10 years is succeeded by an accelerated progressive phase during the last 1-2 years before the development of clinically symptomatic invasive cancer. In both maps the final progressive phase of the disease appeared to be driven by **β** clone mutations with high proliferative advantage.

Our data supports the concept that the wide-spread dysregulation of mucosal immune environment involving multiple interleukins plays an important role in bladder cancer initiation. The interleukins act in a background of dysregulated urothelial differentiation program and multiple activated oncogenic pathways causing an irreversible damage, which precipitates in the development of clinically aggressive invasive cancer.

## Acknowledgements

This study was supported by NCI Genitourinary Bladder SPORE Grant P50CA 91846 (Project 1 and Core C), MDACC Nathan W. Lassiter Endowment; as well as Donor Funds (B.C.); the Polish National Science Centre grants No. 2018/29/B/ST7/02550 (M.K.) and 2016/23/D/ST7/03665 (R.J.); and National Institute of Health grants R01HL136333 and R01HL134880 (Y.C.).

## MATERIAL AND METHODS

### Preparation of whole organ map and DNA extraction

Human samples as well as clinical data were collected and archived according to the laboratory protocol approved by the Institutional Review Board of The University of Texas MD Anderson Cancer Center. The whole-organ histologic and genetic mapping (WOHGM) was performed on the radical cystectomy specimens from nine Caucasian male patients with the mean age 73.6 years (range from 55 to 86 years) with high-grade muscle invasive (T3) urothelial carcinoma (Table X). The preparation of cystectomy specimens for DNA/RNA extraction and whole-organ histologic mapping follows the four steps illustrated **Extended DataFig. 1.**^63, 64^

**Step 1:** Each cystectomy specimen was opened longitudinally along the anterior wall of the bladder and pinned to a paraffin block. Then the mapping grid was applied and pressed down against the bladder mucosa with mechanical screws which provided sealed wells that separated mucosal areas and tumors into 1x2cm (2cm^2^) rectangles. The grooves at the bottom of the mapping grid preserved the urothelium for microscopic examination.

**Step 2:** Phosphate-buffered saline (PBS) (0.4 ml) was poured into each well, and the surface urothelium was scraped with a custom-designed metallic scraper. A small proportion of this fluid containing urothelial or tumor cells (approximately 20 μl) was used for cytospin preparations to assess the purity of cell suspensions for DNA and RNA extractions. In areas which contained visible tumor, the tissue was dissected directly from the bladder.

**Step 3:** Trizol reagent (1.6 ml) was poured into each mapping well, and the fluid was collected into separate Eppendorf tubes labeled with numbers corresponding to individual mapping wells of bladder mucosa. The Trizol reagent fluid contained lysed cells and was used for DNA/RNA extraction. The samples with tumor tissue were cut into small pieces and treated with Trizol.

**Step 4:** In the final step, each well was washed with PBS, and the mapping grid was removed from the surface of the bladder, which was then fixed in formalin overnight. The mapping grid left a permanent impression on the bladder surface, and its grooves preserved the urothelium for histologic mapping of the entire mucosa. After formalin fixation, each piece of bladder surface along with the preserved tissue was embedded in paraffin. One section from each block was stained with hematoxylin and eosin to evaluate the distribution of microscopically normal urothelium, *in situ* precursor lesions, and urothelial carcinoma microscopically. The precursor intraurothelial lesions were dichotomized into low- and high-grade categories and referred to as low-grade intraurothelial neoplasia (LGIN) and high-grade intraurothelial neoplasia (HGIN) as previously described. Map samples containing tumor tissue were classified according to the two-tier histologic tumor grading system of the World Health Organization referred to as low- and high-grade.^65^ The growth pattern of papillary versus non-papillary (or solid), and the depth of invasion were also recorded. Levels of invasion were defined according to the TNM staging system.^66^

There were two steps of quality check controls. The first step comprised of the overall assessment of the cystectomy specimens in terms of the representation of the full spectrum of precursor lesions and tumor samples as well as purity of urothelial and tumor cell preparations. Only those cystectomy specimens showing intact areas of cancer involving no more than 50% of the bladder mucosa are accepted for whole-organ mapping. Moreover, a cystectomy specimen accepted for likely informative mapping showed bladder mucosa with intact surface urothelium in over 90% of the sampling wells, and the full spectrum of microscopically recognizable precursor intraurothelial lesions. In addition, only those samples that contained more than 90% microscopically recognizable intact urothelial or tumor cells were used for DNA/RNA extraction yielding 5 to 70 mg of DNA.

The second step comprised of the assessment of the quality of the final DNA/RNA preparations for genomic profiling. The quality of DNA/RNA preparations was verified using NanoDrop, Bioanalyzer, and Qubit. Using these quality checks, only those whole-organ cystectomy specimens which contain a sufficient amount of high quality of DNA and RNA in at least 90% of mucosal samples of the cystectomy were accepted for genomic profiling. This procedure provided approximately 30-40 DNA/RNA samples per cystectomy corresponding to specific mucosa areas yielding 10-15 μg of DNA and 25-30 μg of RNA per sample.

### RNA Sequencing and Data Analysis

The RNA integrity and RIN number was assessed using a 2100 Bioanalyzer (Agilent). RNA concentration was determined using RiboGreen quantification (Quant-iT RiboGreen RNA Assay Kit from Invitrogen). RNA samples meeting a quantity threshold of 1 µg with the RNA integrity number (RIN) ≥7 were analyzed by the Advanced Technology Genomics (ATCG) Core. Prior to RNA library construction ribosomal RNA was removed from total-RNA preparations and cDNA synthesis utilizing oligo-d(T) and random hexamers was performed. The library was made up of random fragments that represent the entire sample. It was created by shearing DNA into 150-400 base fragments which were ligated to specific adapters. Following a sample cleanup step, the resultant library was quantified by qPCR and checked for quality using the Agilent TapeStation. The analyses were performed on 79 RNA samples from 74 mucosal samples from two maps (34 samples from Map 19 and 40 samples from Map 24) and from five sex-matched normal control urothelial suspensions, which were prepared from the ureters of nephrectomy specimens that were free of urothelial neoplasia.^67^

Quality control was conducted using RSeQC^68^ and FastQC. Sequencing reads were aligned to the GRCh38 reference genome using STAR v2.7.3a^69^, with GENCODE v32 transcript annotations.^70^ Read counts for individual genes were obtained using featureCounts from the Subread package.^71^ On average we obtained 27 mln reads per sample associated with more than 58 thousand unique genes, both coding and non-coding.

From this set 44 thousand genes showing more than 10 reads in at least one sample were selected and used in the subsequent analysis. Genes differentially expressed between specific sample groups were identified with DESeq2 v.1.26.0^67^ (Wald test), using a design formula that includes batch effect correction. In all tests we used Benjamini and Hochberg correction for multiple testing.^72^

For the assessment of luminal and basal phenotypes we used the expression levels of previously developed 28 luminal and 20 basal marker genes. For the quantitative assessment of molecular subtypes of bladder cancer we used previously developed basal to luminal transition (BLT) score.^73^ In brief, for the assessment of the luminal phenotype we used the 14 luminal markers from the original classifier.^14,40^22,30 In order to increase the power of our analyses these markers were complemented by 14 PPARγ target genes previously shown to be significantly enriched in luminal cancers.^14,40^22,30 Similarly, for the assessment of the basal phenotype, we used the 9 basal markers from the original classifier and complemented them with additional 11 p63 target genes which were shown to be significantly enriched in basal cancers.^14,40^22,30 Linear discriminant analysis (LDA) was performed to assess the power of individual markers to identify molecular subtypes of bladder cancer.^74^44 The unidimensional BLT score was defined as ∑ *W*_*i*_ ∗ *E*_*i*_, where *W*_*i*_ is the negative coefficient of linear discriminant (LD) and *E*_*i*_ is the expression of marker genes. Then a least absolute shrinkage and selection operator (LASSO) analysis was used to select the best 16 luminal and 12 basal markers to combat multicollinearity.^75^ Specifically, LASSO applied the L1 parameter as a constrain on the sum of the absolute values of the model parameters. In the process, 28 genes with a non-zero coefficient after the regularization process were selected for the calculation of the BLT score. We used the TCGA cohort as a training set to build a LDA model with 28 selected genes.

To assess the status of EMT in the evolution of bladder cancer from mucosal field effects we first analyzed the expression levels of signature transcription factors involved in the activation of EMT of SNAIL, TWIST, ZEB, FOX, SOX, and KLF families complemented with the analyses of homotypic adhesion molecules such as E-cadherin (CDH1), claudin 1 (CLDN1), and tight junction protein 1 (TJP1). To quantitatively assess the level of EMT, we calculated the EMT score based on a 76-gene expression signature reported in Byers et al as previously described.^18, 76^ 21,38 For each sample, the score was calculated as a weighted sum of 76 gene expression levels: 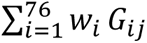 where *w*_*i*_ is the correlation coefficient between the *i*th gene expression in the signature and that of E-cadherin and *G*_*ij*_ is the *i*th gene’s normalized expression in the *j*th sample. We centered the scores by subtracting the mean across all tumor samples so that the grand mean of the score was zero.

To analyze immune gene expression signatures of bladder cancer evolution from field effect dendrogram nodes corresponding to genes expressed in specific immune cell types were identified through DAVID functional annotation clustering and Ingenuity Systems (www.ingenuity.com) analysis. The immune expression signature was quantitatively assessed by calculating the immune scores for the expression profile of 128 genes as previously described.^18^ Specifically, the immune score for the *i*th sample was defined as 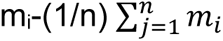, where *m_i_* is the median expression level across the *i*th sample’s immune expression profile and 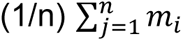 is the grand mean of medians across all n samples. Additional analysis of immune infiltrate was performed by the CIBERSORT algorithm (http://cibersort.standford.edu/runcibersort.php). The expression profile of 547 genes using normalized mRNA levels with absolute mode and default parameters was used to assess the presence of 22 immune cell types51.^77^ An empirical p value was calculated using 500 permutations to test against the null hypothesis that no cell type is enriched in each sample. Then a Fisher Exact test was used to test against the null hypothesis of no association between sample types and their statistical significance.

### Copy Number Genotyping and Data Analysis. Illumina microarrays

Quality control and pre-processing of Illumina Infinium Omni2.5-8 microarray data was conducted using Illumina Genome Studio ver. 2.0, using library files provided by Illumina (ver. 1.4-a1), with GRCh37 reference genome coordinates. In total 64 samples (32 for both map19 and map24) were studied as a single batch. Log R ratios and B-allele frequencies were obtained with the cnvPartition Genome Studio plugin (ver. 3.2.0), using GC wave adjustment option. Log R ratios and B-allele frequencies were exported from Genome Studio as a single table in wide format and then split into sample specific files using kcolumn script, which is a part of the PennCNV software (ver. 1.0.5).^78^ After adjusting the column order in the data files using our custom shell script, detection of copy number variable regions was conducted using OncoSNP ver. 1.4.^79^ OncoSNP was executed for tumor-normal sample pairs defined in the batch-file with both stromal contamination and intra-tumor heterogeneity models. Log R ratios and B-allele frequency plots were created using OncoSNP for all chromosomes. Additional visualizations of CN-altered regions, based on .cnvs and .qc result files provided by OncoSNP, were created using our custom R functions.

### Methylation and Data Analysis

We used the MethylationEPIC BeadChip method, which allows to interrogate over 850,000 methylation sites quantitatively across the genome at single-nucleotide resolution. In brief we perform bisulfite conversion on 1 µg of genomic DNA using EpiTech kit from Qiagen according to the manufacturer’s instructions. Bisulfite-converted DNAs were enzymatically fragmented prior to hybridization to BeadChip arrays. BeadArrays were scanned using the Illumina iScan technology to produce IDAT files. We preprocessed and normalized the raw IDAT files for each sample using the minfi R package (version 1.30.0.^80, 81^ CpG sites that were on the sex chromosome, cross-reactive, and with SNPs were filtered out. Gene-level methylation data were summarized based on Spearman correlations between the CpG sites within each gene and its expression level based on the results of an external dataset, the TCGA bladder cancer tissue samples. We represented the methylation level for each gene as of the single probe that has the most negative correlation between methylation and expression in the 391 TCGA samples by interrogating either the 1st exon, 5’UTR, or up to 1500bp upstream from the transcription start site (TSS). The data set in total contained 14,744 genes with 36 samples in map 19 and map 24 respectively, and 8 normal controls for downstream analysis.

The analyses were performed on 72 mucosal samples from two maps (36 samples from each map) and from sex-matched normal control urothelial suspensions, which were prepared from the ureters of five nephrectomy specimens that were free of urothelial neoplasia. The methylation levels were transformed into log2 ratios comparing the samples in NU/LGIN, HGIN, and UC groups with the normal controls. Thresholds to identify hyper- and hypo-methylated genes were set at 0.8/-0.8 for all samples in each group of map 24. In map 19, the threshold to identify genes with field effect was set at 0.7/-0.7, and differentially methylated genes that occur at HGIN and UC groups were identified at threshold 0.6/-0.6. In particular, to identify genes with changes only in HGIN and UC groups the percentage of samples that pass the threshold was 80%. We also conducted SAM^82^ by comparing NU/LGIN vs. control, HGIN vs. control, and UC vs. control respectively. The permutation was set to 100 with the pre-specified q-value error control threshold to be 0.2. In the luminal map we identified 6,912 genes with methylation and 1,380 of them showed monotonic change related to the progression of the disease i.e. 125 genes showed hyper- or hypomethylation in NU/LGIN through HGIN to UC samples, two genes showed hyper- or hypomethylation in HGIN in UC samples, and 1,253 genes were hyper- or hypomethylated in UC samples. In the basal map, we identified 7,976 genes with methylation changes and 1,658 of them showed distinctive monotonic change related to the progression of the disease i.e. 427 genes showed hyper- or hypomethylation in NU/LGIN through HGIN to UC samples, 48 genes showed hyper- or hypomethylation in HGIN and UC samples and 1,183 genes were hyper- or hypomethylated in UC samples only.

### Whole-Exome Sequencing and Data Analysis

We used a sequencing pipeline in the Genomic Core Facility at MD Anderson Cancer Center and whole exome sequencing was performed on an Illumina NovaSeq6000 sequencer using high output flow cell with 150 nucleotides paired-end runs with an average coverage across the samples of 200X ± 75.5 SD. All steps, including Illumina library preparations, exome capture, Illumina sequencing, as well as downstream exome data processing, mutation calling and annotation were performed essentially as outlined in recent cancer sequencing project publications.^83, 84^ In brief, for data analysis BWA-MEM (version 0.7.12) to align reads to the GRCh38 reference genome was applied. Samtools (version 1.4) and Picard (version 2.5.0) were used to sort and convert between formats and remove duplicate reads.^85^ The Genome Analysis Toolkit (GATK, version 3.4-46) was used to generate realigned and recalibrated BAM files.^86^ MuTect2 and Oncotator (version 1.8.0.0) were used to identify non-silent and silent mutations and to produce function-based annotations of the single nucleotide variants (SNVs) and insertions/deletions (Indels).^87–89^

The mutations identified in two maps were analyzed in the TCGA cohort for which the mutational data were downloaded from the TCGA portal (https://portal.gdc.cancer.gov/). MutSigCV (version1.4; https://software.broadinstitute.org/cancer/cga/mutsig) was used to identify genes that were mutated more often than expected by chance given the background mutational process. The list of significant genes was obtained using a false discovery rate (FDR) cutoff of 0.05.^3^

### Mutational signatures

We used non-silent mutations identified in both maps which were present in at least one sample corresponding to the following base pair substitutions: C > A, C > G, C > T, T > A, T > C, T > G. Fisher’s exact test was used to test against the null hypothesis that they are equally distributed in the three groups of samples corresponding to NU/LGIN, HGIN, and UC. The genomic context of SNVs referred to as fingerprints which included the two flanking bases on 5’ and 3’ sides to each position for a total of 96 possible mutational patterns was assembled. Wilcoxon Rank Sum tests were used to test against the hypothesis of no difference in the frequency of any fingerprint between any two groups of mucosal samples. The Benjamini and Hochberg (BH) method was applied to control the false discovery rate (FDR).^72^ For each sample, we used its mutational fingerprints (V) and the quadratic programming method to compute a weight score (H) for each of 30 canonical mutational signatures (W) available from the Sanger Institute (https://cancer.sanger.ac.uk/cosmic/signatures): We took the 96 by 30 matrix of canonical signatures (W) and given the 96 by 1 mutational profile of a sample (V) we computed the 30 by 1 vector (H) for each of the canonical signatures’ relative contributions to the sample profile by solving the following optimization problem:

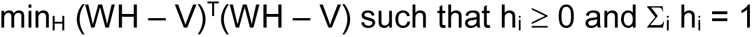

The optimization problems were solved in quadratic programming using the R package ‘‘quadprog’’ (version 1.5-5). Kruskal-Wallis test was used to verify the null hypothesis of no difference in weight scores among three groups of samples: NU/LGIN, HGIN, and UC. In order to assess the significance of the contribution of mutational signatures in individual mucosal samples we applied bootstrapping analysis. We resampled mutational fingerprints (V) for each sample with replacement and computed the weight score as above 2000 times. The one-sided empirical p value was calculated as the percentage of weight scores that were greater than or equal to the observed sample weight score in the resampling distribution.

### Phylogenetic Analysis and Modeling of Bladder Cancer Evolution from Field Effects

In order to reconstruct the phylogenetic tree we calculated the Hamming distances among mucosal samples using a binary matrix of all non-silent and silent mutations present in at least one sample and applied the maximum parsimony method.^90, 91^ In this representation, each node models a population of cells: the length of the edge connecting two nodes is proportional to the number of mutated genes while a branch represents a time point in the evolution where two distinct populations emerge. The length of the branch is proportional to the number of mutated genes that are private for each population.

To reconstruct the cancer time evolution from mucosal field effects, the time-continuous Markov branching process with immigration and parsimonious principles was used as previously described.^92^ In brief, a mutation *j* appears at time 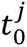 in a progenitor cell of the urinary bladder urothelial lining and gives rise to a mutant clone. Mutant cells divide at rate λ(1/yr) and after division, one cell enters self-renewal and the other differentiates with probability 1 −*s_j_*, or both cells enter self-renewal with probability *s_j_*. As a consequence, the mutant clone grows exponentially as exp(*λs_j_t*), where *t* is the age of *j*-th mutant’s clone counted from 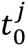. The secondary clones expand involving different areas of bladder mucosa at the times 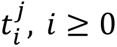 modeled by a stochastic Poisson process with intensity *v* (1/yr).^93^ If we denote the expected cell counts in the successive *j*-th mutant clones by 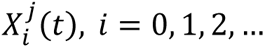, and the number of haploid genomes in normal uroprogenitor cells by 2*N*, the corresponding variant allele frequencies 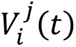 are defined as the ratios 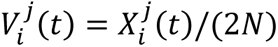 and are computed as:^92^

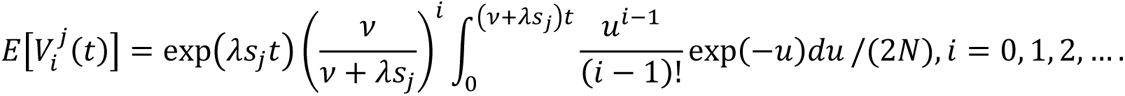

For any mutation *j* of age *t*_*i*_, the sequence of expectations 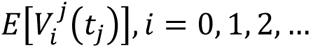, was computed to estimate the coefficients *a_j_* = *λs_j_t_j_* and *b_j_* = *vt_j_*. The coefficient *c*= 2*N* is a constant parameter representing an estimate of the number of uroprogenitor cells in the sampled area. The computations were performed for 10^2^ − 10^5^ uroprogenitor cells of the sampled mucosal area, which did not significantly change the results, but the best fit was obtained with 5 × 10^3^ of uroprogenitor cells for which the data are presented. With the cell division rate *λ* and migration rate *ν*, parameter *b*_*j*_ is the proxy for mutation age *t*_*j*_, while the ratio *a*_*j*_/*b*_*j*_ is the proxy, for selection coefficient *s*_*j*_. We used the fitting algorithm with the optimization programs fminsearch and fminbnd in MATLAB programming language to estimate the sequence of mutations.^94–96^ The resulting estimates were presented as bar diagrams representing the age of mutations and point charts of the corresponding selection coefficients.

### Control and Reference Samples

The mutational and copy number variation analyses were performed in comparison to paired normal genomic DNA of the same patient. Normal genomic DNA was extracted from the buffy coat of peripheral blood after Ficoll centrifugation. Additional normal genomic DNA was extracted from bladder smooth muscle. Samples of bladder smooth muscle (2-3 each measuring approximately 1cm^3^) were snap-frozen. The absence of tumor infiltration was confirmed by microscopic examination of H&E frozen sections subjected to DNA extraction. For the analysis of gene expression and methylation status, we used normal urothelium obtained from ureters of nephrectomy specimens performed for renal cell carcinoma confined to the kidney. Ureters were open longitudinally, pinned down to the paraffin block, and the urothelial surface was scraped with a surgical blade. The quality and purity of urothelial cell suspensions were checked by cytospin preparations and the samples were snap-frozen in PBS for subsequent DNA/RNA extraction.

### Verification of Mutations by PCR

Selected 17 mutations from clusters **α** and **β** (CDKN1A, APC, FBXW7, BRAF, RCC1, PLCB3, PACS1, OTOP1, PCDH10, P53, KDM6A, ERCC2, FBXW7-2, CDKN2A, KMT2C, SMARCA4, and ZMIZ1) were verified by PCR-based Sanger sequencing. Low VAFs of BAP1 and CAPRIN1 founder mutations were verified by blocking PCR.^97^ Briefly, NEB Taq (M0273S) was used to amplify 25ng gDNA from non-repliG amplified gDNA. Flanking primers were used at 0.4 uM while the blocking primer was at 4.0 uM. Two-step PCR (34 cycles at 95°C for 30” and 62.8°C for 30” to amplify CAPRIN1 and 34 cycles at 95°C for 30” and 58.6°C for 30” for BAP1) were used with the following primers: CAPRIN1-F GTTTTGGTCACCTTTGCAGTTCATT, CAPRIN1-R AGTGATCCTCCCATCTCAGC, CAPRIN1-B TTGCAGTTCATTCTGAATCTAGACTTGCTCAaaaa, and CAPRIN1-Seq CCATCTCAGCCTCCTAAAGTACTAGG and BAP1-F GCTAGTCTTGATGGACAGAGGAATT, BAP1-R CCCTTGCTTCACATCTTCTCG, BAP1-B AGAGGAATTGAGAGGTCCTTCTGGataa, and BAP1-Seq ATTGAGCGGTTCTGCTGATG. The amplified products were sequenced by Sanger method using AB373OXL sequencer.

### Calculation of Tumor-Specific Total mRNA Expression Score (TmS)

In order to assess the effect of global RNA expression changes to the development of BC from field effects, we used a recently developed tumor-specific total mRNA expression score (TmS) modified as follows.^98^ Since we wanted to estimate mRNA changes in microscopically normal appearing urothelium harboring field effects and in *in situ* intraurothelial precursor lesions progressing to invasive BC, we define the TmS as the ratio of total mRNA per cell or urothelial cells of interest over the normal control urothelium. We used *GTL_cell type of interest_* to denote the total mRNA transcript level per cell of the cell type of the interest and *GTL_normal control_* to denote the total mRNA transcript level per cell of the control urothelium. We calculated the TmS as 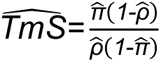, where 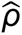 is the estimated purity of the cell type of interest from pathological review, and 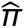 is the total mRNA proportion of the cell type of interest computed by the RNAseq deconvolution method DeMixT using data normalized at the seventy-fifth percentile based on the DSS package.^98–100^

## EXTENDED DATA FIGURE LEGENDS

**Extended Data Fig. 1. Gene Expression Changes in Bladder Cancer Evolving from Field Effects Along the Luminal and Basal Tracks. (A)** Heatmap showing the expression patterns of genes in cystectomy with luminal cancer. **(B)** Heatmap showing the expression patterns of genes in cystectomy with basal cancer.

**Extended Data Fig. 2. Evolution of Expression Changes from Field Effects Through HGIN to UC in Cancer Developing Along the Basal Track. (A)** Hierarchical clustering using top 60 downregulated and overexpressed genes showing monotonic expression change in samples corresponding to NU/LGIN through HGIN to UC, HGIN and UC and UC only. **(B)** Whole-organ expression map of chromosome 11 showing chromosomal diagram and expression pattern of genes in individual samples of cystectomy specimen classified as NU/LGIN, HGIN, and UC. **(C)** Expression pattern of downregulated and overexpressed genes with monotonic change involving NU/LGIN, HGIN, and UC. **(D)** 3D pattern of downregulated and overexpressed genes as it relates to the whole-organ map of a cystectomy specimen shown below filtered as in panel **C**.

**Extended Data Fig. 3. Expression Patterns of the Top 10 Overexpressed and Downregulated Genes Identified in the Early Field Effects in the TCGA Cohort. (A)** Expression pattern of the top 10 upregulated genes in the field effect of the luminal map. **(B)** Expression pattern of the top 10 downregulated genes in the field effect of the luminal map. **(C)** Expression pattern of the top 10 upregulated genes in the field effect of the basal map. **(D)** Expression pattern of the top 10 downregulated genes in the field effect of the basal map. **(E)** Expression levels of the top 10 upregulated genes in the field effect of the luminal map. **(F)** Expression levels of the top 10 downregulated genes in the field effect of the luminal map. **(G)** Proportion of cases with upregulation of the top 10 upregulated genes in luminal and basal subtypes of bladder cancer identified in the luminal map. **(H)** Proportion of cases with downregulation of the top 10 downregulated genes in luminal and basal subtypes of bladder cancer identified in the luminal map. **(I)** Expression levels of the top 10 upregulated genes in the field effect of the basal map. **(J)** Expression levels of the top 10 downregulated genes in the luminal and basal subtypes of bladder cancer identified in the basal map. **(K)** Proportion of cases with upregulation of the top 10 upregulated genes in luminal and basal subtypes of bladder cancer identified in the basal map. **(L)** Proportion of cases with downregulation of the top 10 downregulated genes in luminal and basal subtypes of bladder cancer identified in the basal map.

**Extended Data Fig. 4. EMT in the Evolution of Bladder Cancer from Field Effects Along the Luminal and Basal Tracks. (A)** Expression pattern of selected EMT-related genes an EMT score in mucosal samples of cystectomy specimens with luminal cancer. **(B)** Expression pattern of selected EMT-related genes an EMT score in mucosal samples of cystectomy specimens with basal cancer.

**Extended Data Fig. 5. Immune Landscape of Bladder Cancer Evolving from Field Effects Along the Luminal and Basal Tracks. (A)** Expression pattern of immune-related genes in mucosal samples of cystectomy specimens with luminal cancer. **(B)** Expression pattern of immune-related genes in mucosal samples of cystectomy specimens with basal cancer.

**Extended Data Fig. 6. Immune Microenvironment Based on Regulatory Genes in the Evolution of Bladder Cancer Along the Luminal and Basal Tracks. (A)** Expression pattern of immune regulatory genes in mucosal samples of cystectomy specimens with luminal cancer. **(B)** Expression pattern of immune regulatory genes in mucosal samples of cystectomy specimens with basal cancer.

**Extended Data Fig. 7. Gene Methylation Changes in Bladder Cancer Evolving from Field Effects Along the Luminal and Basal Tracks. (A)** Heatmap showing the methylation patterns of genes in cystectomy with luminal cancer. **(B)** Heatmap showing the methylation patterns of genes in cystectomy with basal cancer.

**Extended Data Fig. 8. Evolution of Methylation Changes from Field Effects Through HGIN to UC in Cancer Developing Along the Basal Track. (A)** Hierarchical clustering using top 52 hypo- and hypermethylated genes showing monotonic expression change in samples corresponding to NU/LGIN through HGIN to UC, HGIN and UC and UC only. **(B)** Whole-organ expression map of chromosome 13 showing chromosomal diagram and methylation pattern of genes in individual samples of cystectomy specimen classified as NU/LGIN, HGIN, and UC. **(C)** Methylation pattern of hypo- and hypermethylated genes with monotonic change involving NU/LGIN, HGIN, and UC. **(D)** 3D pattern of hypo- and hypermethylated genes as it relates to the whole-organ map of a cystectomy specimen shown below filtered as in panel **C**.

**Extended Data Fig. 9. Methylation Patterns of the Top 10 Hypermethylated and Hypomethylated Genes Identified in the Early Field Effects of the Two Maps Validated in the TCGA Cohort. (A)** Methylation pattern of the top 10 hypermethylated genes in the field effect of the luminal map. **(B)** Methylation pattern of the top 10 hypomethylated genes in the field effect of the luminal map. **(C)** Methylation pattern of the top 10 hypermethylated genes in the field effect of the basal map. **(D)** Methylation pattern of the top 10 hypomethyated genes in the field effect of the basal map. **(E)** Methylation levels of the top 10 hypermethylated genes in the field effect of the luminal map. **(F)** Methylation levels of the top 10 hypomethylated genes in the field effect of the luminal map. **(G)** Proportion of cases with hypermethylation of the top 10 hypermethylated genes in luminal and basal subtypes of bladder cancer identified in the luminal map. **(H)** Proportion of cases with hypomethylation of the top 10 hypomethylated genes in luminal and basal subtypes of bladder cancer identified in the luminal map. **(I)** Methylation levels of the top 10 hypermethylated genes in the field effect of the basal map. **(J)** Methylation levels of the top 10 hypomethylated genes in luminal and basal subtypes of bladder cancer identified in the basal map. **(K)** Proportion of cases with hypermethylation of the top 10 hypermethylated genes in luminal and basal subtypes of bladder cancer identified in the basal map. **(L)** Proportion of cases with hypomethylation of the top 10 hypomethylated genes in luminal and basal subtypes of bladder cancer identified in the basal map.

**Extended Data Fig. 10. Clonal Enrichment of Mutations in Evolution of Bladder Cancer from Field Effect Along the Basal Track. (A)** Heatmap of non-silent mutations showing VAFs in individual mucosal samples. **(B)** Heatmap of VAFs in 198 genes showing variant alleles in at least three samples. **(C)** Density plot representing the clonality of non-silent VAFs in cluster **α** with similar low level of frequency which is decreasing in frequency with progression to HGIN and UC. **(D)** Density plot representing the clonality of non-silent VAFs in cluster **β** with a statistically significant increase in clonality with progression of HGIN and UC. Inset shows the boxplot analysis of VAFs in three groups of samples corresponding to NU/LGIN, HGIN, and UCs. **(E)** Proportion of shared mutations in individual mucosal samples. **(F)** Density plot of VAFs in clusters **α** and **β** analyzed for their significance in progression to HGIN and UC by Kruskal-Wallis rank sum test.

**Extended Data Fig. 11. Evolution of Mutations from Field Effect Along the Luminal Track. (A)** Proportions of shared mutations in individual mucosal samples. **(B)** Density plot of VAFs in clusters **α** and **β** analyzed for their significance in the progression to HGIN and UC by Kruskal-Wallis sum rank test. **(C)** Number of mutations in nodes 1-15 and Δ branch. **(D)** Variant allele frequency (VAF) of all nonsynonymous mutations in nodes 1-15 and Δ branch. **(E)** Proportion of shared mutations in nodes 1-15 and Δ branch. **(F)** Number of mutations of cluster β in nodes 1-15 and Δ branch. **(G)** VAFs of mutations of cluster β in nodes 1-15 and Δ branch.

**Extended Data Fig. 12. Mutational Landscape of 80 Mutations with Variant Allele Frequencies Showing Mutations in at Least Three Samples of the Whole-Organ Luminal Map Analyzed in the TCGA Cohort.** Mutations of 80 genes of cluster **α** identified in the whole-organ luminal map and shown in cluster **α** of **Figure 3B** and analyzed in the TCGA cohort. Bars on the right show the number of specific substitutions for individual genes. The bars on the left side show the frequency of mutations in luminal and basal subsets of bladder cancer.

**Extended Data Fig. 13. Mutational Landscape of 43 Mutations with Variant Allele Frequencies** Showing **Mutations in at Least Three Samples of the Whole-Organ Basal Map Analyzed in the TCGA Cohort.** Mutations of 43 genes of cluster **α** identified in the whole-organ basal map and shown in cluster **α** of **Extended Data Fig. 10B** and analyzed in the TCGA cohort. Bars on the right show the number of specific substitutions for individual genes. The bars on the left side show the frequency of mutations in luminal and basal subsets of bladder cancer.

**Extended Data Fig. 14. Mutational Landscape of 77 Mutations with Variant Allele Frequencies Showing Mutations in at Least Three Samples of the Whole-Organ Luminal Map Analyzed in the TCGA Cohort.** Mutations of 77 genes of cluster **β** identified in the whole-organ luminal map and shown in cluster **β** of **Figure 3B** and analyzed in the TCGA cohort. Bars on the right show the number of specific substitutions for individual genes. The bars on the left side show the frequency of mutations in luminal and basal subsets of bladder cancer.

**Extended Data Fig. 15. Mutational Landscape of 155 Mutations with Variant Allele Frequencies Showing Mutations in at Least Three Samples of the Whole-Organ Basal Map Analyzed in the TCGA Cohort.** Mutations of 155 genes of cluster **β** identified in the whole-organ basal map and shown in cluster **β** of **Extended Data Fig. 10B** and analyzed in the TCGA cohort. Bars on the right show the number of specific substitutions for individual genes. The bars on the left side show the frequency of mutations in luminal and basal subsets of bladder cancer.

**Extended Data Fig. 16. Mutagenesis Patterns as they Evolve from Field Effect in Cancer Developing Along the Luminal Track. (A)** Composite bar graphs showing the distribution of all nucleotide substitutions in relation to cancer evolution from NU/LGIN through HGIN to UC. It shows statistically significant increase in C>T mutations (p<0.001) that parallel the evolution to HGIN and UCs. **(B)** Proportion of CNVs in specific nucleotide motifs for each category of substitution in three sets of samples corresponding to NU/LGIN, HGIN, and UC. **(C)** False discovery rate (FDR) for specific nucleotide motifs in the progression of neoplasia from NU/LGIN through HGIN to UC. **(D)** Weight score of mutagenesis patterns in three groups of samples corresponding to NU/LGIN, HGIN, and UC. **(E)** Weight scores of mutagenesis patterns in individual samples of bladder mucosa. **(F)** Statistical significance (p-value) of mutational patterns in progression of neoplasia from NU/LGIN through HGIN to UC. **(G)** Significance of contributions for mutagenesis signatures in individual samples after bootstrapping. Blue boxes indicate p<0.005.

**Extended Data Fig. 17. Mutagenesis Patterns as they Evolve from Field Effect in Cancer Developing Along the Basal Track. (A)** Composite bar graphs showing the distribution of all nucleotide substitutions in relation to cancer evolution from NU/LGIN through HGIN to UC. It shows statistically significant increase in C>T mutations (p<0.001) that parallel the evolution to HGIN and UCs. **(B)** Proportion of CNVs in specific nucleotide motifs for each category of substitution in three sets of samples corresponding to NU/LGIN, HGIN, and UC. **(C)** False discovery rate (FDR) for specific nucleotide motifs in the progression of neoplasia from NU/LGIN through HGIN to UC. **(D)** Weight score of mutagenesis patterns in three groups of samples corresponding to NU/LGIN, HGIN, and UC. **(E)** Weight scores of mutagenesis patterns in individual samples of bladder mucosa. **(F)** Statistical significance (p-value) of mutational patterns in progression of neoplasia from NU/LGIN through HGIN to UC. **(G)** Significance of contributions for mutagenesis signatures in individual samples after bootstrapping. Blue boxes indicate p<0.005.

**Extended Data Fig. 18. Evolution of Copy Number Changes from Field Effects to Carcinoma in the Luminal Map. (A)** Log R ratios (top panel) and B-allele frequencies (bottom panel) of all chromosomes in representative samples of bladder mucosa reflecting the progression pattern from NU/LGIN through HGIN to UC. **(B)** Expanded view of chromosome 3 showing Log R ratios (top panel) and B-allele frequencies (bottom panel) showing segmental loss of 3p in representative samples of bladder mucosa reflecting the progression pattern from NU/LGIN through HGIN and UC. **(C)** Histologic map of the cystectomy specimen. The red line outlines a plaque of bladder mucosa with wide-spread genomic changes of copy number variations. **(D)** CNV difference calculated as Hamming distance in the two groups of samples corresponding to NU/LGIN and HGIN to UC.

**Extended Data Fig. 19. Evolution of Copy Number Changes from Field Effects to Carcinoma in the Basal Map. (A)** Log R ratios (top panel) and B-allele frequencies (bottom panel) of all chromosomes in representative samples of bladder mucosa reflecting the progression pattern from NU/LGIN through HGIN to UC. **(B)** Expanded view of chromosome 11 showing Log R ratios (top panel) and B-allele frequencies (bottom panel) with segmental amplification 11q containing CDKD1 in representative samples of bladder mucosa reflecting the progression pattern from NU/LGIN through HGIN to UC. **(C)** Histologic map of the cystectomy specimen. The red line outlines a plaque of bladder mucosa with wide-spread genomic changes of copy number variations. **(D)** CNV difference calculated as Hamming distance in the two groups of samples corresponding to NU/LGIN and HGIN to UC.

**Extended Data Fig. 20. Reconstruction of the Evolutionary Tree in Cancer Developing Along the Basal Track. (A)** Parsimony analysis showing evolutionary tree with 17 nodes of clonal expansion of successive clones in the field effect corresponding to NU/LGIN. Node 13 signifies the progression to HGIN and all samples of HGIN and UC evolved through four additional waves of clonal evolution (nodes 14-17). **(B)** Number of all nonsynonymous mutations in mucosal samples of nodes 1-12 and nodes >12. **(C)** Variant allele frequency (VAF) of all nonsynonymous mutations in samples connected to nodes 1-12 and nodes >12. **(D)** Proportion of shared mutations in nodes 1-12 and >12. **(E)** Number of mutations in cluster **β** in nodes 1-12 and nodes >12. **(F)** VAFs of mutations in cluster **β** in nodes 1-12 and >12. **(G)** Histologic map of the cystectomy specimen. The blue line outlines a plaque of bladder mucosa with mutations of founder genes (CAPRIN1, MRE11A, PDE11A, OR2J3, and BIRC6). The red line outlines a plaque of bladder mucosa with widespread genomic changes of copy number variations (CNV). **(H)** Mathematical modeling of cancer evolution from field effect using all nonsynonymous mutations. Inset, shows selection coefficient related to age of mutations. **(I)** Mathematical modeling of cancer evolution from field effect using clonally expanding mutations of clusters **α** and **β**. Inset, shows selection coefficient related to age of mutations. **(J)** Mathematical modeling of cancer evolution from field effect using clonally expanding mutations of clusters **β**. Inset, shows selection coefficient related to age of mutations.

**Extended Data Fig. 21. RNASeq-based Pathways Monotonically Dysregulated in Mucosal Field Effects. (A)** Pathways dysregulated in the luminal map. **(B)** Pathways dysregulated in the basal map.

**Extended Data Fig. 22. Methylation-based Pathways Monotonically Dysregulated in Mucosal Field Effects. (A)** Pathways dysregulated in the luminal map. **(B)** Pathways dysregulated in the basal map.

**Extended Data Fig. 23. Mutations-based Pathways Monotonically Dysregulated in Mucosal Field Effects. (A)** Dysregulated pathways related to mutations of cluster α in the luminal map. **(B)** Dysregulated pathways related to mutations of cluster α in the basal map. **(C)** Dysregulated pathways related to mutations of cluster β in the luminal map. **(D)** Dysregulated pathways related to mutations of cluster β in the basal map.

