## Supplementary figures and images for "The Origin of Bladder Cancer from Mucosal Field Effects"

### Extended Data Fig. 1

#### Extended Data Fig. 1

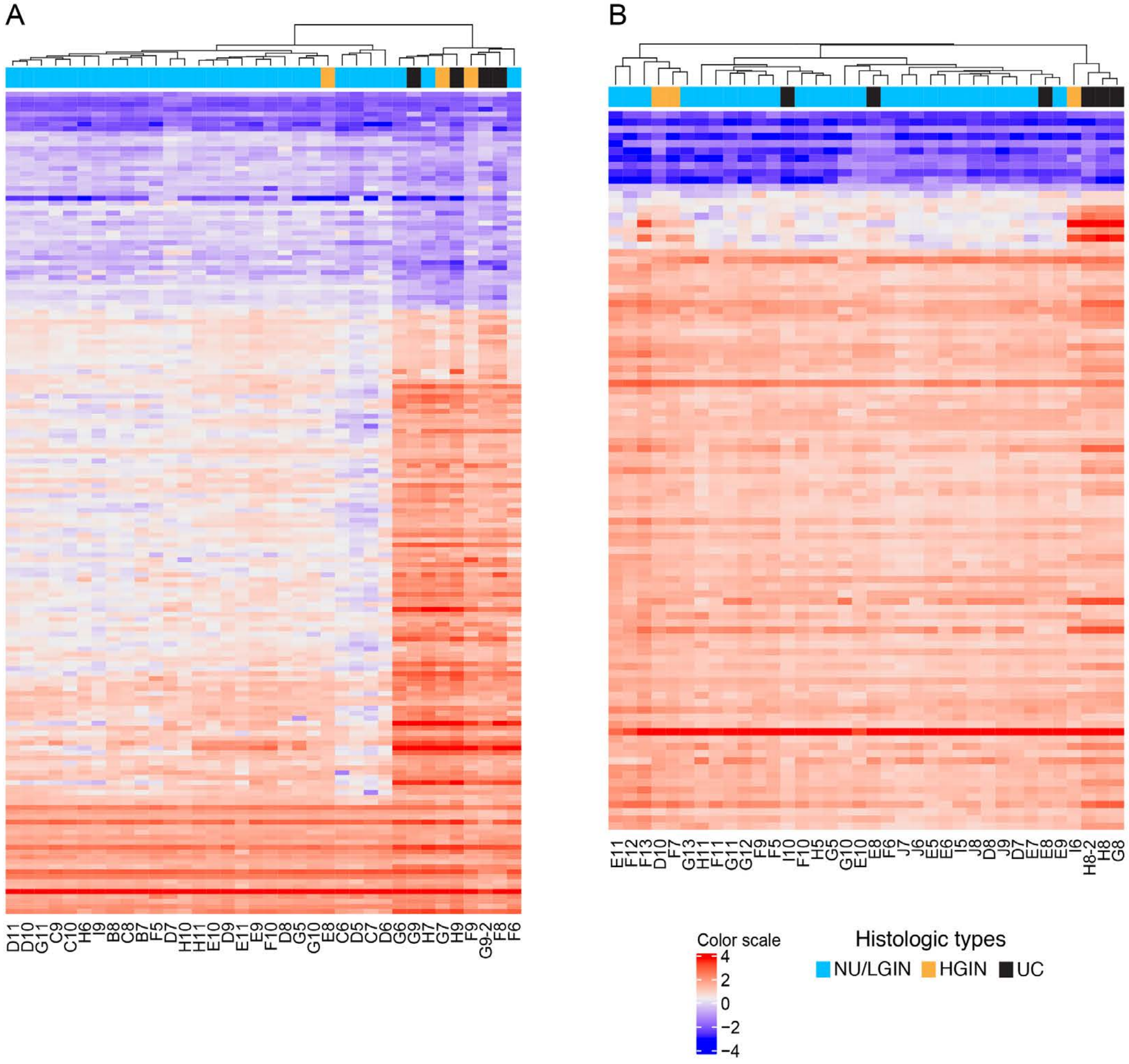

### Extended Data Fig. 2

#### Extended Data Fig. 2

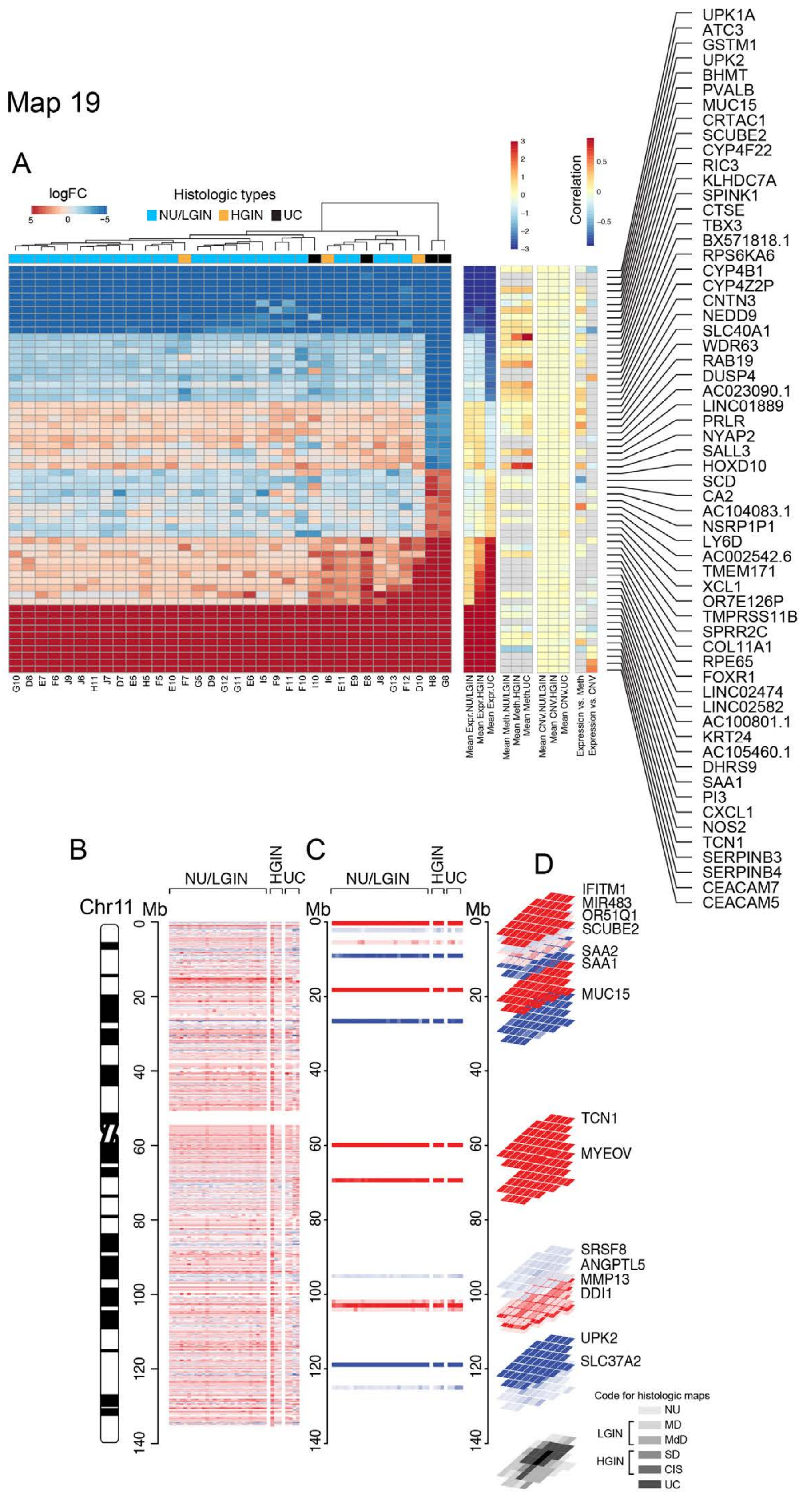

### Extended Data Fig. 3

#### Extended Data Fig. 3

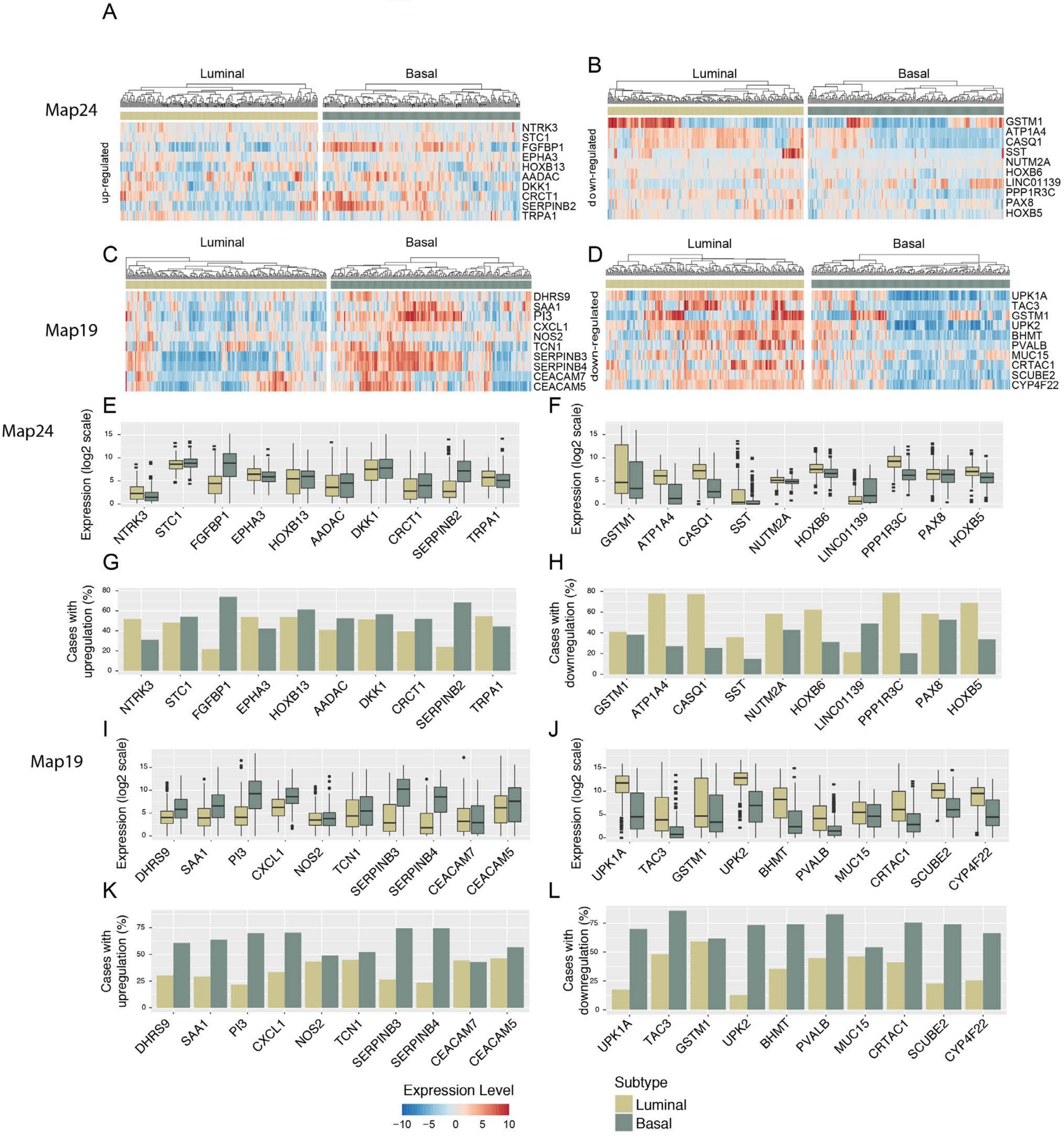

### Extended Data Fig. 4

#### Extended Data Fig. 4

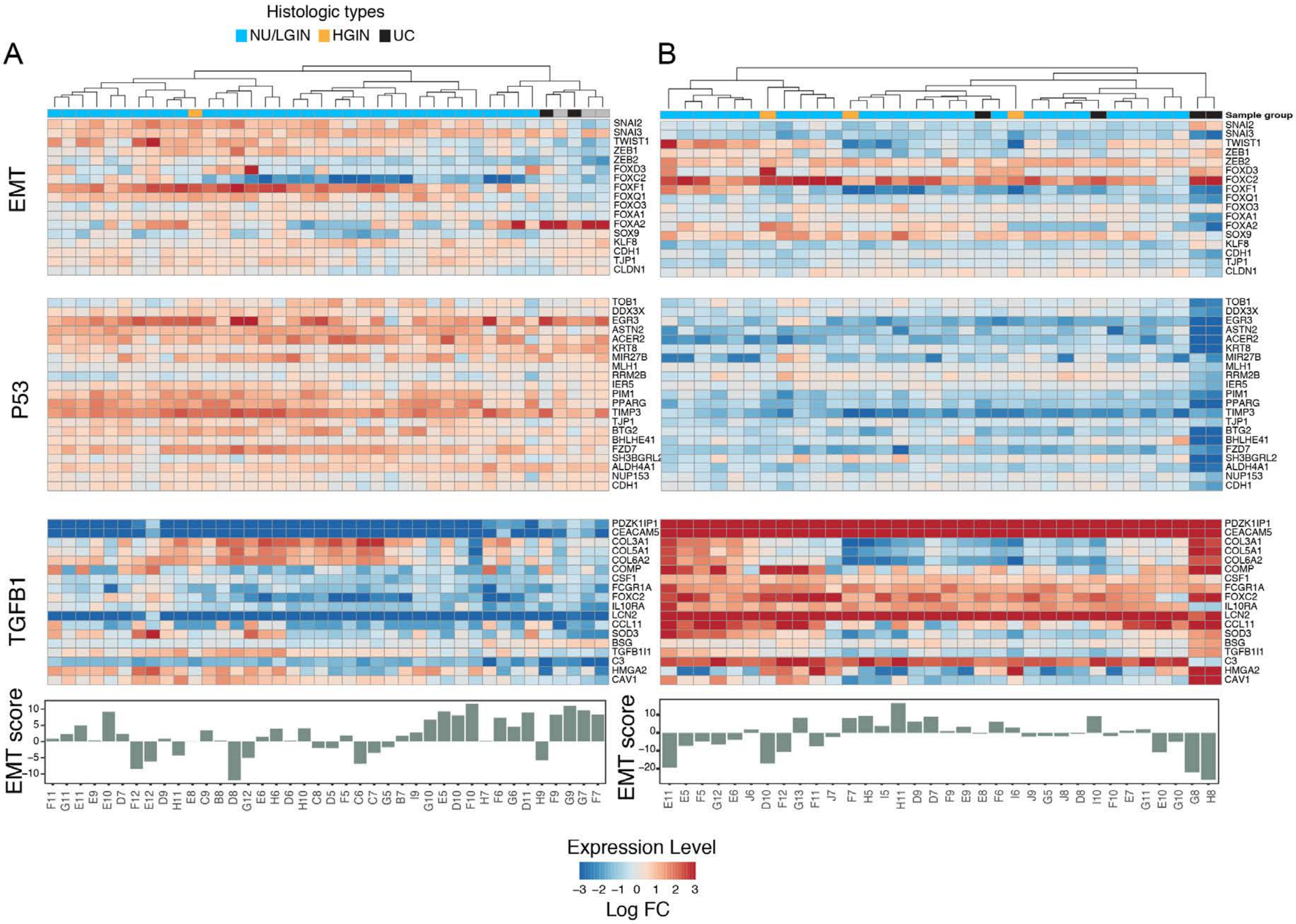

### Extended Data Fig. 5

#### Extended Data Fig. 5

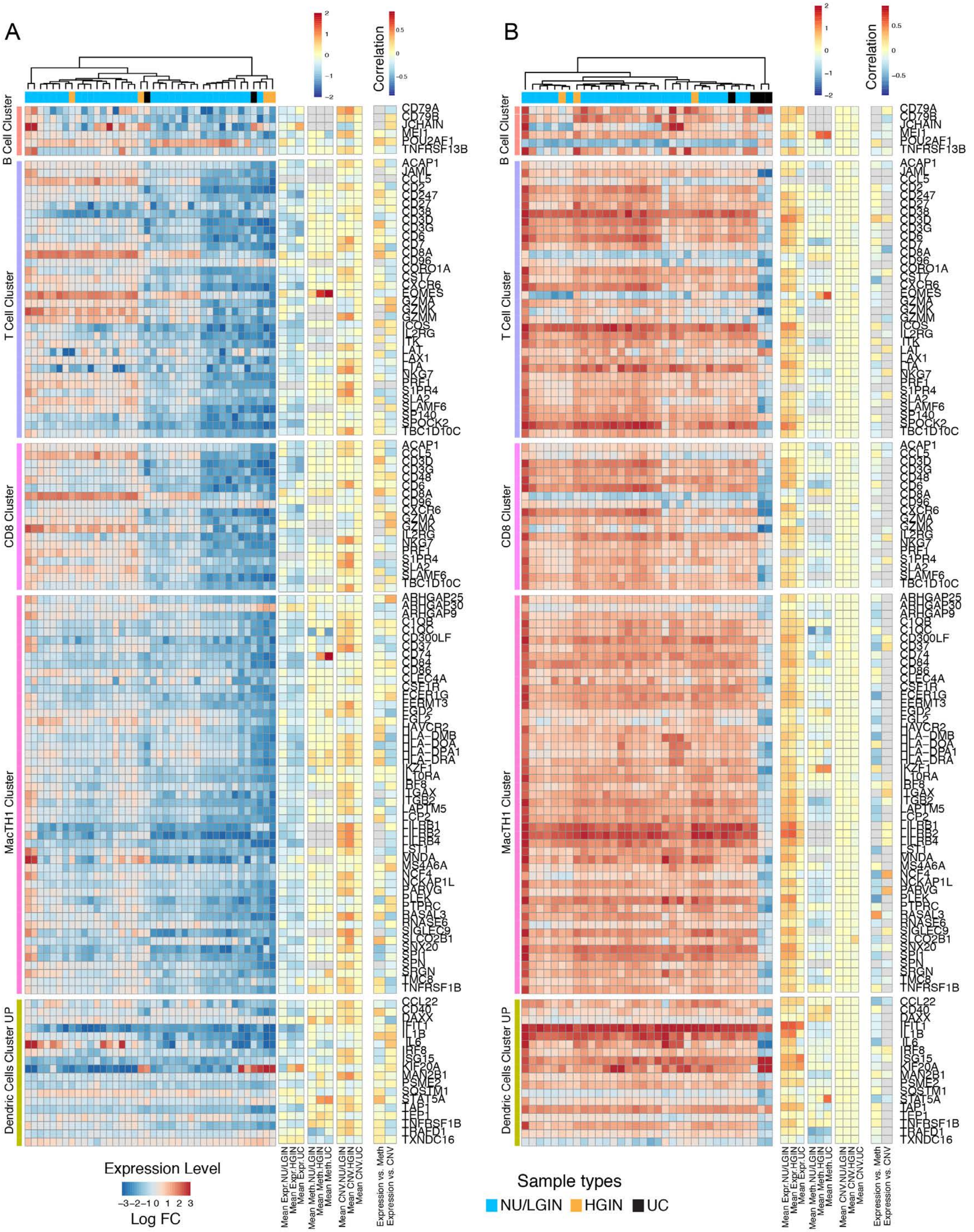

### Extended Data Fig. 6

#### Extended Data Fig. 6

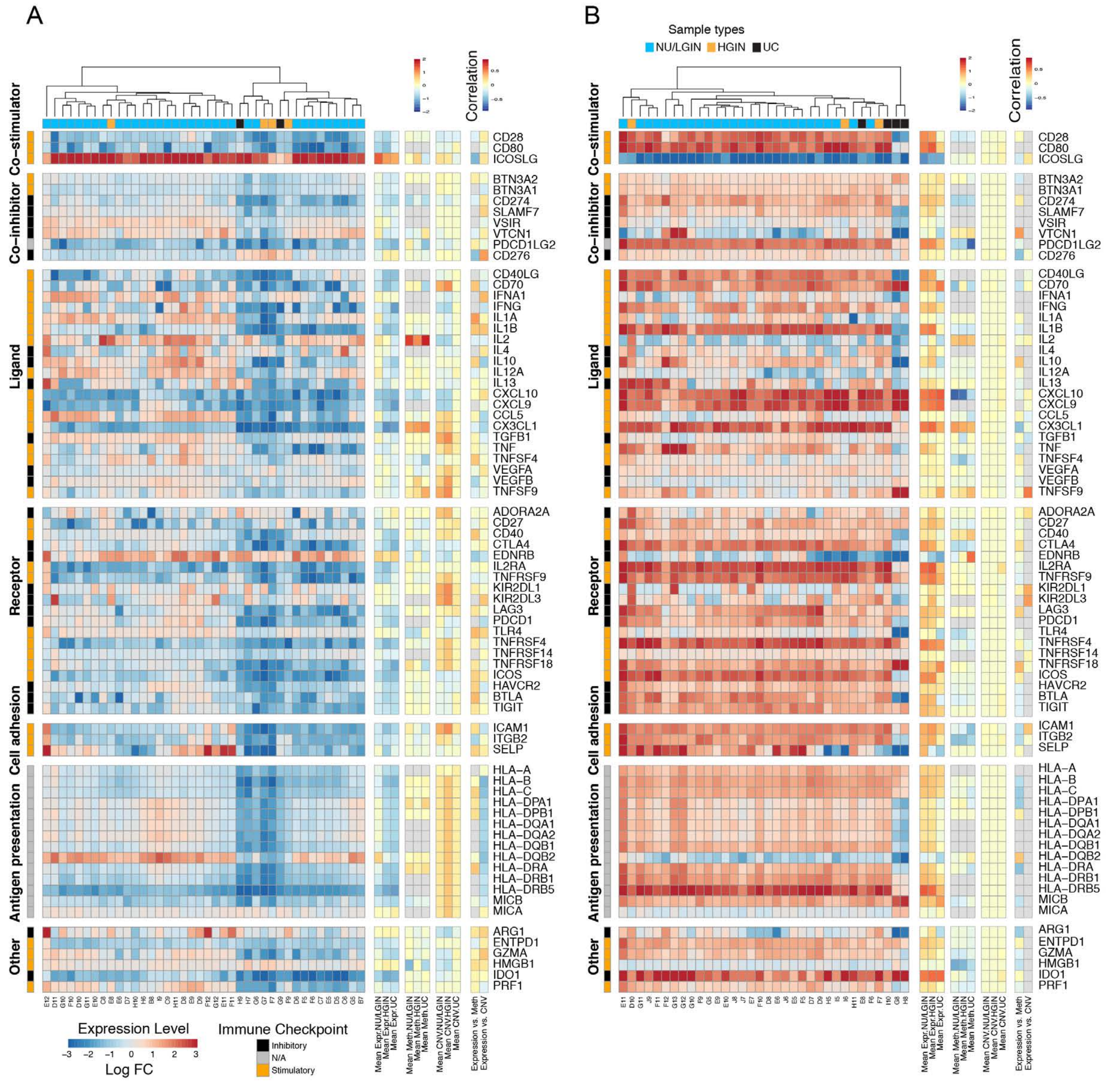

### Extended Data Fig. 7

#### Extended Data Fig. 7

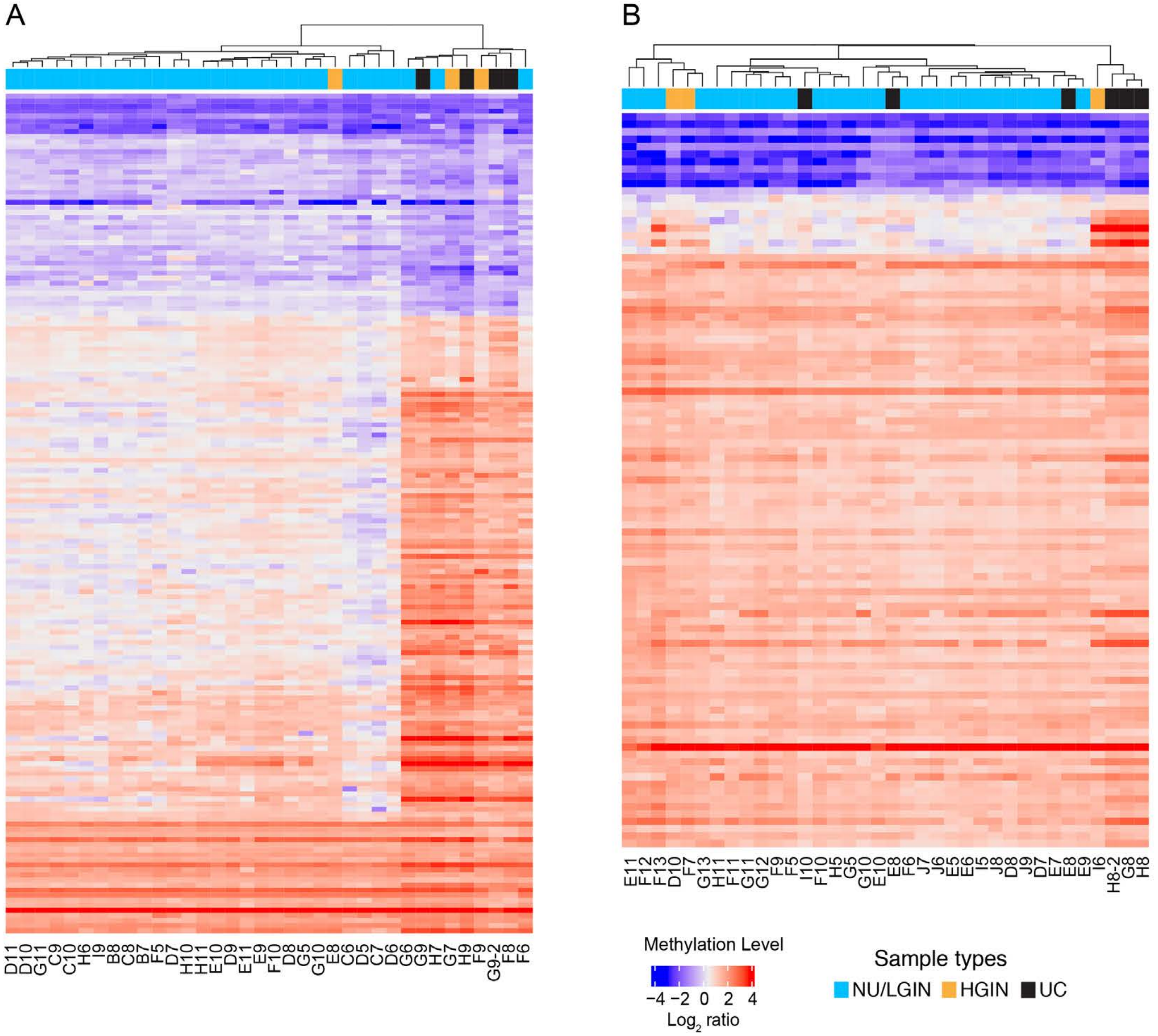

### Extended Data Fig. 8

#### Extended Data Fig. 8

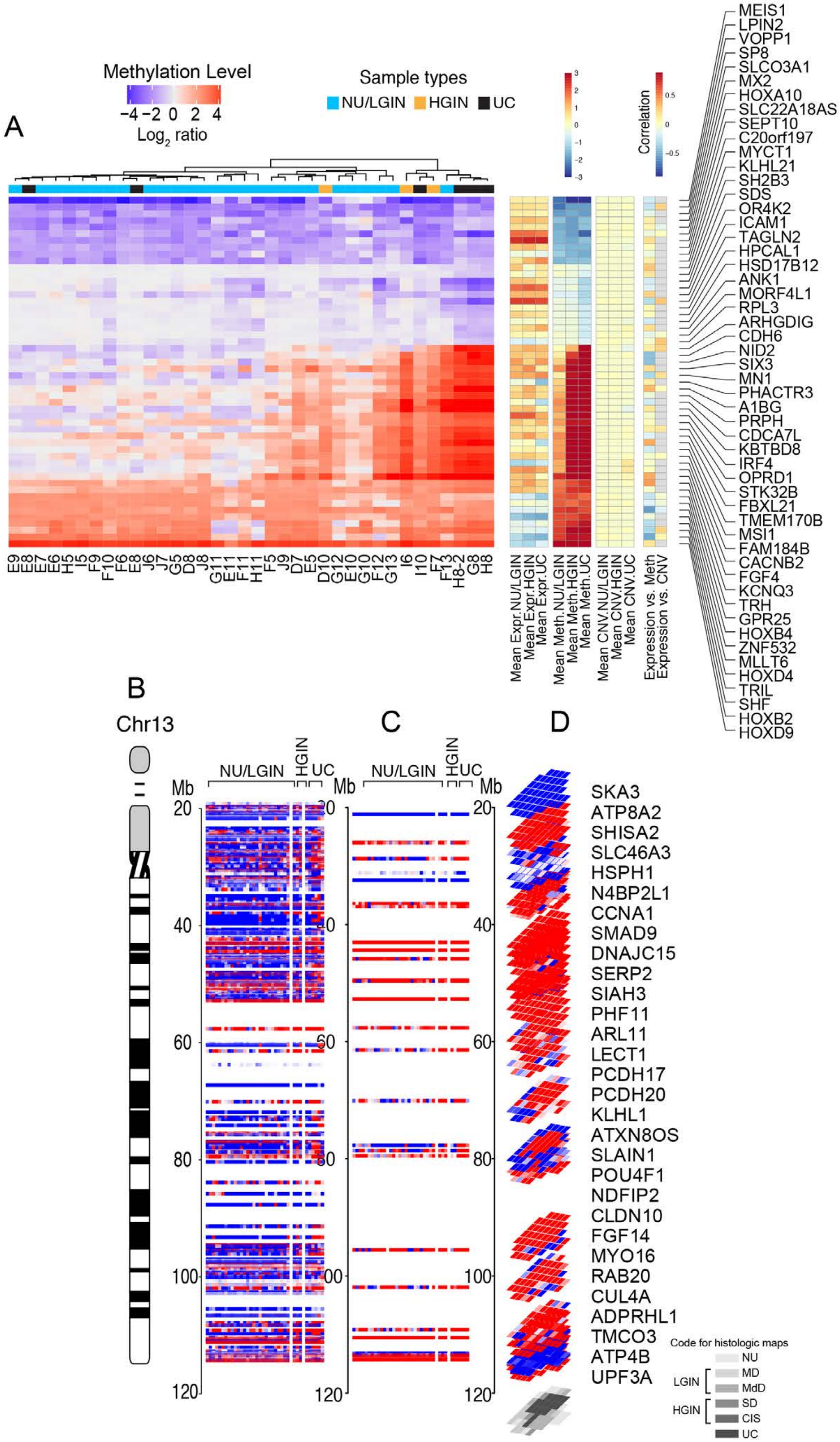

### Extended Data Fig. 9

#### Extended Data Fig. 9

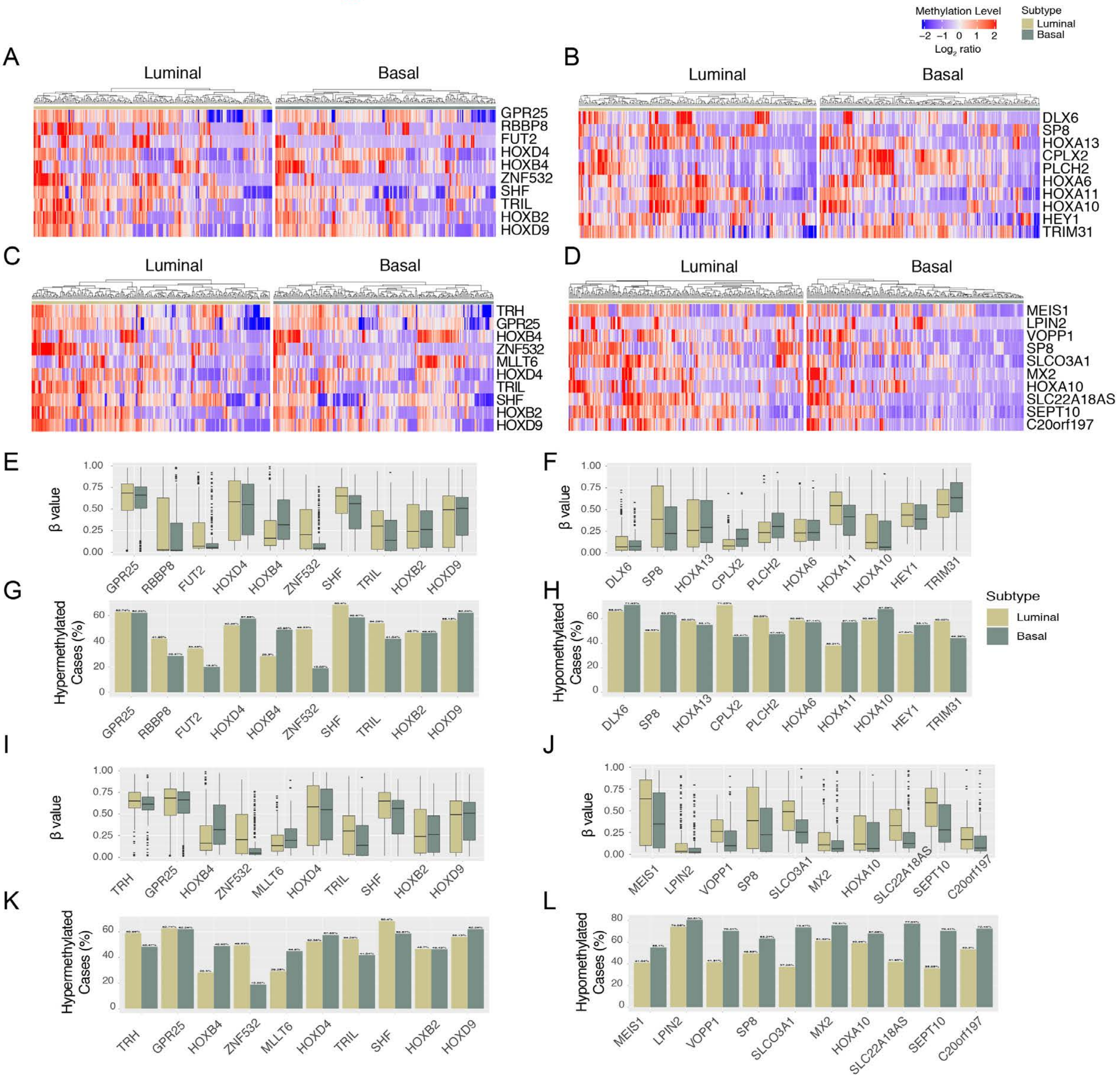

### Extended Data Fig. 10

#### Extended Data Fig. 10

Map19

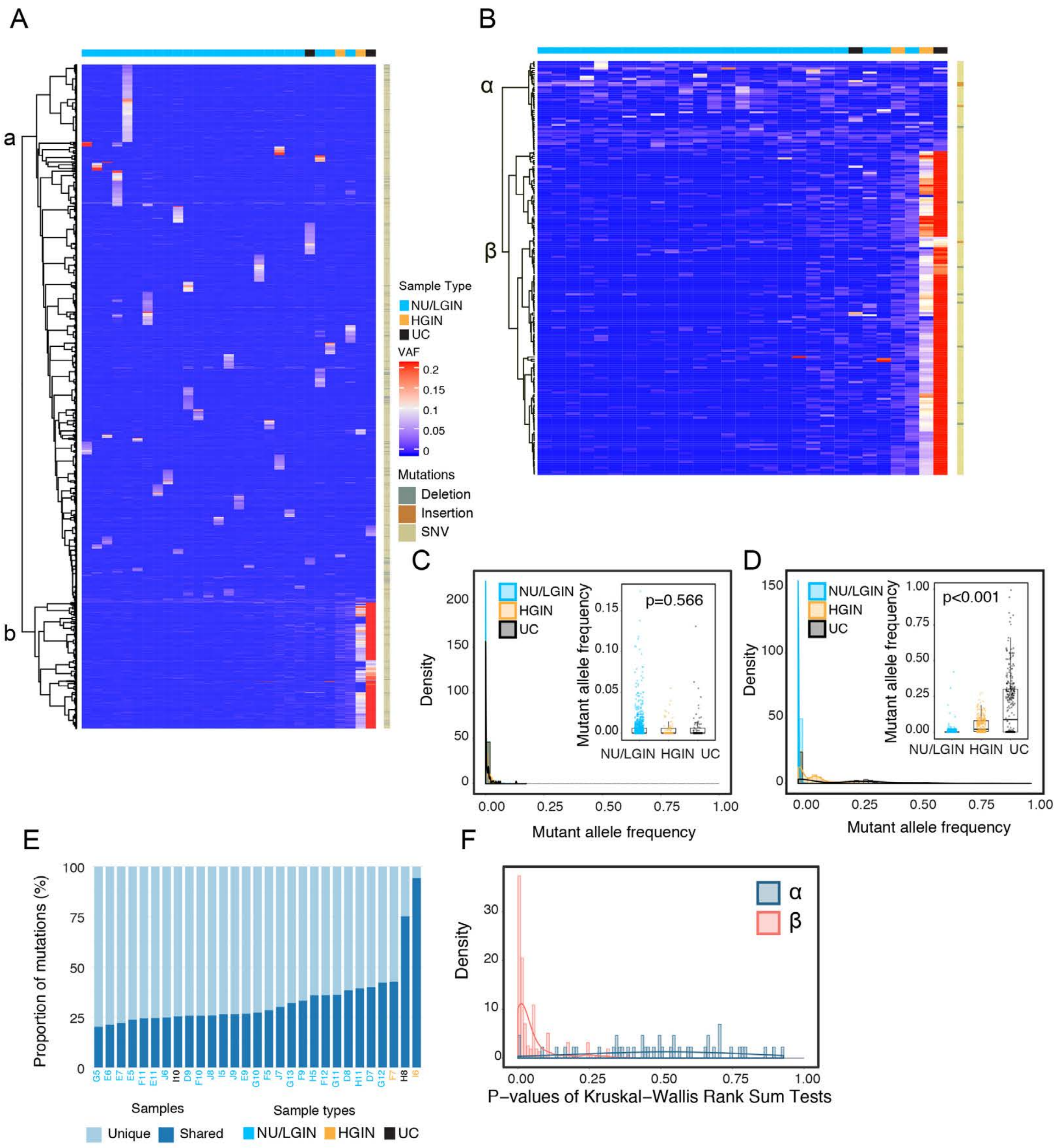

### Extended Data Fig. 11

#### Extended Data Fig. 11

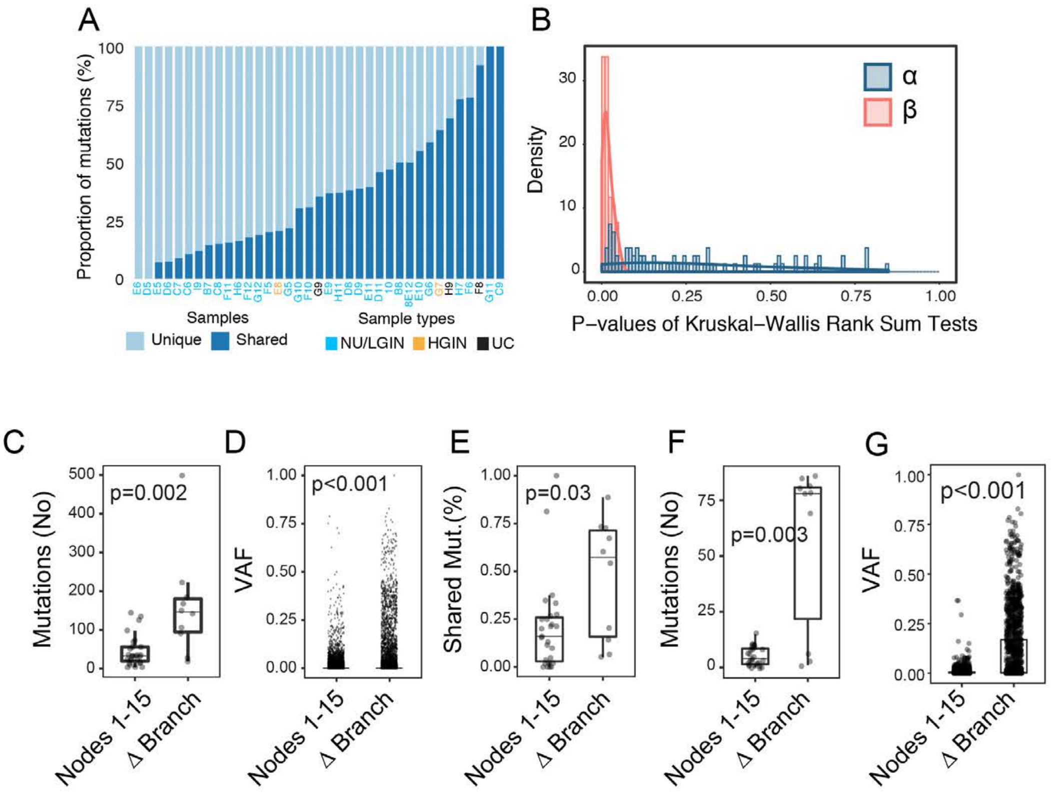

### Extended Data Fig. 12

#### Extended Data Fig. 12

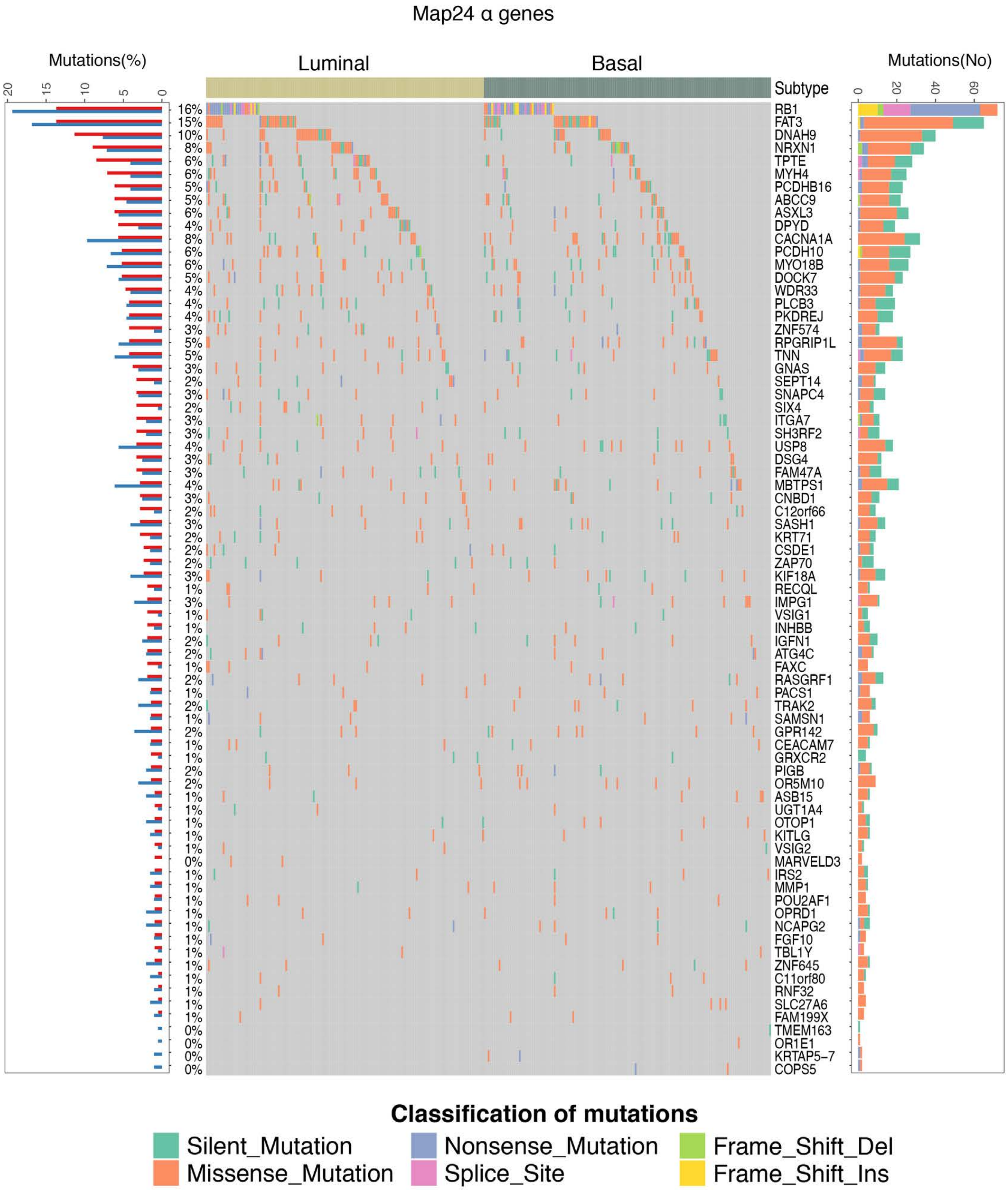

### Extended Data Fig. 13

#### Extended Data Fig. 13

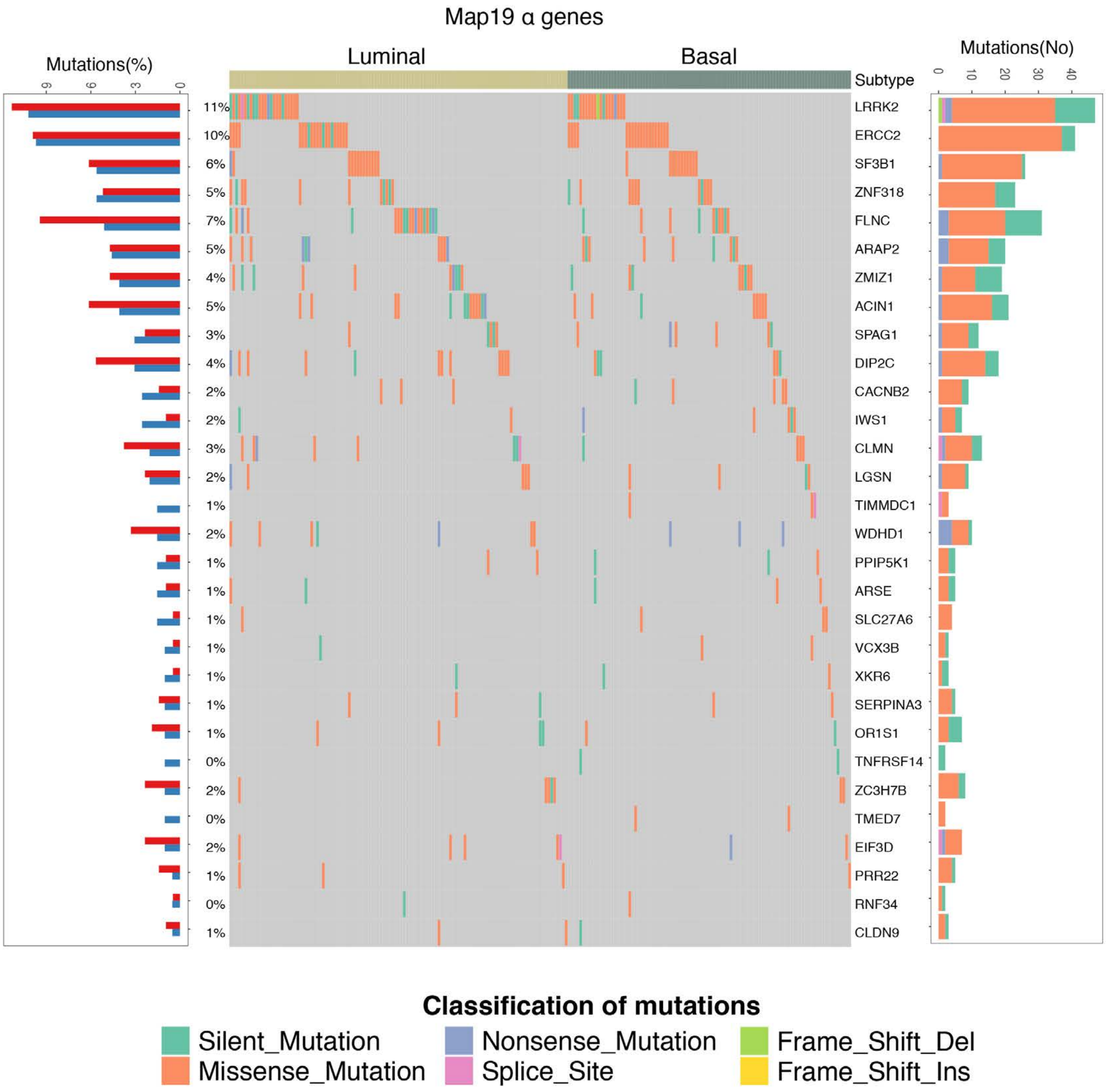

### Extended Data Fig. 14

#### Extended Data Fig. 14

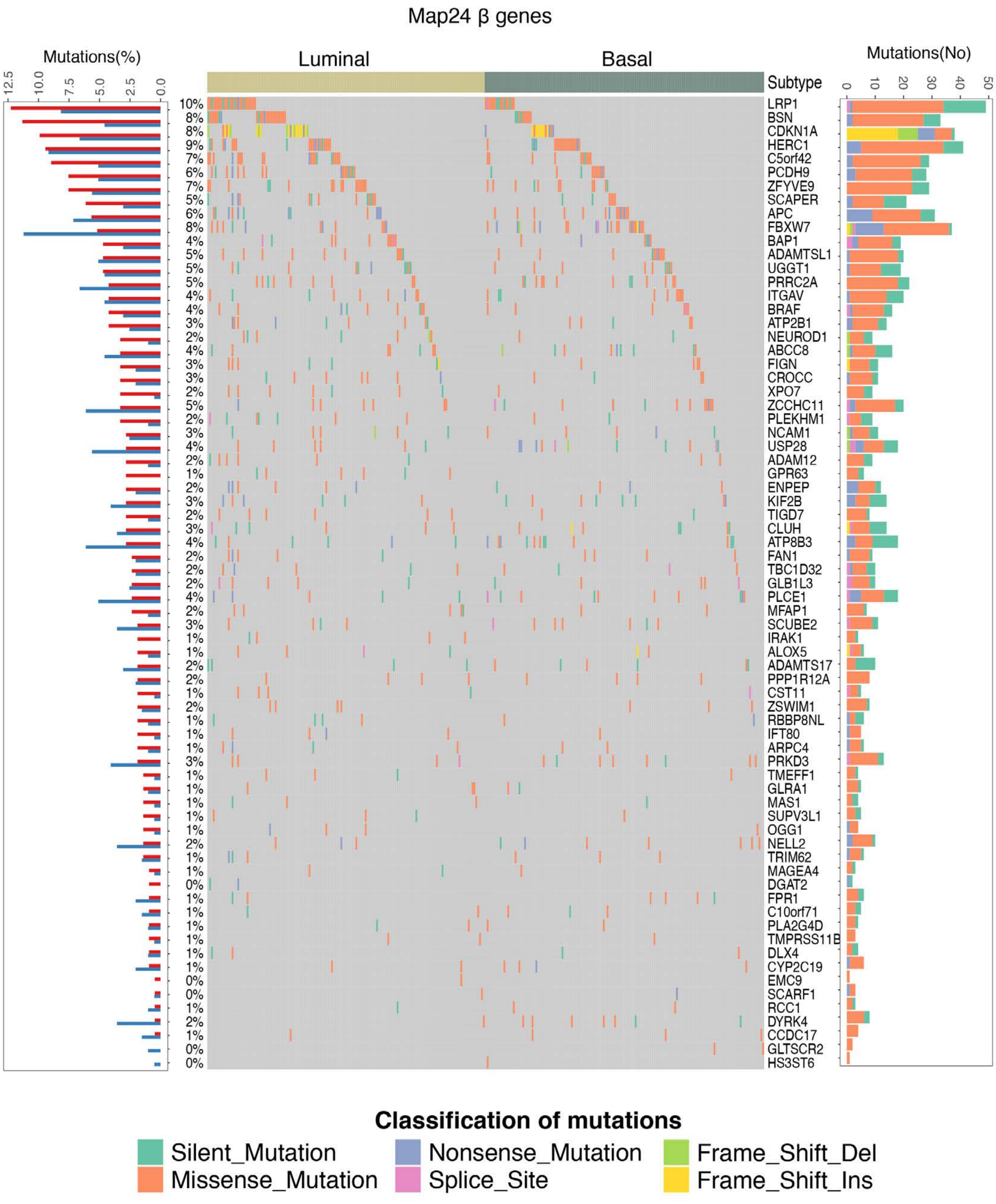

### Extended Data Fig. 15

#### Extended Data Fig. 15

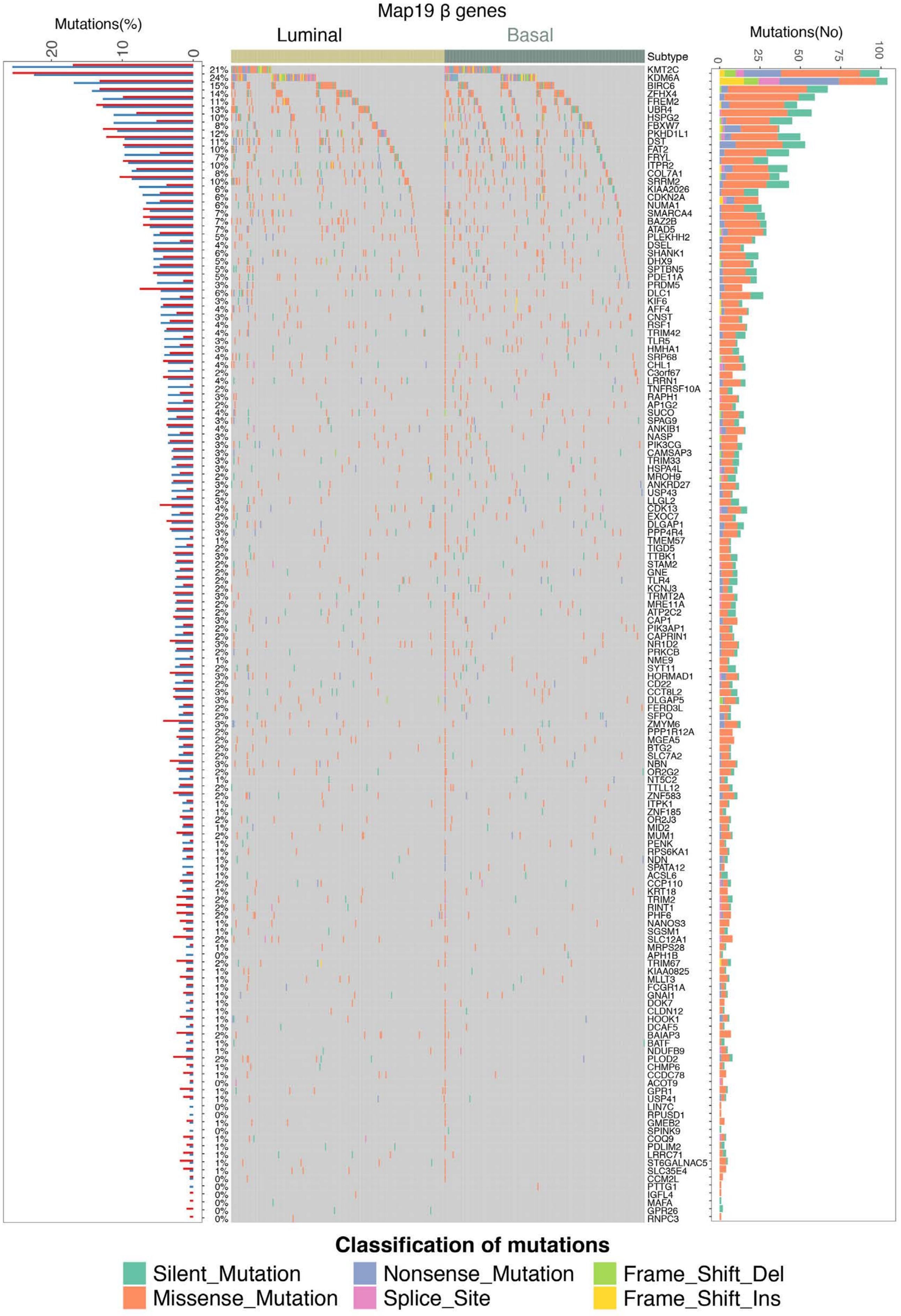

### Extended Data Fig. 16

#### Extended Data Fig. 16

Map 24

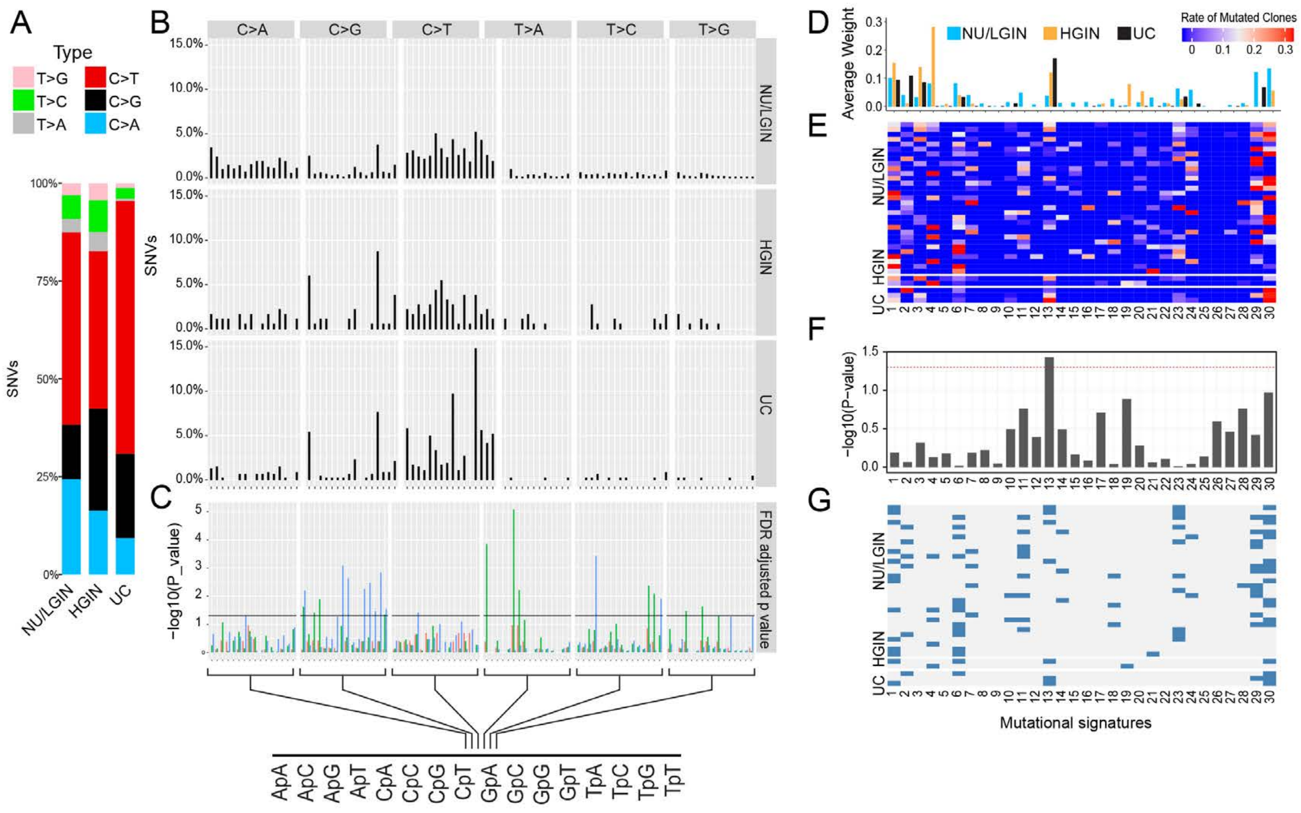

### Extended Data Fig. 17

#### Extended Data Fig. 17

Map 19

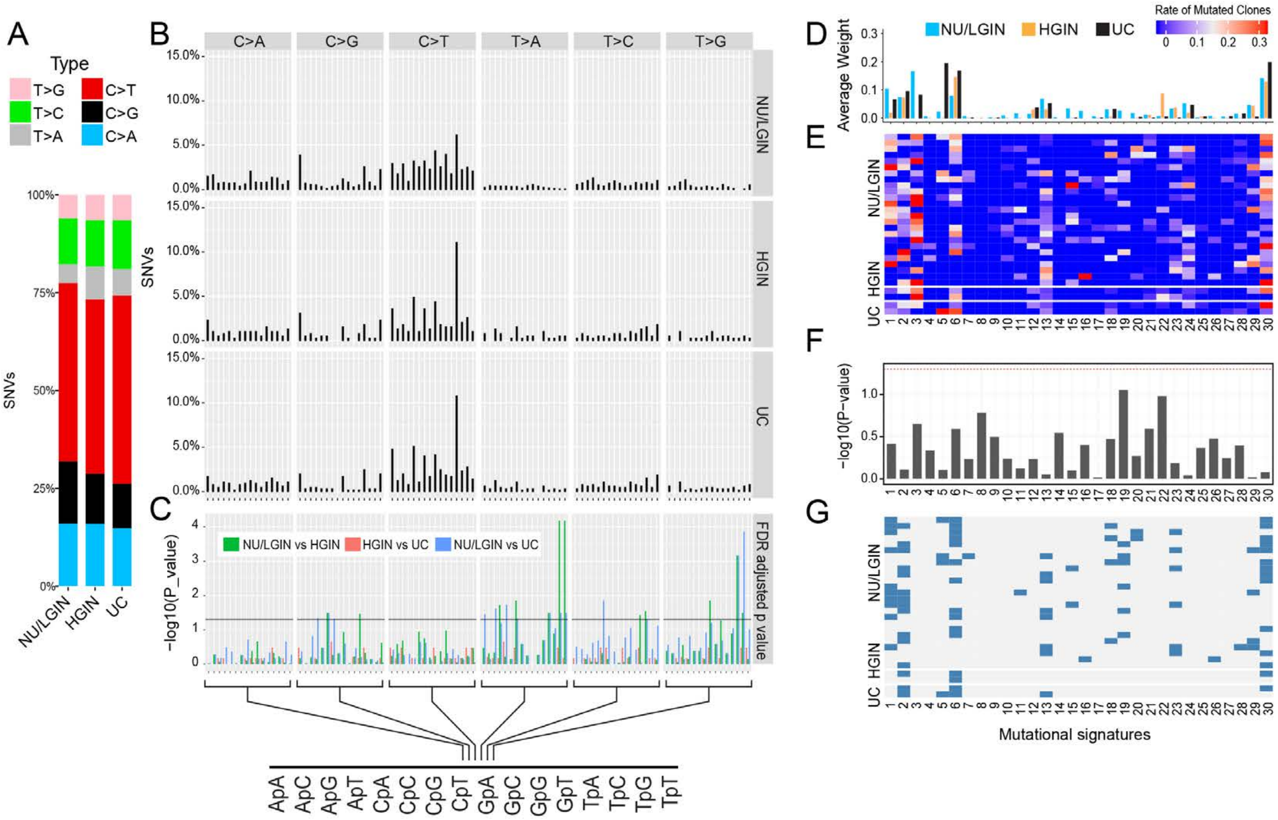

### Extended Data Fig. 18

#### Extended Data Fig. 18

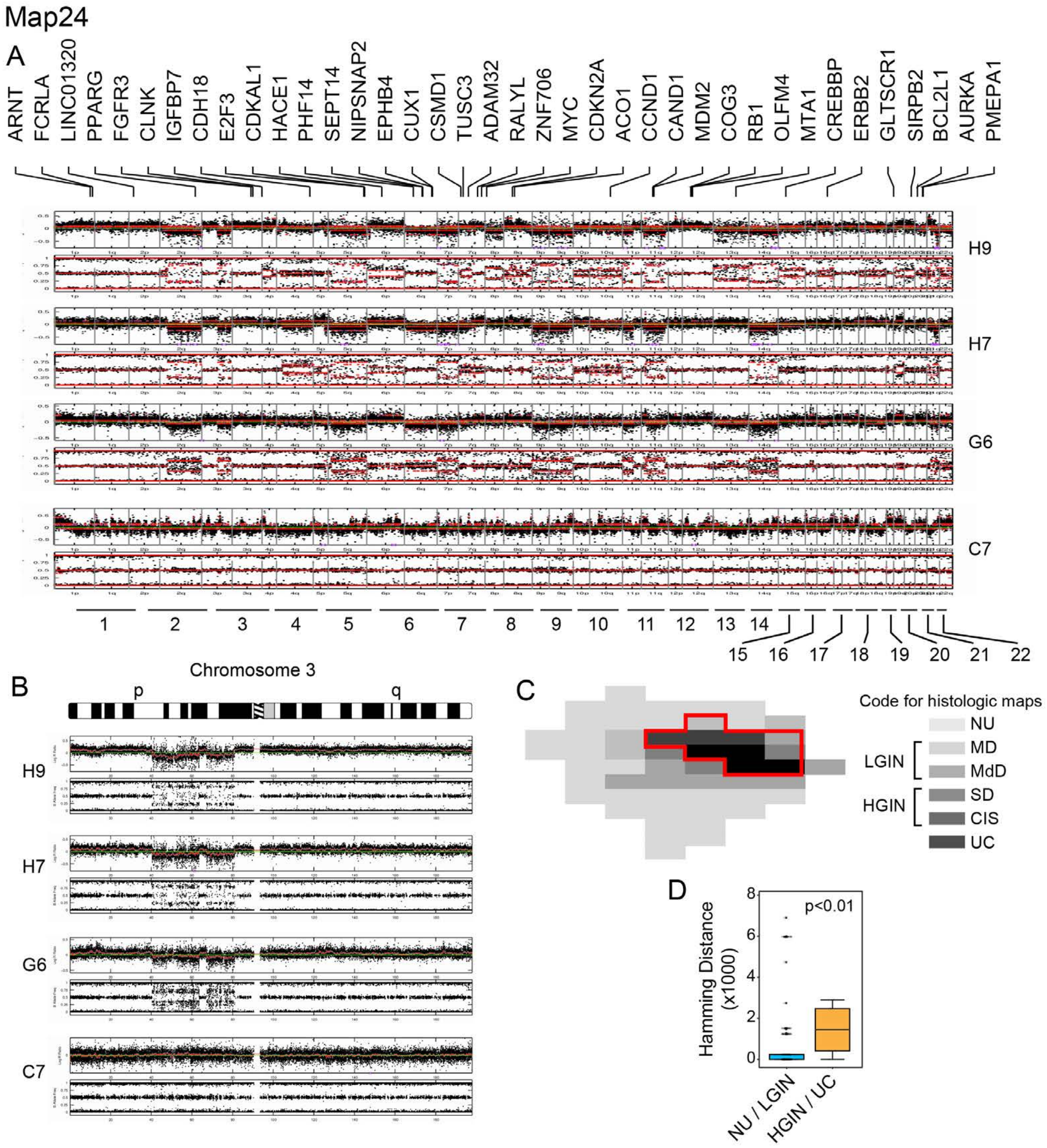

### Extended Data Fig. 19

#### Extended Data Fig. 19

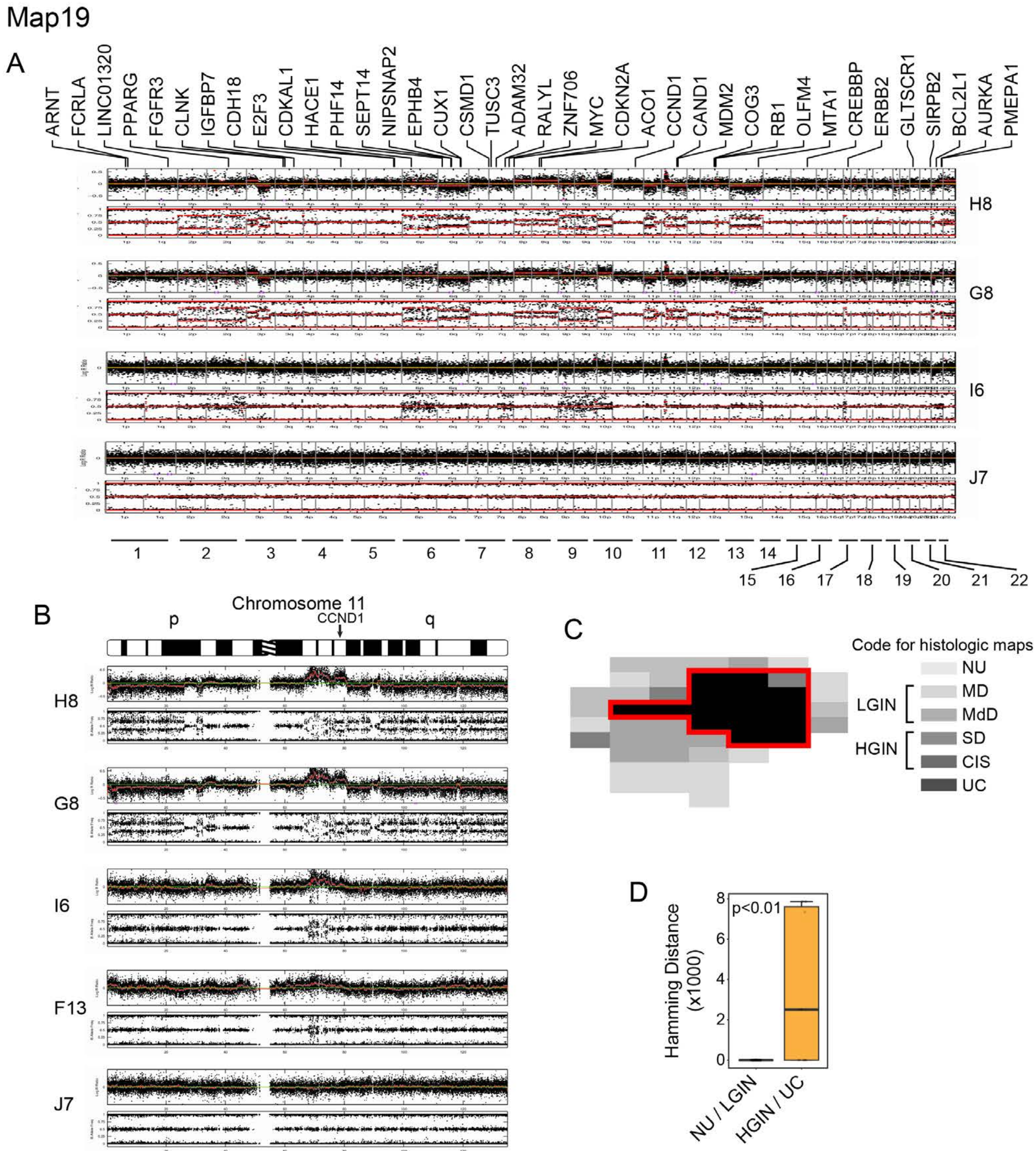

### Extended Data Fig. 20

#### Extended Data Fig. 20

Map19

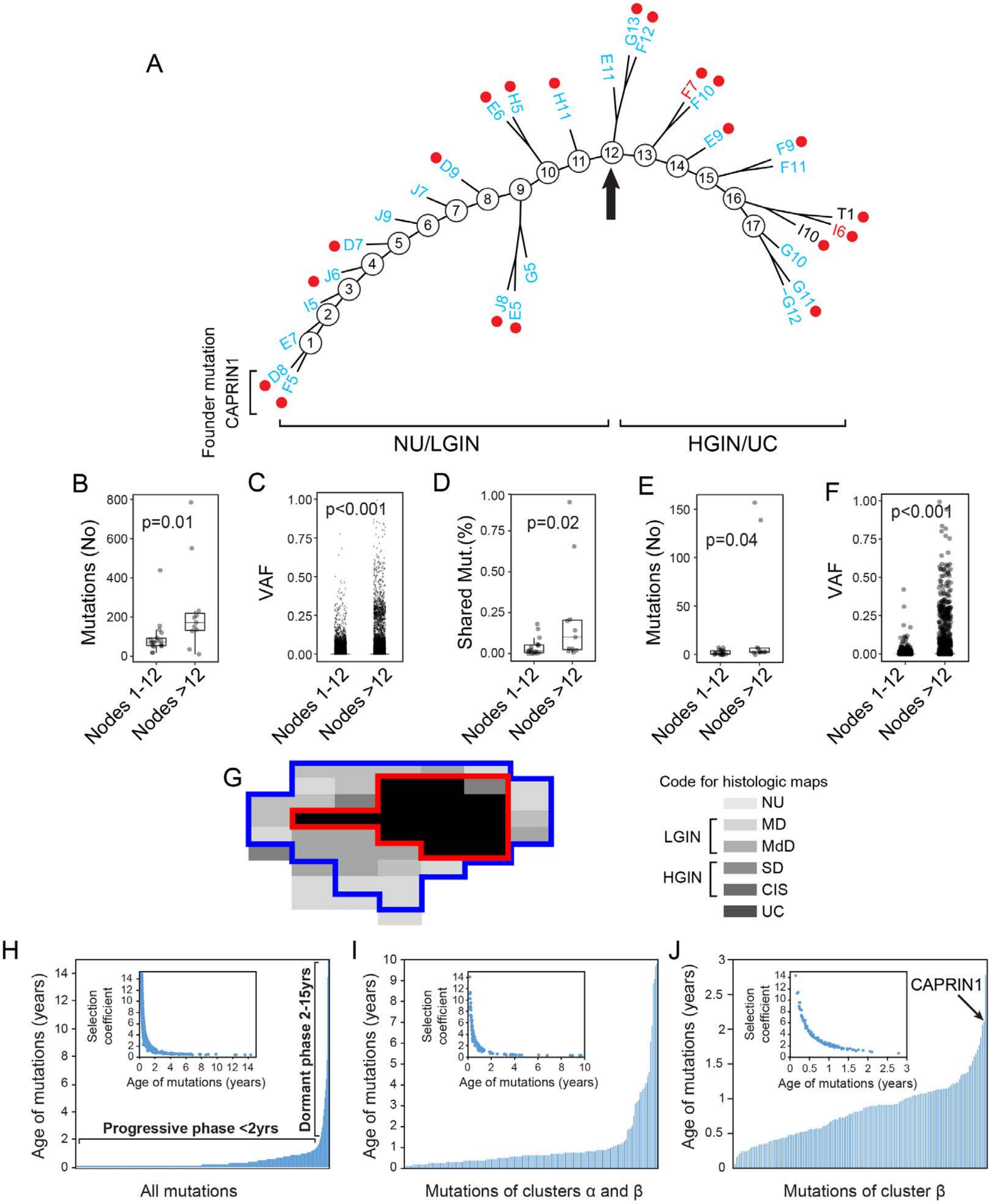

### Extended Data Fig. 21

#### Extended Data Fig. 21

RNA seq

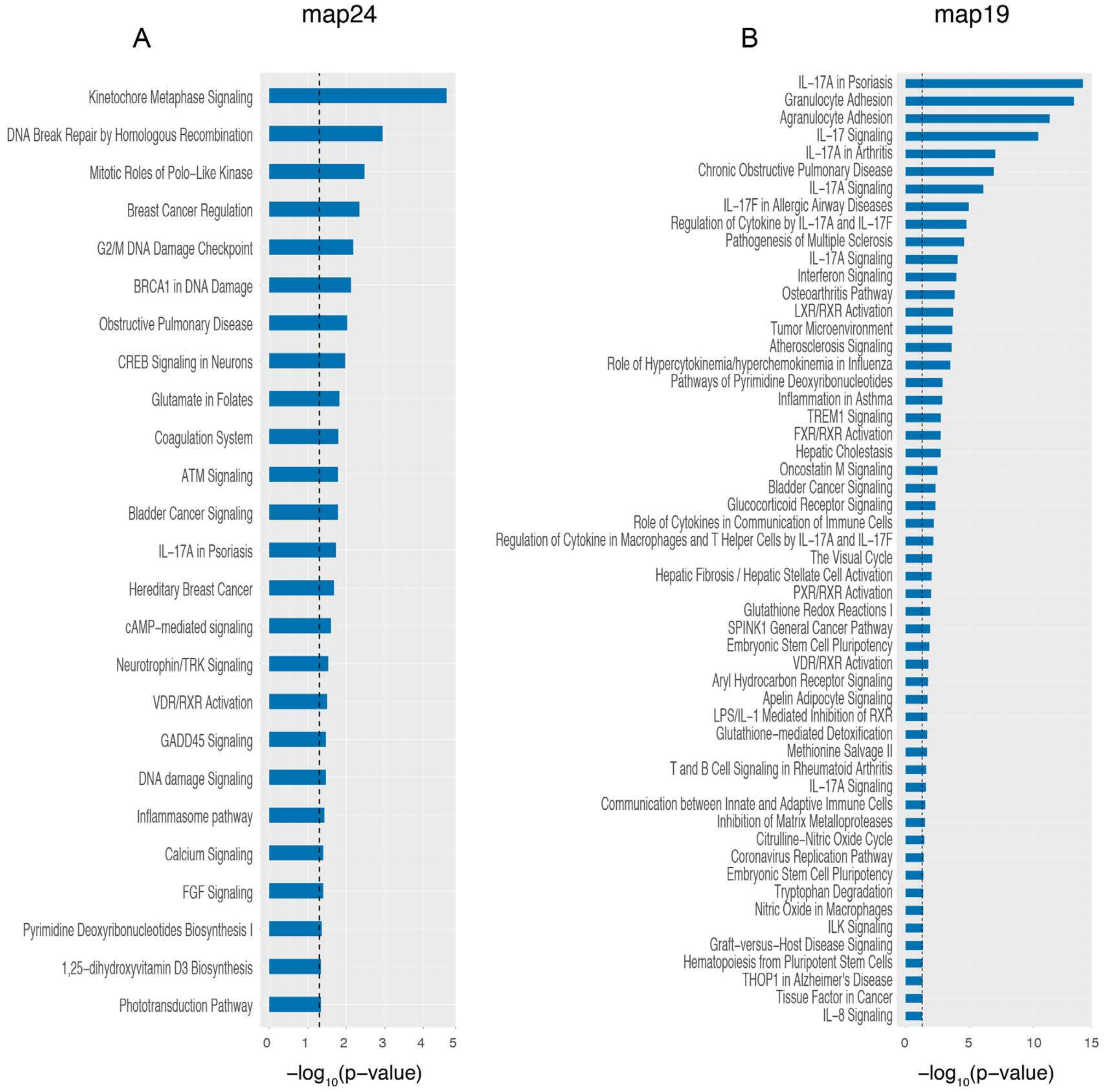

### Extended Data Fig. 22

#### Extended Data Fig. 22

Methylation

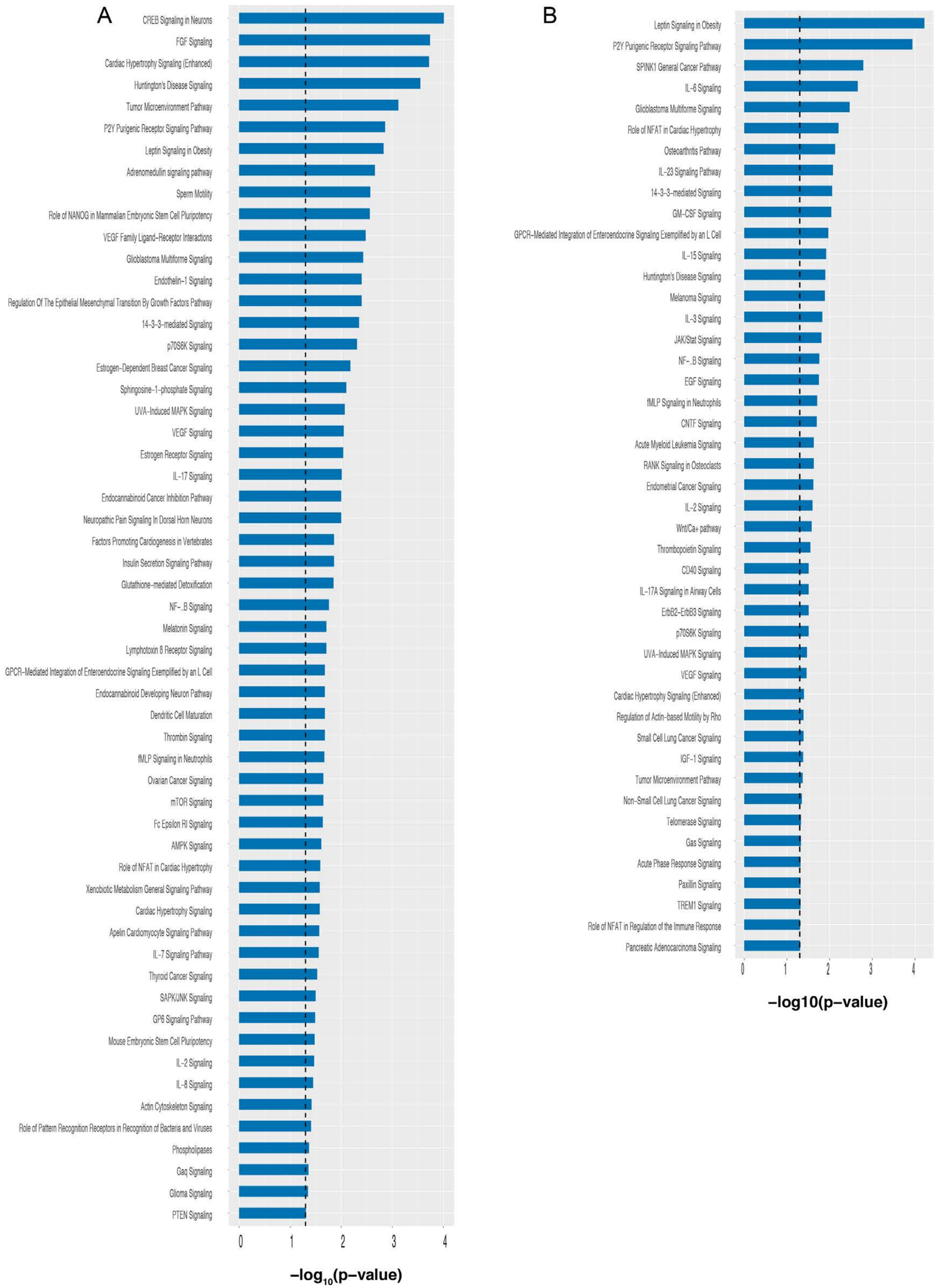

### Extended Data Fig. 23

#### Extended Data Fig. 23

Mutations Alpha and Beta

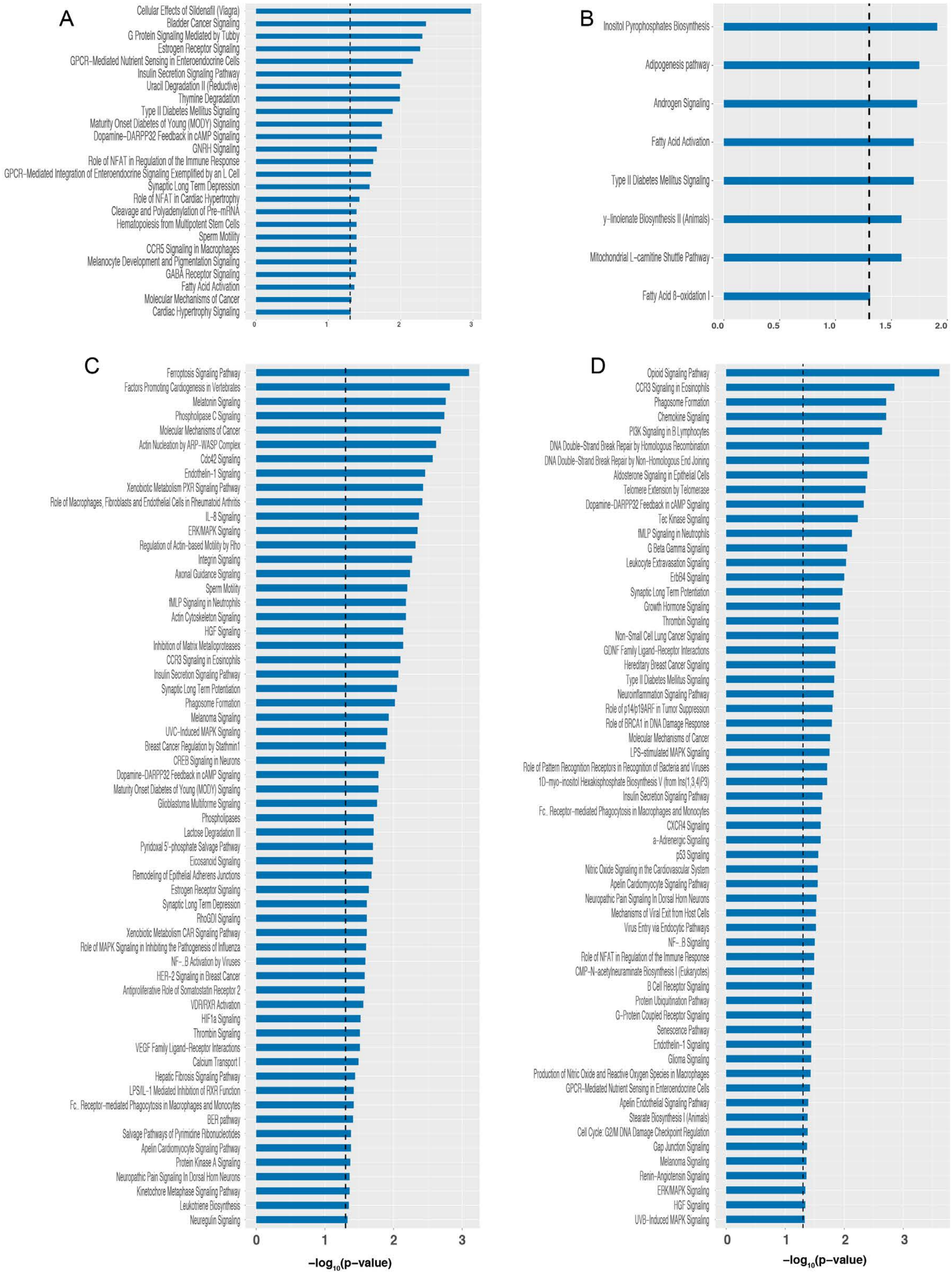
