## Extended Data Table 1 for "The Origin of Bladder Cancer from Mucosal Field Effects"

### Extended Data Table 1. Characterization of Whole-Organ Mucosal Samples from Cystectomy Specimens

| | | | | | | LGIN | | | HGIN | | | UC | |
|---|---|---|---|---|---|---|---|---|---|---|---|---|---|
| | Map # | Age | Origin | Gender | Normal urotheliu m | Mild dysplasia | Moderate dysplasia | N/LGIN | Severe dysplasia | CIS | HGIN | UC | All samples |
| 1 | 16 | 71 | Caucasian | M | 21 | 18 | 5 | 44 | 13 | 7 | 20 | 14 | 78 |
| 2 | 17 | 80 | Caucasian | M | 16 | 5 | 0 | 21 | 0 | 18 | 18 | 0 | 39 |
| 3 | 18 | 81 | Caucasian | M | 0 | 1 | 4 | 5 | 2 | 18 | 20 | 30 | 55 |
| 4 | 19 | 86 | Caucasian | M | 13 | 9 | 9 | 31 | 3 | 0 | 3 | 9 | 43 |
| 5 | 20 | 55 | Caucasian | M | 14 | 13 | 11 | 38 | 2 | 0 | 2 | 2 | 42 |
| 6 | 21 | 69 | Caucasian | M | 7 | 9 | 13 | 29 | 1 | 0 | 1 | 15 | 45 |
| 7 | 22 | 73 | Caucasian | M | 0 | 3 | 6 | 9 | 5 | 13 | 18 | 7 | 34 |
| 8 | 23 | 79 | Caucasian | M | 8 | 7 | 7 | 22 | 1 | 0 | 1 | 18 | 41 |
| 9 | 24 | 68 | Caucasian | M | 32 | 4 | 8 | 44 | 3 | 3 | 6 | 4 | 54 |
| **Total** | | | | | 111 | 69 | 63 | 243 | 30 | 59 | 89 | 99 | 431 |
