## Extended Data Table 2 for "The Origin of Bladder Cancer from Mucosal Field Effects"

**Extended Data Table 2. Summary of Mutations Identified in Luminal and Basal Cystectomy Specimens**

| | Luminal (MAP24) | Basal (MAP19) |
|---|---|---|
| **Cluster A** | | |
| Non-silent mutations | 1303 | 2176 |
| SNV | 1250 | 1924 |
| Insertions | 10 | 29 |
| Deletions | 43 | 223 |
| **Cluster B** | | |
| Non-silent mutations | 76 | 511 |
| SNV | 74 | 475 |
| Insertions | 1 | 6 |
| Deletions | 1 | 30 |
| **Cluster α** | | |
| Non-silent mutations | 80 | 43 |
| SNV | 77 | 41 |
| Insertions | 1 | 0 |
| Deletions | 2 | 2 |
| **Cluster β** | | |
| Non-silent mutations | 77 | 155 |
| SNV | 75 | 147 |
| Insertions | 1 | 1 |
| Deletions | 1 | 7 |
