## Extended Data Table 3 for "The Origin of Bladder Cancer from Mucosal Field Effects"

**Extended Data Table 3. List of Mutations in Cluster Alpha in the Luminal Map (Map 24)**

| | Mutation | Chr | Start | End | Ref | Alt | Protein_Change | Variant_Classification | Effect | Mut_Type | Mut_ID |
|---|---|---|---|---|---|---|---|---|---|---|---|
| 1 | TMEM163:chr2:134718832 | chr2 | 134718832 | 134718832 | G | A | p.P35L | Missense_Mutation | nonsilent | SNV | TMEM163:chr2:134718832 |
| 2 | PACS1:chr11:66070590 | chr11 | 66070590 | 66070590 | A | C | p.Q35P | Missense_Mutation | nonsilent | SNV | PACS1:chr11:66070590 |
| 3 | ASB15:chr7:123617707 | chr7 | 123617707 | 123617707 | A | C | p.N141H | Missense_Mutation | nonsilent | SNV | ASB15:chr7:123617707 |
| 4 | ASB15:chr7:123617682 | chr7 | 123617682 | 123617682 | A | C | p.E132D | Missense_Mutation | nonsilent | SNV | ASB15:chr7:123617682 |
| 5 | PCDH10:chr4:133152499 | chr4 | 133152499 | 133152499 | A | C | p.I787L | Missense_Mutation | nonsilent | SNV | PCDH10:chr4:133152499 |
| 6 | WDR33:chr2:127726703 | chr2 | 127726703 | 127726703 | G | A | p.Q267* | Nonsense_Mutation | null | SNV | WDR33:chr2:127726703 |
| 7 | MBTPS1:chr16:84091800 | chr16 | 84091800 | 84091800 | T | G | p.K299Q | Missense_Mutation | nonsilent | SNV | MBTPS1:chr16:84091800 |
| 8 | UGT1A4:chr2:233718899 | chr2 | 233718899 | 233718899 | G | T | p.E27* | Nonsense_Mutation | null | SNV | UGT1A4:chr2:233718899 |
| 9 | UGT1A4:chr2:233719010 | chr2 | 233719010 | 233719010 | G | A | p.E64K | Missense_Mutation | nonsilent | SNV | UGT1A4:chr2:233719010 |
| 10 | RECQL:chr12:21470988 | chr12 | 21470988 | 21470988 | G | A | p.S593F | Missense_Mutation | nonsilent | SNV | RECQL:chr12:21470988 |
| 11 | OTOP1:chr4:4197744 | chr4 | 4197744 | 4197744 | C | T | p.A364T | Missense_Mutation | nonsilent | SNV | OTOP1:chr4:4197744 |
| 12 | VSIG2:chr11:124750891 | chr11 | 124750891 | 124750891 | G | A | p.P84S | Missense_Mutation | nonsilent | SNV | VSIG2:chr11:124750891 |
| 13 | SEPT14:chr7:55842957 | chr7 | 55842957 | 55842957 | C | A | p.K181N | Missense_Mutation | nonsilent | SNV | SEPT14:chr7:55842957 |
| 14 | MARVELD3:chr16:71626363 | chr16 | 71626363 | 71626363 | G | A | p.R45Q | Missense_Mutation | nonsilent | SNV | MARVELD3:chr16:71626363 |
| 15 | RB1:chr13:48465295 | chr13 | 48465295 | 48465296 | - | T | p.S807fs | Frame_Shift_Ins | null | Insertion | RB1:chr13:48465295 |
| 16 | PKDREJ:chr22:46262713 | chr22 | 46262713 | 46262713 | C | T | p.V204I | Missense_Mutation | nonsilent | SNV | PKDREJ:chr22:46262713 |
| 17 | CSDE1:chr1:114733742 | chr1 | 114733742 | 114733742 | C | T | p.S146N | Missense_Mutation | nonsilent | SNV | CSDE1:chr1:114733742 |
| 18 | PLCB3:chr11:64267452 | chr11 | 64267452 | 64267452 | G | A | p.D1201N | Missense_Mutation | nonsilent | SNV | PLCB3:chr11:64267452 |
| 19 | KITLG:chr12:88518810 | chr12 | 88518810 | 88518810 | G | A | p.L84F | Missense_Mutation | nonsilent | SNV | KITLG:chr12:88518810 |
| 20 | CNBD1:chr8:87353727 | chr8 | 87353727 | 87353727 | G | A | p.C415Y | Missense_Mutation | nonsilent | SNV | CNBD1:chr8:87353727 |
| 21 | IRS2:chr13:109785170 | chr13 | 109785170 | 109785170 | T | A | p.K295M | Missense_Mutation | nonsilent | SNV | IRS2:chr13:109785170 |
| 22 | MMP1:chr11:102795539 | chr11 | 102795539 | 102795539 | T | C | p.I232V | Missense_Mutation | nonsilent | SNV | MMP1:chr11:102795539 |
| 23 | TPTE:chr21:10596087 | chr21 | 10596087 | 10596087 | C | T | p.R426C | Splice_Site | null | SNV | TPTE:chr21:10596087 |
| 24 | SNAPC4:chr9:136381953 | chr9 | 136381953 | 136381953 | G | A | p.R730C | Missense_Mutation | nonsilent | SNV | SNAPC4:chr9:136381953 |
| 25 | PCDHB16:chr5:141184457 | chr5 | 141184457 | 141184457 | G | T | p.R633L | Missense_Mutation | nonsilent | SNV | PCDHB16:chr5:141184457 |
| 26 | TRAK2:chr2:201386338 | chr2 | 201386338 | 201386338 | C | T | p.E615K | Missense_Mutation | nonsilent | SNV | TRAK2:chr2:201386338 |
| 27 | OR1E1:chr17:3397615 | chr17 | 3397615 | 3397615 | T | C | p.S266G | Missense_Mutation | nonsilent | SNV | OR1E1:chr17:3397615 |
| 28 | IMPG1:chr6:75931152 | chr6 | 75931152 | 75931152 | C | A | NA | Splice_Site | null | SNV | IMPG1:chr6:75931152 |
| 29 | VSIG1:chrX:108067099 | chrX | 108067099 | 108067099 | G | A | p.G126D | Missense_Mutation | nonsilent | SNV | VSIG1:chrX:108067099 |
| 30 | SAMSN1:chr21:14498592 | chr21 | 14498592 | 14498592 | C | A | p.E257* | Splice_Site | null | SNV | SAMSN1:chr21:14498592 |
| 31 | GPR142:chr17:74372090 | chr17 | 74372090 | 74372090 | C | G | p.F293L | Missense_Mutation | nonsilent | SNV | GPR142:chr17:74372090 |
| 32 | SIX4:chr14:60720380 | chr14 | 60720380 | 60720380 | T | C | p.H310R | Missense_Mutation | nonsilent | SNV | SIX4:chr14:60720380 |
| 33 | MYH4:chr17:10457566 | chr17 | 10457566 | 10457566 | T | C | p.H584R | Missense_Mutation | nonsilent | SNV | MYH4:chr17:10457566 |
| 34 | KRTAP5-7:chr11:71527659 | chr11 | 71527659 | 71527659 | G | A | p.C120Y | Missense_Mutation | nonsilent | SNV | KRTAP5-7:chr11:71527659 |
| 35 | INHBB:chr2:120349204 | chr2 | 120349204 | 120349204 | T | A | p.L185Q | Missense_Mutation | nonsilent | SNV | INHBB:chr2:120349204 |
| 36 | MUC16:chr19:8885277 | chr19 | 8885277 | 8885277 | C | G | NA | Splice_Site | null | SNV | MUC16:chr19:8885277 |
| 37 | ITGA7:chr12:55694808 | chr12 | 55694808 | 55694808 | G | T | p.H722Q | Missense_Mutation | nonsilent | SNV | ITGA7:chr12:55694808 |
| 38 | ZNF574:chr19:42080798 | chr19 | 42080798 | 42080798 | C | G | p.A731G | Missense_Mutation | nonsilent | SNV | ZNF574:chr19:42080798 |
| 39 | C12orf66:chr12:64194197 | chr12 | 64194197 | 64194197 | T | A | p.H328L | Missense_Mutation | nonsilent | SNV | C12orf66:chr12:64194197 |
| 40 | COPS5:chr8:67059360 | chr8 | 67059360 | 67059360 | C | T | p.V13M | Missense_Mutation | nonsilent | SNV | COPS5:chr8:67059360 |
| 41 | POU2AF1:chr11:111357545 | chr11 | 111357545 | 111357545 | G | A | p.S119L | Missense_Mutation | nonsilent | SNV | POU2AF1:chr11:111357545 |
| 42 | OPRD1:chr1:28862856 | chr1 | 28862856 | 28862856 | T | A | p.V231E | Missense_Mutation | nonsilent | SNV | OPRD1:chr1:28862856 |
| 43 | RPGRIP1L:chr16:53637725 | chr16 | 53637725 | 53637725 | T | C | p.T1030A | Missense_Mutation | nonsilent | SNV | RPGRIP1L:chr16:53637725 |
| 44 | SASH1:chr6:148544104 | chr6 | 148544104 | 148544104 | G | C | p.K878N | Missense_Mutation | nonsilent | SNV | SASH1:chr6:148544104 |
| 45 | IGFN1:chr1:201206758 | chr1 | 201206758 | 201206758 | C | T | p.P622L | Missense_Mutation | nonsilent | SNV | IGFN1:chr1:201206758 |
| 46 | ATG4C:chr1:62834116 | chr1 | 62834116 | 62834116 | G | C | p.D338H | Splice_Site | null | SNV | ATG4C:chr1:62834116 |
| 47 | C11orf80:chr11:66756380 | chr11 | 66756380 | 66756380 | C | T | p.P66L | Missense_Mutation | nonsilent | SNV | C11orf80:chr11:66756380 |
| 48 | MYO18B:chr22:25823622 | chr22 | 25823622 | 25823622 | A | T | p.Q880L | Missense_Mutation | nonsilent | SNV | MYO18B:chr22:25823622 |
| 49 | CEACAM7:chr19:41677414 | chr19 | 41677414 | 41677414 | A | G | p.*266Q | Nonstop_Mutation | null | SNV | CEACAM7:chr19:41677414 |
| 50 | ZAP70:chr2:97732905 | chr2 | 97732905 | 97732918 | GGCACATACGCCCT | - | p.G196fs | Frame_Shift_Del | null | Deletion | ZAP70:chr2:97732905 |
| 51 | SH3RF2:chr5:146013951 | chr5 | 146013951 | 146013951 | G | A | p.G317R | Missense_Mutation | nonsilent | SNV | SH3RF2:chr5:146013951 |

**Extended Data Table 3. List of Mutations in Cluster Alpha in the Luminal Map (Map 24)**

| | Mutation | Chr | Start | End | Ref | Alt | Protein_Change | Variant_Classification | Effect | Mut_Type | Mut_ID |
|---|---|---|---|---|---|---|---|---|---|---|---|
| 52 | DPYD:chr1:97549639 | chr1 | 97549639 | 97549639 | A | G | p.V482A | Missense_Mutation | nonsilent | SNV | DPYD:chr1:97549639 |
| 53 | NCAPG2:chr7:158652293 | chr7 | 158652293 | 158652293 | C | A | p.R978R | Splice_Site | null | SNV | NCAPG2:chr7:158652293 |
| 54 | FGF10:chr5:44388576 | chr5 | 44388576 | 44388576 | G | A | p.T36I | Missense_Mutation | nonsilent | SNV | FGF10:chr5:44388576 |
| 55 | FAT3:chr11:92840649 | chr11 | 92840649 | 92840649 | A | G | p.I3336V | Missense_Mutation | nonsilent | SNV | FAT3:chr11:92840649 |
| 56 | TNN:chr1:175077743 | chr1 | 175077743 | 175077743 | C | A | p.L109M | Missense_Mutation | nonsilent | SNV | TNN:chr1:175077743 |
| 57 | GNAS:chr20:58854895 | chr20 | 58854895 | 58854895 | C | T | p.R544W | Missense_Mutation | nonsilent | SNV | GNAS:chr20:58854895 |
| 58 | USP8:chr15:50495917 | chr15 | 50495917 | 50495917 | G | A | p.A804T | Missense_Mutation | nonsilent | SNV | USP8:chr15:50495917 |
| 59 | GRXCR2:chr5:145866507 | chr5 | 145866507 | 145866507 | A | C | p.Y186* | Nonsense_Mutation | null | SNV | GRXCR2:chr5:145866507 |
| 60 | TBL1Y:chrY:7063918 | chrY | 7063918 | 7063918 | C | T | p.R76C | Missense_Mutation | nonsilent | SNV | TBL1Y:chrY:7063918 |
| 61 | PIGB:chr15:55350844 | chr15 | 55350844 | 55350844 | C | A | p.Y423* | Nonsense_Mutation | null | SNV | PIGB:chr15:55350844 |
| 62 | RNF32:chr7:156675744 | chr7 | 156675744 | 156675744 | G | T | p.E245* | Nonsense_Mutation | null | SNV | RNF32:chr7:156675744 |
| 63 | OR5M10:chr11:56577516 | chr11 | 56577516 | 56577516 | A | G | p.V69A | Missense_Mutation | nonsilent | SNV | OR5M10:chr11:56577516 |
| 64 | ADGB:chr6:146764102 | chr6 | 146764102 | 146764102 | T | C | NA | Splice_Site | null | SNV | ADGB:chr6:146764102 |
| 65 | ZNF645:chrX:22274249 | chrX | 22274249 | 22274249 | A | G | p.R420G | Missense_Mutation | nonsilent | SNV | ZNF645:chrX:22274249 |
| 66 | ABCC9:chr12:21817195 | chr12 | 21817195 | 21817195 | C | T | p.G1295D | Missense_Mutation | nonsilent | SNV | ABCC9:chr12:21817195 |
| 67 | DSG4:chr18:31411348 | chr18 | 31411348 | 31411348 | C | G | p.A752G | Missense_Mutation | nonsilent | SNV | DSG4:chr18:31411348 |
| 68 | SLC27A6:chr5:129016036 | chr5 | 129016036 | 129016036 | C | T | p.T374I | Missense_Mutation | nonsilent | SNV | SLC27A6:chr5:129016036 |
| 69 | FAM199X:chrX:104188281 | chrX | 104188281 | 104188281 | C | A | p.A324E | Missense_Mutation | nonsilent | SNV | FAM199X:chrX:104188281 |
| 70 | ASXL3:chr18:33740213 | chr18 | 33740213 | 33740213 | T | A | p.S937T | Missense_Mutation | nonsilent | SNV | ASXL3:chr18:33740213 |
| 71 | KRT71:chr12:52548701 | chr12 | 52548701 | 52548701 | G | A | p.A271A | Splice_Site | null | SNV | KRT71:chr12:52548701 |
| 72 | DOCK7:chr1:62578916 | chr1 | 62578916 | 62578916 | G | T | p.T641N | Missense_Mutation | nonsilent | SNV | DOCK7:chr1:62578916 |
| 73 | CACNA1A:chr19:13208878 | chr19 | 13208878 | 13208880 | GAT | - | p.H2219del | In_Frame_Del | null | Deletion | CACNA1A:chr19:13208878 |
| 74 | MUC6:chr11:1016770 | chr11 | 1016770 | 1016770 | G | A | p.P2011S | Missense_Mutation | nonsilent | SNV | MUC6:chr11:1016770 |
| 75 | KIF18A:chr11:28088712 | chr11 | 28088712 | 28088712 | G | A | p.R237* | Nonsense_Mutation | null | SNV | KIF18A:chr11:28088712 |
| 76 | NRXN1:chr2:50053527 | chr2 | 50053527 | 50053527 | T | G | p.Q1331P | Missense_Mutation | nonsilent | SNV | NRXN1:chr2:50053527 |
| 77 | FAM47A:chrX:34130558 | chrX | 34130558 | 34130558 | T | A | p.K574M | Missense_Mutation | nonsilent | SNV | FAM47A:chrX:34130558 |
| 78 | FAXC:chr6:99323536 | chr6 | 99323536 | 99323536 | C | T | p.R244H | Missense_Mutation | nonsilent | SNV | FAXC:chr6:99323536 |
| 79 | DNAH9:chr17:11854013 | chr17 | 11854013 | 11854013 | C | A | p.T3173K | Missense_Mutation | nonsilent | SNV | DNAH9:chr17:11854013 |
| 80 | RASGRF1:chr15:79058468 | chr15 | 79058468 | 79058468 | C | T | p.A133T | Missense_Mutation | nonsilent | SNV | RASGRF1:chr15:79058468 |
