## Extended Data Table 4 for "The Origin of Bladder Cancer from Mucosal Field Effects"

**Extended Data Table 4. List of Mutations in Cluster Alpha in the Basal Map (Map 19)**

| | Mutation | Chr | Start | End | Ref | Alt | Protein_Change | Variant_Classification | Effect | Mut_Type | Mut_ID |
|---|---|---|---|---|---|---|---|---|---|---|---|
| 1 | FLNC:chr7:128842686 | chr7 | 128842686 | 128842686 | G | T | p.E793* | Nonsense_Mutation | null | SNV | FLNC:chr7:128842686 |
| 2 | ERCC2:chr19:45364429 | chr19 | 45364429 | 45364429 | T | C | p.N238S | Missense_Mutation | nonsilent | SNV | ERCC2:chr19:45364429 |
| 3 | TIMMDC1:chr3:119498789 | chr3 | 119498789 | 119498789 | T | G | p.F19C | Missense_Mutation | nonsilent | SNV | TIMMDC1:chr3:119498789 |
| 4 | VCX3B:chrX:8466330 | chrX | 8466330 | 8466330 | A | G | p.M230V | Missense_Mutation | nonsilent | SNV | VCX3B:chrX:8466330 |
| 5 | ZMIZ1:chr10:79292307 | chr10 | 79292307 | 79292307 | T | A | p.V303E | Missense_Mutation | nonsilent | SNV | ZMIZ1:chr10:79292307 |
| 6 | CACNB2:chr10:18538203 | chr10 | 18538203 | 18538203 | T | G | p.D414E | Missense_Mutation | nonsilent | SNV | CACNB2:chr10:18538203 |
| 7 | SPAG1:chr8:100220398 | chr8 | 100220398 | 100220398 | G | A | p.C552Y | Missense_Mutation | nonsilent | SNV | SPAG1:chr8:100220398 |
| 8 | PRR22:chr19:5783829 | chr19 | 5783829 | 5783829 | G | C | p.P140A | Missense_Mutation | nonsilent | SNV | PRR22:chr19:5783829 |
| 9 | LRRK2:chr12:40259572 | chr12 | 40259572 | 40259572 | C | T | p.A504V | Missense_Mutation | nonsilent | SNV | LRRK2:chr12:40259572 |
| 10 | IWS1:chr2:127505742 | chr2 | 127505742 | 127505742 | C | A | p.S54I | Missense_Mutation | nonsilent | SNV | IWS1:chr2:127505742 |
| 11 | ACIN1:chr14:23080168 | chr14 | 23080168 | 23080169 | - | G | p.V448fs | Frame_Shift_Ins | null | Insertion | ACIN1:chr14:23080168 |
| 12 | XKR6:chr8:11200724 | chr8 | 11200724 | 11200725 | - | G | p.M206fs | Frame_Shift_Ins | null | Insertion | XKR6:chr8:11200724 |
| 13 | WDHD1:chr14:54944459 | chr14 | 54944459 | 54944459 | C | T | p.G1021E | Missense_Mutation | nonsilent | SNV | WDHD1:chr14:54944459 |
| 14 | CLMN:chr14:95203283 | chr14 | 95203283 | 95203283 | C | T | p.S689N | Missense_Mutation | nonsilent | SNV | CLMN:chr14:95203283 |
| 15 | SERPINA3:chr14:94619419 | chr14 | 94619419 | 94619420 | AT | CA | p.M290Q | Missense_Mutation | nonsilent | SNV | SERPINA3:chr14:94619419 |
| 16 | RP11-51L5.4:chr17:62260772 | chr17 | 62260772 | 62260772 | A | G | NA | Splice_Site | null | SNV | RP11-51L5.4:chr17:62260772 |
| 17 | RP11-402P6.15:chrX:71670304 | chrX | 71670304 | 71670305 | GC | AT | p.Q481* | Nonsense_Mutation | null | SNV | RP11-402P6.15:chrX:71670304 |
| 18 | RP11-402P6.15:chrX:71668128 | chrX | 71668128 | 71668128 | C | A | p.P109T | Missense_Mutation | nonsilent | SNV | RP11-402P6.15:chrX:71668128 |
| 19 | RP11-402P6.15:chrX:71668120 | chrX | 71668120 | 71668120 | C | T | p.A106V | Missense_Mutation | nonsilent | SNV | RP11-402P6.15:chrX:71668120 |
| 20 | PPIP5K1:chr15:43536239 | chr15 | 43536239 | 43536240 | CA | TG | p.C1105H | Missense_Mutation | nonsilent | SNV | PPIP5K1:chr15:43536239 |
| 21 | OR1S1:chr11:58215638 | chr11 | 58215638 | 58215638 | A | G | p.I285M | Missense_Mutation | nonsilent | SNV | OR1S1:chr11:58215638 |
| 22 | OR7E36P:chr13:41431248 | chr13 | 41431248 | 41431249 | - | AAG | NA | Splice_Site | null | Insertion | OR7E36P:chr13:41431248 |
| 23 | OR1S1:chr11:58215047 | chr11 | 58215047 | 58215047 | G | C | p.K88N | Missense_Mutation | nonsilent | SNV | OR1S1:chr11:58215047 |
| 24 | OTOG:chr11:17593745 | chr11 | 17593745 | 17593745 | C | T | p.P1093S | Missense_Mutation | nonsilent | SNV | OTOG:chr11:17593745 |
| 25 | TNFRSF14:chr1:2561679 | chr1 | 2561679 | 2561679 | C | G | p.S186R | Missense_Mutation | nonsilent | SNV | TNFRSF14:chr1:2561679 |
| 26 | KIAA1671:chr22:25177409 | chr22 | 25177409 | 25177409 | G | A | p.R1654H | Missense_Mutation | nonsilent | SNV | KIAA1671:chr22:25177409 |
| 27 | GRIP2:chr3:14540271 | chr3 | 14540271 | 14540271 | G | A | p.A13V | Missense_Mutation | nonsilent | SNV | GRIP2:chr3:14540271 |
| 28 | DIP2C:chr10:399191 | chr10 | 399191 | 399191 | G | A | p.P393L | Missense_Mutation | nonsilent | SNV | DIP2C:chr10:399191 |
| 29 | RNF34:chr12:121416160 | chr12 | 121416160 | 121416160 | C | T | p.A3V | Splice_Site | null | SNV | RNF34:chr12:121416160 |
| 30 | SF3B1:chr2:197400743 | chr2 | 197400743 | 197400743 | A | C | p.L897R | Missense_Mutation | nonsilent | SNV | SF3B1:chr2:197400743 |
| 31 | STKLD1:chr9:133403800 | chr9 | 133403800 | 133403800 | C | A | p.C525* | Nonsense_Mutation | null | SNV | STKLD1:chr9:133403800 |
| 32 | ZNF318:chr6:43368972 | chr6 | 43368972 | 43368979 | TGCCGCTG | - | p.D129fs | Frame_Shift_Del | null | Deletion | ZNF318:chr6:43368972 |
| 33 | ARSE:chrX:2949610 | chrX | 2949610 | 2949610 | C | T | p.R183H | Missense_Mutation | nonsilent | SNV | ARSE:chrX:2949610 |
| 34 | RP11-683L23.1:chr18:48009 | chr18 | 48009 | 48009 | C | T | p.C239Y | Missense_Mutation | nonsilent | SNV | RP11-683L23.1:chr18:48009 |
| 35 | CLDN9:chr16:3013447 | chr16 | 3013447 | 3013447 | C | A | p.L29M | Missense_Mutation | nonsilent | SNV | CLDN9:chr16:3013447 |
| 36 | ZC3H7B:chr22:41320707 | chr22 | 41320707 | 41320707 | T | A | p.F16Y | Missense_Mutation | nonsilent | SNV | ZC3H7B:chr22:41320707 |
| 37 | ARAP2:chr4:36148419 | chr4 | 36148419 | 36148419 | C | A | p.E996* | Nonsense_Mutation | null | SNV | ARAP2:chr4:36148419 |
| 38 | RBMXL3:chrX:115191182 | chrX | 115191182 | 115191183 | CA | TG | p.Q581W | Missense_Mutation | nonsilent | SNV | RBMXL3:chrX:115191182 |
| 39 | RBMXL3:chrX:115191188 | chrX | 115191188 | 115191188 | A | C | p.N583H | Missense_Mutation | nonsilent | SNV | RBMXL3:chrX:115191188 |
| 40 | TMED7:chr5:115625789 | chr5 | 115625789 | 115625789 | G | A | p.P2S | Missense_Mutation | nonsilent | SNV | TMED7:chr5:115625789 |
| 41 | SLC27A6:chr5:129029625 | chr5 | 129029625 | 129029625 | C | A | p.T534K | Missense_Mutation | nonsilent | SNV | SLC27A6:chr5:129029625 |
| 42 | LGSN:chr6:63280565 | chr6 | 63280565 | 63280565 | C | A | p.C329F | Missense_Mutation | nonsilent | SNV | LGSN:chr6:63280565 |
| 43 | EIF3D:chr22:36519493 | chr22 | 36519493 | 36519493 | C | T | p.R208H | Missense_Mutation | nonsilent | SNV | EIF3D:chr22:36519493 |
