## Extended Data Table 5 for "The Origin of Bladder Cancer from Mucosal Field Effects"

**Extended Data Table 5. List of Mutations in Cluster Beta in the Luminal Map (Map 24)**

| | Mutation | Chr | Start | End | Ref | Alt | Protein_Change | Variant_Classification | Effect | Mut_Type | Mut_ID |
|---|---|---|---|---|---|---|---|---|---|---|---|
| 1 | MAGEA4:chrX:151924617 | chrX | 1.52E+08 | 1.52E+08 | G | C | p.*318S | Nonstop_Mutation | null | SNV | MAGEA4:chrX:151924617 |
| 2 | EMC9:chr14:24139634 | chr14 | 24139634 | 24139634 | G | C | p.L12V | Missense_Mutation | nonsilent | SNV | EMC9:chr14:24139634 |
| 3 | SCUBE2:chr11:9047414 | chr11 | 9047414 | 9047415 | TG | - | p.T648fs | Frame_Shift_Del | null | Deletion | SCUBE2:chr11:9047414 |
| 4 | BAP1:chr3:52403766 | chr3 | 52403766 | 52403766 | G | C | p.S460* | Nonsense_Mutation | null | SNV | BAP1:chr3:52403766 |
| 5 | NEUROD1:chr2:181678217 | chr2 | 1.82E+08 | 1.82E+08 | G | C | p.P215R | Missense_Mutation | nonsilent | SNV | NEUROD1:chr2:181678217 |
| 6 | NCAM1:chr11:113255947 | chr11 | 1.13E+08 | 1.13E+08 | C | G | p.I623M | Missense_Mutation | nonsilent | SNV | NCAM1:chr11:113255947 |
| 7 | C5orf42:chr5:37226683 | chr5 | 37226683 | 37226683 | G | T | p.Q638K | Missense_Mutation | nonsilent | SNV | C5orf42:chr5:37226683 |
| 8 | TMEFF1:chr9:100572613 | chr9 | 1.01E+08 | 1.01E+08 | T | C | p.I332T | Missense_Mutation | nonsilent | SNV | TMEFF1:chr9:100572613 |
| 9 | ADAMTSL1:chr9:18622334 | chr9 | 18622334 | 18622334 | G | A | p.R189Q | Missense_Mutation | nonsilent | SNV | ADAMTSL1:chr9:18622334 |
| 10 | GLRA1:chr5:151851604 | chr5 | 1.52E+08 | 1.52E+08 | C | G | p.G233A | Splice_Site | null | SNV | GLRA1:chr5:151851604 |
| 11 | ABCC8:chr11:17413399 | chr11 | 17413399 | 17413399 | C | T | p.E824K | Missense_Mutation | nonsilent | SNV | ABCC8:chr11:17413399 |
| 12 | APC:chr5:112839465 | chr5 | 1.13E+08 | 1.13E+08 | C | T | p.Q1291* | Nonsense_Mutation | null | SNV | APC:chr5:112839465 |
| 13 | IRAK1:chrX:154019788 | chrX | 1.54E+08 | 1.54E+08 | C | T | p.E9K | Missense_Mutation | nonsilent | SNV | IRAK1:chrX:154019788 |
| 14 | BSN:chr3:49651075 | chr3 | 49651075 | 49651075 | C | A | p.A661E | Missense_Mutation | nonsilent | SNV | BSN:chr3:49651075 |
| 15 | DGAT2:chr11:75798366 | chr11 | 75798366 | 75798366 | C | T | p.R317* | Nonsense_Mutation | null | SNV | DGAT2:chr11:75798366 |
| 16 | ALOX5:chr10:45443465 | chr10 | 45443465 | 45443465 | G | C | p.E501Q | Missense_Mutation | nonsilent | SNV | ALOX5:chr10:45443465 |
| 17 | PCDH9:chr13:66304893 | chr13 | 66304893 | 66304893 | G | A | p.S1159L | Missense_Mutation | nonsilent | SNV | PCDH9:chr13:66304893 |
| 18 | FPR1:chr19:51746492 | chr19 | 51746492 | 51746492 | G | A | p.P168L | Missense_Mutation | nonsilent | SNV | FPR1:chr19:51746492 |
| 19 | C10orf71:chr10:49324658 | chr10 | 49324658 | 49324658 | G | A | p.E705K | Missense_Mutation | nonsilent | SNV | C10orf71:chr10:49324658 |
| 20 | CDKN1A:chr6:36684429 | chr6 | 36684429 | 36684430 | - | A | p.H110fs | Frame_Shift_Ins | null | Insertion | CDKN1A:chr6:36684429 |
| 21 | ADAMTS17:chr15:99997486 | chr15 | 99997486 | 99997486 | C | T | p.A899T | Missense_Mutation | nonsilent | SNV | ADAMTS17:chr15:99997486 |
| 22 | USP28:chr11:113823688 | chr11 | 1.14E+08 | 1.14E+08 | C | G | p.R400S | Missense_Mutation | nonsilent | SNV | USP28:chr11:113823688 |
| 23 | MUC5B:chr11:1248411 | chr11 | 1248411 | 1248411 | C | A | p.T3844N | Missense_Mutation | nonsilent | SNV | MUC5B:chr11:1248411 |
| 24 | PRRC2A:chr6:31632385 | chr6 | 31632385 | 31632385 | G | C | p.E1238Q | Missense_Mutation | nonsilent | SNV | PRRC2A:chr6:31632385 |
| 25 | C5orf42:chr5:37226785 | chr5 | 37226785 | 37226785 | G | A | p.H604Y | Missense_Mutation | nonsilent | SNV | C5orf42:chr5:37226785 |
| 26 | FIGN:chr2:163610118 | chr2 | 1.64E+08 | 1.64E+08 | G | T | p.L572I | Missense_Mutation | nonsilent | SNV | FIGN:chr2:163610118 |
| 27 | UGGT1:chr2:128113209 | chr2 | 1.28E+08 | 1.28E+08 | C | G | p.S216* | Nonsense_Mutation | null | SNV | UGGT1:chr2:128113209 |
| 28 | ITGAV:chr2:186625495 | chr2 | 1.87E+08 | 1.87E+08 | G | A | p.W144* | Nonsense_Mutation | null | SNV | ITGAV:chr2:186625495 |
| 29 | CROCC:chr1:16966080 | chr1 | 16966080 | 16966080 | G | A | p.A1553T | Missense_Mutation | nonsilent | SNV | CROCC:chr1:16966080 |
| 30 | XPO7:chr8:22003299 | chr8 | 22003299 | 22003299 | G | C | p.L1008F | Missense_Mutation | nonsilent | SNV | XPO7:chr8:22003299 |
| 31 | ZCCHC11:chr1:52481843 | chr1 | 52481843 | 52481843 | C | G | p.Q532H | Missense_Mutation | nonsilent | SNV | ZCCHC11:chr1:52481843 |
| 32 | PLA2G4D:chr15:42070059 | chr15 | 42070059 | 42070059 | C | T | p.G694R | Missense_Mutation | nonsilent | SNV | PLA2G4D:chr15:42070059 |
| 33 | SCAPER:chr15:76348651 | chr15 | 76348651 | 76348651 | C | G | p.L1395F | Missense_Mutation | nonsilent | SNV | SCAPER:chr15:76348651 |
| 34 | HERC1:chr15:63666389 | chr15 | 63666389 | 63666389 | G | C | p.L2764V | Missense_Mutation | nonsilent | SNV | HERC1:chr15:63666389 |
| 35 | ADAM12:chr10:126049362 | chr10 | 1.26E+08 | 1.26E+08 | G | A | p.P606L | Missense_Mutation | nonsilent | SNV | ADAM12:chr10:126049362 |
| 36 | MAS1:chr6:159907088 | chr6 | 1.6E+08 | 1.6E+08 | G | A | p.V45M | Missense_Mutation | nonsilent | SNV | MAS1:chr6:159907088 |
| 37 | GPR63:chr6:96798634 | chr6 | 96798634 | 96798634 | C | G | p.L366F | Missense_Mutation | nonsilent | SNV | GPR63:chr6:96798634 |
| 38 | ENPEP:chr4:110476761 | chr4 | 1.1E+08 | 1.1E+08 | C | T | p.T116M | Missense_Mutation | nonsilent | SNV | ENPEP:chr4:110476761 |
| 39 | FBXW7:chr4:152337818 | chr4 | 1.52E+08 | 1.52E+08 | G | C | p.S202* | Nonsense_Mutation | null | SNV | FBXW7:chr4:152337818 |
| 40 | PPP1R12A:chr12:79795669 | chr12 | 79795669 | 79795669 | C | T | p.R851K | Missense_Mutation | nonsilent | SNV | PPP1R12A:chr12:79795669 |
| 41 | GLTSCR2:chr19:47755416 | chr19 | 47755416 | 47755416 | C | G | p.I374M | Missense_Mutation | nonsilent | SNV | GLTSCR2:chr19:47755416 |
| 42 | SCARF1:chr17:1643670 | chr17 | 1643670 | 1643670 | C | T | p.R188H | Missense_Mutation | nonsilent | SNV | SCARF1:chr17:1643670 |
| 43 | SUPV3L1:chr10:69189317 | chr10 | 69189317 | 69189317 | C | T | p.P208L | Missense_Mutation | nonsilent | SNV | SUPV3L1:chr10:69189317 |
| 44 | FAN1:chr15:30905493 | chr15 | 30905493 | 30905493 | C | G | p.S277C | Missense_Mutation | nonsilent | SNV | FAN1:chr15:30905493 |
| 45 | TBC1D32:chr6:121131690 | chr6 | 1.21E+08 | 1.21E+08 | C | G | p.E987Q | Missense_Mutation | nonsilent | SNV | TBC1D32:chr6:121131690 |

**Extended Data Table 5. List of Mutations in Cluster Beta in the Luminal Map (Map 24)**

| | Mutation | Chr | Start | End | Ref | Alt | Protein_Change | Variant_Classification | Effect | Mut_Type | Mut_ID |
|---|---|---|---|---|---|---|---|---|---|---|---|
| 46 | GLB1L3:chr11:134277852 | chr11 | 1.34E+08 | 1.34E+08 | G | A | p.R101K | Missense_Mutation | nonsilent | SNV | GLB1L3:chr11:134277852 |
| 47 | CST11:chr20:23452691 | chr20 | 23452691 | 23452691 | C | G | p.E41Q | Missense_Mutation | nonsilent | SNV | CST11:chr20:23452691 |
| 48 | ZSWIM1:chr20:45884033 | chr20 | 45884033 | 45884033 | G | C | p.E481Q | Missense_Mutation | nonsilent | SNV | ZSWIM1:chr20:45884033 |
| 49 | RBBP8NL:chr20:62414348 | chr20 | 62414348 | 62414348 | C | G | p.E335Q | Missense_Mutation | nonsilent | SNV | RBBP8NL:chr20:62414348 |
| 50 | BRAF:chr7:140781611 | chr7 | 1.41E+08 | 1.41E+08 | C | T | p.G466E | Missense_Mutation | nonsilent | SNV | BRAF:chr7:140781611 |
| 51 | IFT80:chr3:160319905 | chr3 | 1.6E+08 | 1.6E+08 | C | T | p.S271N | Missense_Mutation | nonsilent | SNV | IFT80:chr3:160319905 |
| 52 | MUC16:chr19:8937521 | chr19 | 8937521 | 8937521 | G | T | p.S11145* | Nonsense_Mutation | null | SNV | MUC16:chr19:8937521 |
| 53 | PLCE1:chr10:94313273 | chr10 | 94313273 | 94313273 | G | C | p.R2008T | Missense_Mutation | nonsilent | SNV | PLCE1:chr10:94313273 |
| 54 | ARPC4:chr3:9803843 | chr3 | 9803843 | 9803843 | G | A | p.G111R | Splice_Site | null | SNV | ARPC4:chr3:9803843 |
| 55 | LRP1:chr12:57192941 | chr12 | 57192941 | 57192941 | G | A | p.R2509Q | Missense_Mutation | nonsilent | SNV | LRP1:chr12:57192941 |
| 56 | OGG1:chr3:9751889 | chr3 | 9751889 | 9751889 | C | T | p.R169W | Missense_Mutation | nonsilent | SNV | OGG1:chr3:9751889 |
| 57 | PRKD3:chr2:37316467 | chr2 | 37316467 | 37316467 | G | C | p.P20A | Missense_Mutation | nonsilent | SNV | PRKD3:chr2:37316467 |
| 58 | PRKD3:chr2:37316242 | chr2 | 37316242 | 37316242 | G | C | p.Q95E | Missense_Mutation | nonsilent | SNV | PRKD3:chr2:37316242 |
| 59 | PLEKHM1:chr17:45458393 | chr17 | 45458393 | 45458393 | C | A | p.R452L | Missense_Mutation | nonsilent | SNV | PLEKHM1:chr17:45458393 |
| 60 | RCC1:chr1:28532243 | chr1 | 28532243 | 28532243 | C | T | p.P112S | Missense_Mutation | nonsilent | SNV | RCC1:chr1:28532243 |
| 61 | NELL2:chr12:44779966 | chr12 | 44779966 | 44779966 | G | A | p.S131L | Missense_Mutation | nonsilent | SNV | NELL2:chr12:44779966 |
| 62 | DYRK4:chr12:4604947 | chr12 | 4604947 | 4604947 | C | T | p.S272F | Missense_Mutation | nonsilent | SNV | DYRK4:chr12:4604947 |
| 63 | ATP2B1:chr12:89655717 | chr12 | 89655717 | 89655717 | T | C | p.Y57C | Missense_Mutation | nonsilent | SNV | ATP2B1:chr12:89655717 |
| 64 | KIF2B:chr17:53823520 | chr17 | 53823520 | 53823520 | C | T | p.R163C | Missense_Mutation | nonsilent | SNV | KIF2B:chr17:53823520 |
| 65 | ADGRL3:chr4:62063565 | chr4 | 62063565 | 62063565 | C | A | p.P1292H | Missense_Mutation | nonsilent | SNV | ADGRL3:chr4:62063565 |
| 66 | ZFYVE9:chr1:52266657 | chr1 | 52266657 | 52266657 | C | G | p.Q761E | Missense_Mutation | nonsilent | SNV | ZFYVE9:chr1:52266657 |
| 67 | TRIM62:chr1:33147393 | chr1 | 33147393 | 33147393 | C | T | p.W283* | Nonsense_Mutation | null | SNV | TRIM62:chr1:33147393 |
| 68 | TMPRSS11B:chr4:68234565 | chr4 | 68234565 | 68234565 | C | T | p.E123K | Missense_Mutation | nonsilent | SNV | TMPRSS11B:chr4:68234565 |
| 69 | TIGD7:chr16:3299119 | chr16 | 3299119 | 3299119 | C | G | p.R499T | Missense_Mutation | nonsilent | SNV | TIGD7:chr16:3299119 |
| 70 | DLX4:chr17:49973080 | chr17 | 49973080 | 49973080 | G | C | p.E97D | Missense_Mutation | nonsilent | SNV | DLX4:chr17:49973080 |
| 71 | CCDC17:chr1:45623902 | chr1 | 45623902 | 45623902 | G | A | p.S3F | Missense_Mutation | nonsilent | SNV | CCDC17:chr1:45623902 |
| 72 | CLUH:chr17:2703463 | chr17 | 2703463 | 2703463 | C | G | p.E72D | Missense_Mutation | nonsilent | SNV | CLUH:chr17:2703463 |
| 73 | HS3ST6:chr16:1911844 | chr16 | 1911844 | 1911844 | C | A | p.E259* | Nonsense_Mutation | null | SNV | HS3ST6:chr16:1911844 |
| 74 | ANHX:chr12:133231629 | chr12 | 1.33E+08 | 1.33E+08 | C | T | p.G89R | Missense_Mutation | nonsilent | SNV | ANHX:chr12:133231629 |
| 75 | ATP8B3:chr19:1800082 | chr19 | 1800082 | 1800082 | C | T | p.V473M | Missense_Mutation | nonsilent | SNV | ATP8B3:chr19:1800082 |
| 76 | CYP2C19:chr10:94842987 | chr10 | 94842987 | 94842987 | C | T | p.T371I | Missense_Mutation | nonsilent | SNV | CYP2C19:chr10:94842987 |
| 77 | MFAP1:chr15:43812994 | chr15 | 43812994 | 43812994 | G | C | p.R294G | Missense_Mutation | nonsilent | SNV | MFAP1:chr15:43812994 |
