## Extended Data Table 6 for "The Origin of Bladder Cancer from Mucosal Field Effects"

**Extended Data Table 6. List of Mutations in Cluster Beta in the Basal Map (Map19 )**

| | Mutation | Chr | Start | End | Ref | Alt | hange | Variant_Classification | Effect | Mut_Type | Mut_ID |
|---|---|---|---|---|---|---|---|---|---|---|---|
| 1 | TRIM2:chr4:153322817 | chr4 | 153322817 | 153322817 | G | A | NA | Splice_Site | null | SNV | TRIM2:chr4:153322817 |
| 2 | CHMP6:chr17:80997062 | chr17 | 80997062 | 80997062 | A | G | p.E135G | Missense_Mutation | nonsilent | SNV | CHMP6:chr17:80997062 |
| 3 | CARD19:chr9:93112274 | chr9 | 93112274 | 93112274 | T | C | p.Y141H | Missense_Mutation | nonsilent | SNV | CARD19:chr9:93112274 |
| 4 | KDM6A:chrX:45063557 | chrX | 45063557 | 45063557 | C | T | p.Q555* | Nonsense_Mutation | null | SNV | KDM6A:chrX:45063557 |
| 5 | CHL1:chr3:325986 | chr3 | 325986 | 325986 | A | T | p.Q40L | Missense_Mutation | nonsilent | SNV | CHL1:chr3:325986 |
| 6 | TMEM57:chr1:25484235 | chr1 | 25484235 | 25484235 | T | G | p.M425R | Missense_Mutation | nonsilent | SNV | TMEM57:chr1:25484235 |
| 7 | FREM2:chr13:38689002 | chr13 | 38689002 | 38689002 | T | C | p.L553S | Missense_Mutation | nonsilent | SNV | FREM2:chr13:38689002 |
| 8 | TIGD5:chr8:143598291 | chr8 | 143598291 | 143598291 | G | A | p.A130T | Missense_Mutation | nonsilent | SNV | TIGD5:chr8:143598291 |
| 9 | ITPK1:chr14:92946454 | chr14 | 92946454 | 92946454 | C | T | p.E260K | Missense_Mutation | nonsilent | SNV | ITPK1:chr14:92946454 |
| 10 | C3orf67:chr3:58870294 | chr3 | 58870294 | 58870294 | C | T | p.R39K | Missense_Mutation | nonsilent | SNV | C3orf67:chr3:58870294 |
| 11 | COL7A1:chr3:48581322 | chr3 | 48581322 | 48581322 | C | T | p.G1613R | Missense_Mutation | nonsilent | SNV | COL7A1:chr3:48581322 |
| 12 | DLC1:chr8:13499965 | chr8 | 13499965 | 13499965 | A | G | p.V36A | Missense_Mutation | nonsilent | SNV | DLC1:chr8:13499965 |
| 13 | KIAA2026:chr9:6007547 | chr9 | 6007547 | 6007547 | G | A | p.Q81* | Nonsense_Mutation | null | SNV | KIAA2026:chr9:6007547 |
| 14 | RINT1:chr7:105548609 | chr7 | 105548609 | 105548609 | T | A | p.S299T | Missense_Mutation | nonsilent | SNV | RINT1:chr7:105548609 |
| 15 | LRRN1:chr3:3844711 | chr3 | 3844711 | 3844711 | G | A | p.E24K | Missense_Mutation | nonsilent | SNV | LRRN1:chr3:3844711 |
| 16 | TNFRSF10A:chr8:23199406 | chr8 | 23199406 | 23199406 | C | T | p.E292K | Missense_Mutation | nonsilent | SNV | TNFRSF10A:chr8:23199406 |
| 17 | PHF6:chrX:134377756 | chrX | 134377756 | 134377756 | G | A | NA | Splice_Site | null | SNV | PHF6:chrX:134377756 |
| 18 | ZFHX4:chr8:76705883 | chr8 | 76705883 | 76705883 | G | T | p.G599C | Missense_Mutation | nonsilent | SNV | ZFHX4:chr8:76705883 |
| 19 | TTBK1:chr6:43269662 | chr6 | 43269662 | 43269662 | C | T | p.S618L | Missense_Mutation | nonsilent | SNV | TTBK1:chr6:43269662 |
| 20 | PRSS56:chr2:232524203 | chr2 | 232524203 | 232524203 | G | C | p.G452R | Splice_Site | null | SNV | PRSS56:chr2:232524203 |
| 21 | KIF6:chr6:39540022 | chr6 | 39540022 | 39540022 | A | T | p.S542R | Missense_Mutation | nonsilent | SNV | KIF6:chr6:39540022 |
| 22 | ZNF185:chrX:152959878 | chrX | 152959878 | 152959878 | G | T | p.R529L | Missense_Mutation | nonsilent | SNV | ZNF185:chrX:152959878 |
| 23 | STAM2:chr2:152123931 | chr2 | 152123931 | 152123931 | T | C | p.Y395C | Missense_Mutation | nonsilent | SNV | STAM2:chr2:152123931 |
| 24 | GNE:chr9:36276899 | chr9 | 36276899 | 36276899 | G | A | p.P16S | Missense_Mutation | nonsilent | SNV | GNE:chr9:36276899 |
| 25 | TLR4:chr9:117708721 | chr9 | 117708721 | 117708739 | TTTATCCAGGTAATGAATC | - | NA | Splice_Site | null | Deletion | TLR4:chr9:117708721 |
| 26 | CCDC78:chr16:725278 | chr16 | 725278 | 725278 | T | C | p.T151A | Missense_Mutation | nonsilent | SNV | CCDC78:chr16:725278 |
| 27 | PLEKHH2:chr2:43762375 | chr2 | 43762375 | 43762375 | A | C | p.L1381F | Missense_Mutation | nonsilent | SNV | PLEKHH2:chr2:43762375 |
| 28 | BIRC6:chr2:32513002 | chr2 | 32513002 | 32513002 | G | T | p.M3472I | Missense_Mutation | nonsilent | SNV | BIRC6:chr2:32513002 |
| 29 | EML6:chr2:54813248 | chr2 | 54813248 | 54813248 | G | T | p.D72Y | Missense_Mutation | nonsilent | SNV | EML6:chr2:54813248 |
| 30 | KCNJ3:chr2:154698788 | chr2 | 154698788 | 154698788 | C | T | p.R5* | Nonsense_Mutation | null | SNV | KCNJ3:chr2:154698788 |
| 31 | BIRC6:chr2:32401382 | chr2 | 32401382 | 32401382 | G | A | p.M418I | Missense_Mutation | nonsilent | SNV | BIRC6:chr2:32401382 |
| 32 | ACOT9:chrX:23730544 | chrX | 23730544 | 23730544 | T | A | p.D119V | Missense_Mutation | nonsilent | SNV | ACOT9:chrX:23730544 |
| 33 | DST:chr6:56606403 | chr6 | 56606403 | 56606403 | G | A | p.T2531I | Missense_Mutation | nonsilent | SNV | DST:chr6:56606403 |
| 34 | ANKRD18B:chr9:33529061 | chr9 | 33529061 | 33529061 | A | G | p.N128S | Missense_Mutation | nonsilent | SNV | ANKRD18B:chr9:33529061 |
| 35 | NBDY:chrX:56729484 | chrX | 56729484 | 56729484 | G | A | p.G44D | Missense_Mutation | nonsilent | SNV | NBDY:chrX:56729484 |
| 36 | OR2J3:chr6:29112374 | chr6 | 29112374 | 29112374 | C | A | p.H162N | Missense_Mutation | nonsilent | SNV | OR2J3:chr6:29112374 |
| 37 | RAPH1:chr2:203495305 | chr2 | 203495305 | 203495305 | T | G | p.S17R | Missense_Mutation | nonsilent | SNV | RAPH1:chr2:203495305 |
| 38 | MID2:chrX:107926111 | chrX | 107926111 | 107926111 | G | A | p.D509N | Missense_Mutation | nonsilent | SNV | MID2:chrX:107926111 |
| 39 | GPR1:chr2:206176749 | chr2 | 206176749 | 206176749 | G | C | p.L167V | Missense_Mutation | nonsilent | SNV | GPR1:chr2:206176749 |
| 40 | CDKN2A:chr9:21971210 | chr9 | 21971210 | 21971210 | T | A | NA | Splice_Site | null | SNV | CDKN2A:chr9:21971210 |
| 41 | PDE11A:chr2:178071601 | chr2 | 178071601 | 178071601 | C | A | p.Q279H | Missense_Mutation | nonsilent | SNV | PDE11A:chr2:178071601 |
| 42 | NUMA1:chr11:72013472 | chr11 | 72013472 | 72013472 | G | A | p.S1344F | Missense_Mutation | nonsilent | SNV | NUMA1:chr11:72013472 |
| 43 | CAPRIN1:chr11:34089401 | chr11 | 34089401 | 34089401 | C | G | p.S413C | Missense_Mutation | nonsilent | SNV | CAPRIN1:chr11:34089401 |
| 44 | MLLT3:chr9:20414345 | chr9 | 20414345 | 20414346 | - | CTG | .167_168ins | In_Frame_Ins | null | Insertion | MLLT3:chr9:20414345 |
| 45 | SLC7A2:chr8:17538915 | chr8 | 17538915 | 17538915 | T | G | p.F31V | Missense_Mutation | nonsilent | SNV | SLC7A2:chr8:17538915 |
| 46 | NR1D2:chr3:23977364 | chr3 | 23977364 | 23977364 | G | T | p.R562L | Missense_Mutation | nonsilent | SNV | NR1D2:chr3:23977364 |
| 47 | BAZ2B:chr2:159389475 | chr2 | 159389475 | 159389475 | C | A | p.R1029L | Missense_Mutation | nonsilent | SNV | BAZ2B:chr2:159389475 |
| 48 | PRKCB:chr16:24214724 | chr16 | 24214724 | 24214724 | G | A | p.D644N | Missense_Mutation | nonsilent | SNV | PRKCB:chr16:24214724 |
| 49 | MAFA:chr8:143430396 | chr8 | 143430396 | 143430396 | T | C | p.E4G | Missense_Mutation | nonsilent | SNV | MAFA:chr8:143430396 |
| 50 | SHANK1:chr19:50666505 | chr19 | 50666505 | 50666505 | C | G | p.E1819Q | Missense_Mutation | nonsilent | SNV | SHANK1:chr19:50666505 |

**Extended Data Table 6. List of Mutations in Cluster Beta in the Basal Map (Map19 )**

| | Mutation | Chr | Start | End | Ref | Alt | hange | Variant_Classification | Effect | Mut_Type | Mut_ID |
|---|---|---|---|---|---|---|---|---|---|---|---|
| 51 | FCGR1A:chr1:149791248 | chr1 | 149791248 | 149791248 | C | T | p.P286S | Missense_Mutation | nonsilent | SNV | FCGR1A:chr1:149791248 |
| 52 | PDLIM2:chr8:22585371 | chr8 | 22585371 | 22585371 | C | G | p.S421C | Missense_Mutation | nonsilent | SNV | PDLIM2:chr8:22585371 |
| 53 | HSPG2:chr1:21829004 | chr1 | 21829004 | 21829004 | T | A | p.K4023M | Missense_Mutation | nonsilent | SNV | HSPG2:chr1:21829004 |
| 54 | NBN:chr8:89982786 | chr8 | 89982786 | 89982786 | T | G | p.E36A | Missense_Mutation | nonsilent | SNV | NBN:chr8:89982786 |
| 55 | PKHD1L1:chr8:109448344 | chr8 | 109448344 | 109448344 | T | C | p.V1993A | Missense_Mutation | nonsilent | SNV | PKHD1L1:chr8:109448344 |
| 56 | GNAI1:chr7:80199245 | chr7 | 80199245 | 80199269 | TGCTAGCTGGAGCTGCTGAAG | - | p.F108fs | Frame_Shift_Del | null | Deletion | GNAI1:chr7:80199245 |
| 57 | DOK7:chr4:3473424 | chr4 | 3473424 | 3473424 | T | C | p.V40A | Missense_Mutation | nonsilent | SNV | DOK7:chr4:3473424 |
| 58 | GPR26:chr10:123667006 | chr10 | 123667006 | 123667006 | G | T | p.R200L | Missense_Mutation | nonsilent | SNV | GPR26:chr10:123667006 |
| 59 | RSF1:chr11:77666993 | chr11 | 77666993 | 77666993 | G | C | p.A1417G | Missense_Mutation | nonsilent | SNV | RSF1:chr11:77666993 |
| 60 | USP43:chr17:9646065 | chr17 | 9646065 | 9646065 | G | A | p.E145K | Missense_Mutation | nonsilent | SNV | USP43:chr17:9646065 |
| 61 | LLGL2:chr17:75563386 | chr17 | 75563386 | 75563386 | C | G | p.S250C | Missense_Mutation | nonsilent | SNV | LLGL2:chr17:75563386 |
| 62 | ACSL6:chr5:131974798 | chr5 | 131974798 | 131974798 | G | T | p.S318Y | Missense_Mutation | nonsilent | SNV | ACSL6:chr5:131974798 |
| 63 | CLDN12:chr7:90413086 | chr7 | 90413086 | 90413086 | G | A | p.G137E | Missense_Mutation | nonsilent | SNV | CLDN12:chr7:90413086 |
| 64 | LRRC71:chr1:156924038 | chr1 | 156924038 | 156924038 | C | T | p.R84W | Missense_Mutation | nonsilent | SNV | LRRC71:chr1:156924038 |
| 65 | ST6GALNAC5:chr1:77044267 | chr1 | 77044267 | 77044267 | C | G | p.R109G | Missense_Mutation | nonsilent | SNV | ST6GALNAC5:chr1:77044267 |
| 66 | SRP68:chr17:76072412 | chr17 | 76072412 | 76072412 | C | T | p.S27N | Missense_Mutation | nonsilent | SNV | SRP68:chr17:76072412 |
| 67 | OR2G2:chr1:247589077 | chr1 | 247589077 | 247589077 | G | A | p.A240T | Missense_Mutation | nonsilent | SNV | OR2G2:chr1:247589077 |
| 68 | DHX9:chr1:182876227 | chr1 | 182876227 | 182876227 | A | G | p.T665A | Missense_Mutation | nonsilent | SNV | DHX9:chr1:182876227 |
| 69 | CDK13:chr7:40001877 | chr7 | 40001877 | 40001877 | A | - | p.V736fs | Frame_Shift_Del | null | Deletion | CDK13:chr7:40001877 |
| 70 | SLC35E4:chr22:30646733 | chr22 | 30646733 | 30646733 | C | G | p.S252C | Missense_Mutation | nonsilent | SNV | SLC35E4:chr22:30646733 |
| 71 | HTRA1:chr10:122462125 | chr10 | 122462125 | 122462125 | G | T | NA | Splice_Site | null | SNV | HTRA1:chr10:122462125 |
| 72 | HOOK1:chr1:59843580 | chr1 | 59843580 | 59843580 | T | C | p.L257S | Missense_Mutation | nonsilent | SNV | HOOK1:chr1:59843580 |
| 73 | ATAD5:chr17:30893495 | chr17 | 30893495 | 30893495 | G | - | p.A1548fs | Frame_Shift_Del | null | Deletion | ATAD5:chr17:30893495 |
| 74 | FRYL:chr4:48576220 | chr4 | 48576220 | 48576220 | C | T | p.S844N | Missense_Mutation | nonsilent | SNV | FRYL:chr4:48576220 |
| 75 | NME9:chr3:138306060 | chr3 | 138306060 | 138306060 | C | G | p.E194Q | Missense_Mutation | nonsilent | SNV | NME9:chr3:138306060 |
| 76 | SYT11:chr1:155868387 | chr1 | 155868387 | 155868387 | G | C | p.E153Q | Missense_Mutation | nonsilent | SNV | SYT11:chr1:155868387 |
| 77 | RNPC3:chr1:103544975 | chr1 | 103544975 | 103544975 | A | G | p.I360M | Missense_Mutation | nonsilent | SNV | RNPC3:chr1:103544975 |
| 78 | DCAF5:chr14:69054365 | chr14 | 69054365 | 69054365 | G | T | p.P774H | Missense_Mutation | nonsilent | SNV | DCAF5:chr14:69054365 |
| 79 | CCP110:chr16:19527996 | chr16 | 19527996 | 19527996 | C | T | p.H39Y | Missense_Mutation | nonsilent | SNV | CCP110:chr16:19527996 |
| 80 | BAIAP3:chr16:1345091 | chr16 | 1345091 | 1345091 | C | G | p.I608M | Missense_Mutation | nonsilent | SNV | BAIAP3:chr16:1345091 |
| 81 | PIK3CG:chr7:106867578 | chr7 | 106867578 | 106867578 | A | G | p.Y6C | Missense_Mutation | nonsilent | SNV | PIK3CG:chr7:106867578 |
| 82 | SPTBN5:chr15:41878549 | chr15 | 41878549 | 41878549 | T | A | p.Q1088L | Missense_Mutation | nonsilent | SNV | SPTBN5:chr15:41878549 |
| 83 | EXOC7:chr17:76089303 | chr17 | 76089303 | 76089303 | C | A | p.D276Y | Missense_Mutation | nonsilent | SNV | EXOC7:chr17:76089303 |
| 84 | DLGAP1:chr18:3499240 | chr18 | 3499240 | 3499240 | G | A | p.T668I | Missense_Mutation | nonsilent | SNV | DLGAP1:chr18:3499240 |
| 85 | NT5C2:chr10:103098954 | chr10 | 103098954 | 103098954 | G | T | p.L222I | Missense_Mutation | nonsilent | SNV | NT5C2:chr10:103098954 |
| 86 | BATF:chr14:75522689 | chr14 | 75522689 | 75522689 | C | T | p.H3Y | Missense_Mutation | nonsilent | SNV | BATF:chr14:75522689 |
| 87 | HORMAD1:chr1:150706552 | chr1 | 150706552 | 150706552 | C | A | NA | Splice_Site | null | SNV | HORMAD1:chr1:150706552 |
| 88 | TTLL12:chr22:43174548 | chr22 | 43174548 | 43174548 | C | T | p.E329K | Missense_Mutation | nonsilent | SNV | TTLL12:chr22:43174548 |
| 89 | CCM2L:chr20:32014991 | chr20 | 32014991 | 32014991 | C | T | p.H40Y | Missense_Mutation | nonsilent | SNV | CCM2L:chr20:32014991 |
| 90 | NDUFB9:chr8:124543117 | chr8 | 124543117 | 124543117 | G | A | p.M44I | Missense_Mutation | nonsilent | SNV | NDUFB9:chr8:124543117 |
| 91 | CD22:chr19:35332729 | chr19 | 35332729 | 35332729 | G | A | p.G73R | Missense_Mutation | nonsilent | SNV | CD22:chr19:35332729 |
| 92 | CCT8L2:chr22:16591947 | chr22 | 16591947 | 16591947 | G | T | p.R202S | Missense_Mutation | nonsilent | SNV | CCT8L2:chr22:16591947 |
| 93 | KRT18:chr12:52951733 | chr12 | 52951733 | 52951733 | T | G | p.I275M | Missense_Mutation | nonsilent | SNV | KRT18:chr12:52951733 |
| 94 | PTTG1:chr5:160422706 | chr5 | 160422706 | 160422708 | CAG | - | NA | Splice_Site | null | Deletion | PTTG1:chr5:160422706 |
| 95 | PPP4R4:chr14:94208538 | chr14 | 94208538 | 94208538 | C | T | p.T89M | Missense_Mutation | nonsilent | SNV | PPP4R4:chr14:94208538 |
| 96 | PLOD2:chr3:146079211 | chr3 | 146079211 | 146079211 | C | T | p.G469R | Missense_Mutation | nonsilent | SNV | PLOD2:chr3:146079211 |
| 97 | DLGAP5:chr14:55150819 | chr14 | 55150819 | 55150819 | C | A | p.E800* | Nonsense_Mutation | null | SNV | DLGAP5:chr14:55150819 |
| 98 | ZNF583:chr19:56424155 | chr19 | 56424155 | 56424155 | A | T | p.K499N | Missense_Mutation | nonsilent | SNV | ZNF583:chr19:56424155 |
| 99 | PIK3AP1:chr10:96620477 | chr10 | 96620477 | 96620477 | G | A | p.R606W | Missense_Mutation | nonsilent | SNV | PIK3AP1:chr10:96620477 |
| 100 | SPATA12:chr3:57073911 | chr3 | 57073911 | 57073911 | C | T | p.Q73* | Nonsense_Mutation | null | SNV | SPATA12:chr3:57073911 |

**Extended Data Table 6. List of Mutations in Cluster Beta in the Basal Map (Map19 )**

| | Mutation | Chr | Start | End | Ref | Alt | hange | Variant_Classification | Effect | Mut_Type | Mut_ID |
|---|---|---|---|---|---|---|---|---|---|---|---|
| 101 | HMHA1:chr19:1073268 | chr19 | 1073268 | 1073268 | G | T | p.G197C | Missense_Mutation | nonsilent | SNV | HMHA1:chr19:1073268 |
| 102 | MUM1:chr19:1370737 | chr19 | 1370737 | 1370737 | C | T | p.P550S | Missense_Mutation | nonsilent | SNV | MUM1:chr19:1370737 |
| 103 | HSPG2:chr1:21842831 | chr1 | 21842831 | 21842831 | C | A | p.G2950V | Missense_Mutation | nonsilent | SNV | HSPG2:chr1:21842831 |
| 104 | TRMT2A:chr22:20116153 | chr22 | 20116153 | 20116153 | C | T | p.E162K | Missense_Mutation | nonsilent | SNV | TRMT2A:chr22:20116153 |
| 105 | CAMSAP3:chr19:7611931 | chr19 | 7611931 | 7611931 | G | A | p.D480N | Missense_Mutation | nonsilent | SNV | CAMSAP3:chr19:7611931 |
| 106 | NANOS3:chr19:13877615 | chr19 | 13877615 | 13877615 | C | T | p.H123Y | Missense_Mutation | nonsilent | SNV | NANOS3:chr19:13877615 |
| 107 | USP41:chr22:20380802 | chr22 | 20380802 | 20380802 | T | A | p.Q44L | Missense_Mutation | nonsilent | SNV | USP41:chr22:20380802 |
| 108 | SMARCA4:chr19:10984249 | chr19 | 10984249 | 10984249 | G | A | p.G33D | Missense_Mutation | nonsilent | SNV | SMARCA4:chr19:10984249 |
| 109 | SGSM1:chr22:24886624 | chr22 | 24886624 | 24886624 | C | T | p.R611C | Missense_Mutation | nonsilent | SNV | SGSM1:chr22:24886624 |
| 110 | SLC12A1:chr15:48267587 | chr15 | 48267587 | 48267587 | G | T | p.E727D | Missense_Mutation | nonsilent | SNV | SLC12A1:chr15:48267587 |
| 111 | AP1G2:chr14:23565680 | chr14 | 23565680 | 23565680 | T | C | p.I223V | Missense_Mutation | nonsilent | SNV | AP1G2:chr14:23565680 |
| 112 | LIN7C:chr11:27501562 | chr11 | 27501562 | 27501562 | T | C | p.Y54C | Missense_Mutation | nonsilent | SNV | LIN7C:chr11:27501562 |
| 113 | C1orf167:chr1:11766304 | chr1 | 11766304 | 11766304 | C | G | p.A363G | Missense_Mutation | nonsilent | SNV | C1orf167:chr1:11766304 |
| 114 | SRRM2:chr16:2758992 | chr16 | 2758992 | 2758992 | G | A | p.E201K | Missense_Mutation | nonsilent | SNV | SRRM2:chr16:2758992 |
| 115 | MRE11A:chr11:94467865 | chr11 | 94467865 | 94467865 | C | A | p.R349L | Missense_Mutation | nonsilent | SNV | MRE11A:chr11:94467865 |
| 116 | TRIM42:chr3:140678519 | chr3 | 140678519 | 140678519 | G | A | p.R97H | Missense_Mutation | nonsilent | SNV | TRIM42:chr3:140678519 |
| 117 | KMT2C:chr7:152167145 | chr7 | 152167145 | 152167145 | C | T | NA | Splice_Site | null | SNV | KMT2C:chr7:152167145 |
| 118 | TRIM33:chr1:114510971 | chr1 | 114510971 | 114510971 | G | A | p.P36S | Missense_Mutation | nonsilent | SNV | TRIM33:chr1:114510971 |
| 119 | PENK:chr8:56441310 | chr8 | 56441310 | 56441310 | C | T | p.E256K | Missense_Mutation | nonsilent | SNV | PENK:chr8:56441310 |
| 120 | ITPR2:chr12:26658018 | chr12 | 26658018 | 26658018 | G | A | p.Q667* | Nonsense_Mutation | null | SNV | ITPR2:chr12:26658018 |
| 121 | MRPS28:chr8:80030219 | chr8 | 80030219 | 80030234 | CACAGCACGGGTCCGA | - | p.C5fs | Frame_Shift_Del | null | Deletion | MRPS28:chr8:80030219 |
| 122 | PRDM5:chr4:120922526 | chr4 | 120922526 | 120922526 | C | T | p.R28K | Missense_Mutation | nonsilent | SNV | PRDM5:chr4:120922526 |
| 123 | IGFL4:chr19:46040280 | chr19 | 46040280 | 46040280 | C | A | p.W69C | Missense_Mutation | nonsilent | SNV | IGFL4:chr19:46040280 |
| 124 | FERD3L:chr7:19145329 | chr7 | 19145329 | 19145329 | T | C | p.T12A | Missense_Mutation | nonsilent | SNV | FERD3L:chr7:19145329 |
| 125 | RPUSD1:chr16:786923 | chr16 | 786923 | 786923 | C | T | p.E139K | Missense_Mutation | nonsilent | SNV | RPUSD1:chr16:786923 |
| 126 | AFF4:chr5:132888116 | chr5 | 132888116 | 132888116 | T | A | p.K926M | Missense_Mutation | nonsilent | SNV | AFF4:chr5:132888116 |
| 127 | DSEL:chr18:67511308 | chr18 | 67511308 | 67511308 | G | A | p.H1111Y | Missense_Mutation | nonsilent | SNV | DSEL:chr18:67511308 |
| 128 | APH1B:chr15:63287541 | chr15 | 63287541 | 63287541 | A | G | p.Y158C | Missense_Mutation | nonsilent | SNV | APH1B:chr15:63287541 |
| 129 | TLR5:chr1:223111903 | chr1 | 223111903 | 223111903 | C | T | p.D377N | Missense_Mutation | nonsilent | SNV | TLR5:chr1:223111903 |
| 130 | SFPQ:chr1:35192344 | chr1 | 35192344 | 35192344 | G | A | p.R236* | Nonsense_Mutation | null | SNV | SFPQ:chr1:35192344 |
| 131 | SUCO:chr1:172577765 | chr1 | 172577765 | 172577765 | T | - | p.E430fs | Splice_Site | null | Deletion | SUCO:chr1:172577765 |
| 132 | GMEB2:chr20:63597784 | chr20 | 63597784 | 63597784 | G | A | p.A145V | Missense_Mutation | nonsilent | SNV | GMEB2:chr20:63597784 |
| 133 | UBR4:chr1:19121222 | chr1 | 19121222 | 19121222 | T | C | p.S3370G | Missense_Mutation | nonsilent | SNV | UBR4:chr1:19121222 |
| 134 | SPINK9:chr5:148323790 | chr5 | 148323790 | 148323790 | A | G | p.K14E | Missense_Mutation | nonsilent | SNV | SPINK9:chr5:148323790 |
| 135 | SPAG9:chr17:51120429 | chr17 | 51120429 | 51120429 | C | A | p.E76D | Missense_Mutation | nonsilent | SNV | SPAG9:chr17:51120429 |
| 136 | ZMYM6:chr1:34988419 | chr1 | 34988419 | 34988419 | T | A | p.D888V | Missense_Mutation | nonsilent | SNV | ZMYM6:chr1:34988419 |
| 137 | PPP1R12A:chr12:79808570 | chr12 | 79808570 | 79808570 | T | C | p.D488G | Missense_Mutation | nonsilent | SNV | PPP1R12A:chr12:79808570 |
| 138 | FBXW7:chr4:152352482 | chr4 | 152352482 | 152352482 | C | T | p.M83I | Missense_Mutation | nonsilent | SNV | FBXW7:chr4:152352482 |
| 139 | CNST:chr1:246647169 | chr1 | 246647169 | 246647169 | G | T | p.S323I | Missense_Mutation | nonsilent | SNV | CNST:chr1:246647169 |
| 140 | HSPA4L:chr4:127830689 | chr4 | 127830689 | 127830689 | T | C | p.C740R | Missense_Mutation | nonsilent | SNV | HSPA4L:chr4:127830689 |
| 141 | RPS6KA1:chr1:26558903 | chr1 | 26558903 | 26558903 | G | T | p.R394L | Missense_Mutation | nonsilent | SNV | RPS6KA1:chr1:26558903 |
| 142 | NDN:chr15:23686552 | chr15 | 23686552 | 23686552 | C | T | p.W222* | Nonsense_Mutation | null | SNV | NDN:chr15:23686552 |
| 143 | MROH9:chr1:171024763 | chr1 | 171024763 | 171024763 | C | A | p.L726I | Missense_Mutation | nonsilent | SNV | MROH9:chr1:171024763 |
| 144 | ANKIB1:chr7:92398425 | chr7 | 92398425 | 92398425 | T | G | p.L916V | Missense_Mutation | nonsilent | SNV | ANKIB1:chr7:92398425 |
| 145 | FAT2:chr5:151551528 | chr5 | 151551528 | 151551528 | C | T | p.R1412K | Missense_Mutation | nonsilent | SNV | FAT2:chr5:151551528 |
| 146 | TRIM67:chr1:231206787 | chr1 | 231206787 | 231206787 | G | A | p.D606N | Missense_Mutation | nonsilent | SNV | TRIM67:chr1:231206787 |
| 147 | COQ9:chr16:57458324 | chr16 | 57458324 | 57458324 | C | G | p.H194D | Missense_Mutation | nonsilent | SNV | COQ9:chr16:57458324 |
| 148 | ATP2C2:chr16:84410749 | chr16 | 84410749 | 84410749 | C | T | p.P167S | Missense_Mutation | nonsilent | SNV | ATP2C2:chr16:84410749 |
| 149 | ANKRD27:chr19:32615668 | chr19 | 32615668 | 32615668 | G | A | p.P722L | Missense_Mutation | nonsilent | SNV | ANKRD27:chr19:32615668 |
| 150 | MUC15:chr11:26565425 | chr11 | 26565425 | 26565425 | G | A | p.S145F | Missense_Mutation | nonsilent | SNV | MUC15:chr11:26565425 |

**Extended Data Table 6. List of Mutations in Cluster Beta in the Basal Map (Map19 )**

| | Mutation | Chr | Start | End | Ref | Alt | hange | Variant_Classification | Effect | Mut_Type | Mut_ID |
|---|---|---|---|---|---|---|---|---|---|---|---|
| 151 | NASP:chr1:45606522 | chr1 | 45606522 | 45606522 | C | A | p.H114N | Missense_Mutation | nonsilent | SNV | NASP:chr1:45606522 |
| 152 | KIAA0825:chr5:94396405 | chr5 | 94396405 | 94396405 | C | T | p.E998K | Missense_Mutation | nonsilent | SNV | KIAA0825:chr5:94396405 |
| 153 | MGEA5:chr10:101799147 | chr10 | 101799147 | 101799147 | C | T | p.D502N | Missense_Mutation | nonsilent | SNV | MGEA5:chr10:101799147 |
| 154 | CAP1:chr1:40059455 | chr1 | 40059455 | 40059455 | A | C | p.K37Q | Missense_Mutation | nonsilent | SNV | CAP1:chr1:40059455 |
| 155 | BTG2:chr1:203307254 | chr1 | 203307254 | 203307254 | G | C | p.S98T | Missense_Mutation | nonsilent | SNV | BTG2:chr1:203307254 |
